# The super-pangenome of *Populus* unveil genomic facets for adaptation and diversification in widespread forest trees

**DOI:** 10.1101/2023.07.18.549473

**Authors:** Tingting Shi, Xinxin Zhang, Yukang Hou, Yuanzhong Jiang, Changfu Jia, Qiang Lai, Xuming Dan, Jiajun Feng, Jianju Feng, Tao Ma, Jiali Wu, Shuyu Liu, Lei Zhang, Zhiqin Long, Yulin Zhang, Jiaqi Zhang, Liyang Chen, Nathaniel R. Street, Pär K. Ingvarsson, Jianquan Liu, Tongming Yin, Jing Wang

## Abstract

Understanding the underlying mechanisms between genome evolution and phenotypic and adaptive innovations is a key goal of evolutionary studies. Poplars are the world’s most widely distributed and cultivated trees, with extensive phenotypic diversity and environmental adaptability. Here we report a genus-level super-pangenome of 19 *Populus* genomes. After integrating pan-genomes with transcriptomes, methylomes and chromatin accessibility mapping, we reveal that the evolutionary fate of pan-genes and duplicated genes are largely associated with local genomic landscapes of regulatory and epigenetic architectures. Further comparative genomic analyses enabled to identify 142,202 structural variations (SVs) across species, which overlap with substantial genes and play key roles in both phenotypic and adaptive divergence. We experimentally validated a ∼180 bp presence/absence variant located in the promoter of the *CUC2* gene, which contributed critically to leaf serration divergence between species. Together, this first super-pangenome resource in forest trees will not only accelerate molecular functional studies and genetic breeding of this globally important tree genus, but also lays a foundation for our understanding of tree biology.

## INTRODUCTION

Forests cover approximately 30% of the Earth’s terrestrial surface and provide humanity with clean air, fiber, food and fuel^1^. Large and long-lived forests serve numerous important ecological roles, from providing substantial habitat for terrestrial biodiversity to mitigating the effects of global climate change^2, 3^. Despite their great ecological and economic importance, both genomic and molecular studies of forest trees have lagged behind other model herbaceous plants and crops^4^. Members of the genus *Populus* have been established as model forest trees for diverse research areas because of their relatively small genome sizes and the ease of genetic and experimental manipulation^5^. In addition, poplars are one of the most widely naturally distributed and cultivated trees in the world; they are found throughout the Northern Hemisphere and some tropical regions in the Southern Hemisphere (Fig. 1a). Their diverse habitats vary from arid deserts to wet tropical regions. There are 30 to 80 species in the genus and they are classified into five to eight intrageneric sections depending on different taxonomists^6^. These sections encompass extensive phenotypic diversity, for example, leaf margins range from being entire smooth to deeply serrated^7^ (Fig. 1a). In addition, all poplar species experienced a recent whole genome duplication (WGD) before their species diversification^8, 9^. The subsequent evolution through structural variations and random retention of the duplicated genes may have played an important role in widespread adaptation and the phenotypic diversity of the genus *Populus*. The various species within the genus, therefore, contain rich genetic diversity including allelic variations, private genes and structural variations that are essential for facilitating genetic modifications of the cultivated poplars and for supplementing germplasms for the biotic and abiotic stress tolerance required to adapt to future climate change.

**Fig. 1.**
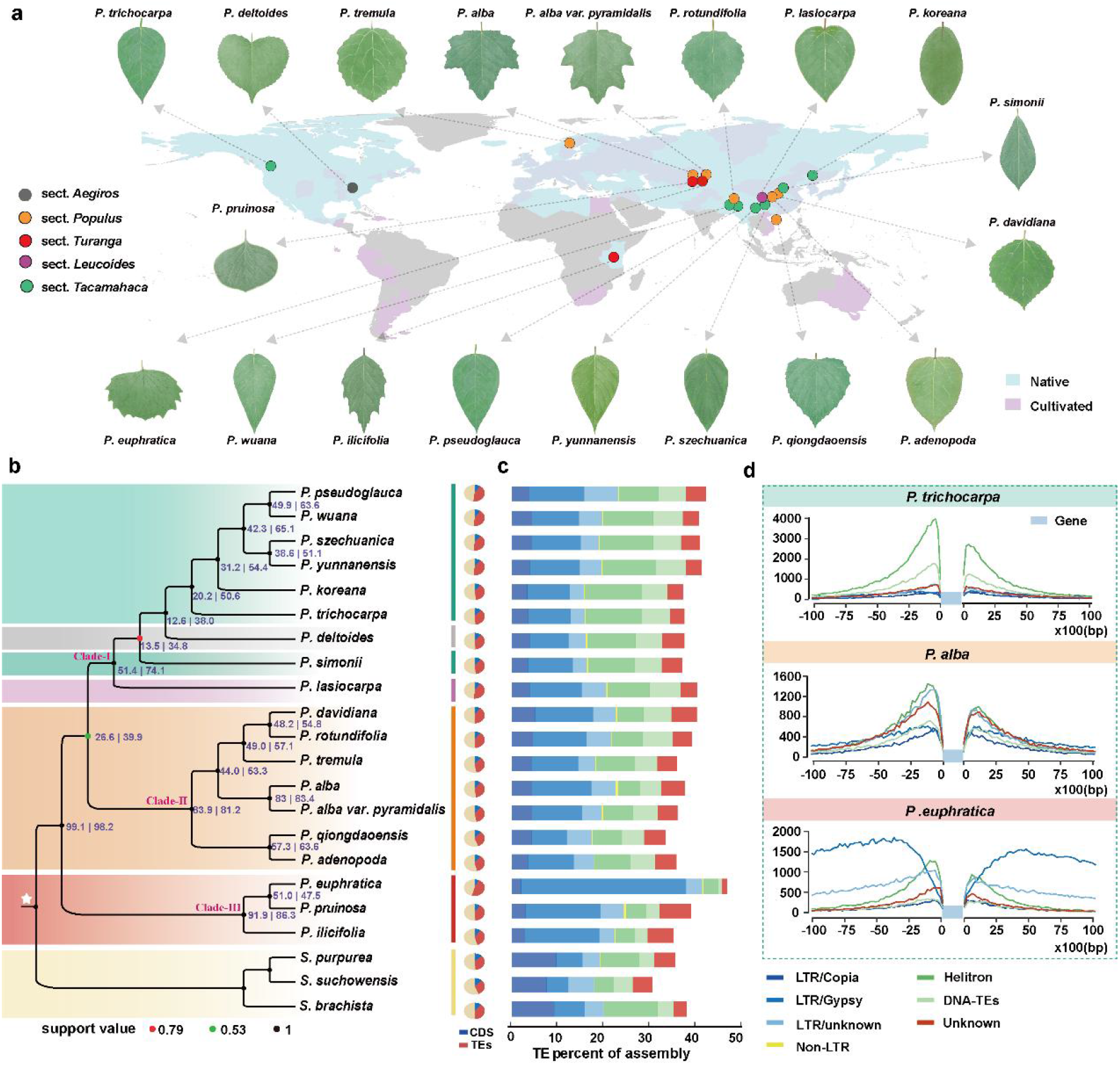
Geographical distribution and genome features of poplar species. **a,** The map of naturally distributed and cultivated poplars in the world, with the colored circles representing the sampling locations of 19 poplar genomes. **b,** From left to right: phylogenetic relationships inferred on the basis of 2,455 single-copy orthologous groups using ASTRAL. Node labels in blue are in the following format: gene concordance factors (gCFs) | site concordance factors (sCFs). For each node in the ASTRAL species tree, gCFs reflect the percentage of gene trees that contain that node as defined by its descendant taxa, and sCFs reflect the percentage of informative sites that support that node via parsimony; fraction of CDS (blue) and TEs (red) in each genome. The five sections of *Populus* and the outgroup *Salix* are shaded by different colors. **c,** Bar plots show the fraction of each TE superfamily (represented by different color as in panel d) in poplar genomes. **d,** TE landscape surrounding genes in three poplars. The 10 kbp upstream of the transcription start site and 10 kbp downstream of the transcription end site across genes were analyzed.

High-quality reference genomes for several species have promoted breeding and functional studies of poplar trees^8, 10, 11^. By resequencing numerous accessions of these and closely related species, single nucleotide polymorphisms (SNPs) and small insertion/deletions (InDels) have been identified to assist in clarifying local adaptation and phenotypic diversification^10, 12, 13^. However, these approaches can only reveal a part of the genetic variation; it is difficult, however, to recover private genes and large structural variants (SVs) at the genus level that may contribute more to genomic evolution and phenotypic diversity^14, 15^. To overcome these limitations, pan-genomic analyses based on multiple assembled *de novo* genomes have been developed to capture the nearly complete spectrum of genetic diversity and reveal the hidden ability to inherit for individual or closely related species^16–18^. In this study, we extended these pan-genomic analyses to the genus level and assembled high-quality chromosome-level genomes of 6 species of the genus *Populus* from different geographical regions. These poplars have different phenotypes and/or evolutionary histories from the currently published poplars, which can be a great complement to poplar genome resources. We further collected previously published genome sequences of 13 additional species/sub-species^8, 10, 11^, with the combined 19 species/sub-species covering major clades of the genus^6^. This panel of genomes was used to perform a super-pangenomic analysis which facilitated the discovery of genomic variations at the genus level, including genes, transposable elements (TEs) and structural variants (SVs). After using complementary Bisulfite-, ATAC- and RNA-sequencing data (Extended Data Fig. 1a), it opens new opportunities to explore the evolutionary causes and consequences of epigenetic and regulatory architectures as species diverged and adapted to a wide range of ecological niches in the genus *Populus.* We particularly aimed to identify SVs and also the hemizygous genes across these representative species and to clarify the functional roles of these SVs in the widespread adaptation and phenotypic diversity of the genus.

## RESULTS

### Chromosome-scale reference genomes of newly sequenced *Populus* species

To characterize the super-pangenome architecture of the genus *Populus*, we selected 19 reference genomes comprising 18 wild species and 1 sub-species from four sections of *Populus*, in which 6 were *de novo* assembled at the chromosome level in this study and the other 13 were published in other studies (Table 1). The 6 new genomes were sequenced using a combination of Illumina short-read sequencing (average coverage depth of 57.1× per genome), Oxford Nanopore long-read sequencing (average coverage depth of 88.5× per genome) and high-throughput chromosome conformation capture (Hi-C) technologies (average sequencing depth of 152.9× per genome) (Supplementary Table 2). Using these sequencing data, the new genomes were assembled with contig N50 sizes ranging from 3.2Mb to 6.3Mb and assembly sizes ranging from 408.0Mb to 448.7Mb after removing redundant sequences and potential contaminated sequences (Table 1 and Supplementary Table 2). Based on Hi-C read pairs, the assembled contigs were further anchored to 19 pseudo-chromosomes with an average anchoring rate of 98.3% across species (Extended Data Fig. 1b and Supplementary Fig. 1). The completeness of all assemblies exceeded 98% when evaluated using Benchmarking Universal Single-Copy Orthologs (BUSCO)^19^ (Extended Data Fig. 1c). Further evaluation using Merqury^20^ showed a quality value (QV) over 30 for all assemblies, which reached the TrioCanu human (NA12878) assembly standard of QV 30^20^ (Supplementary Table 2).

**Table 1.**
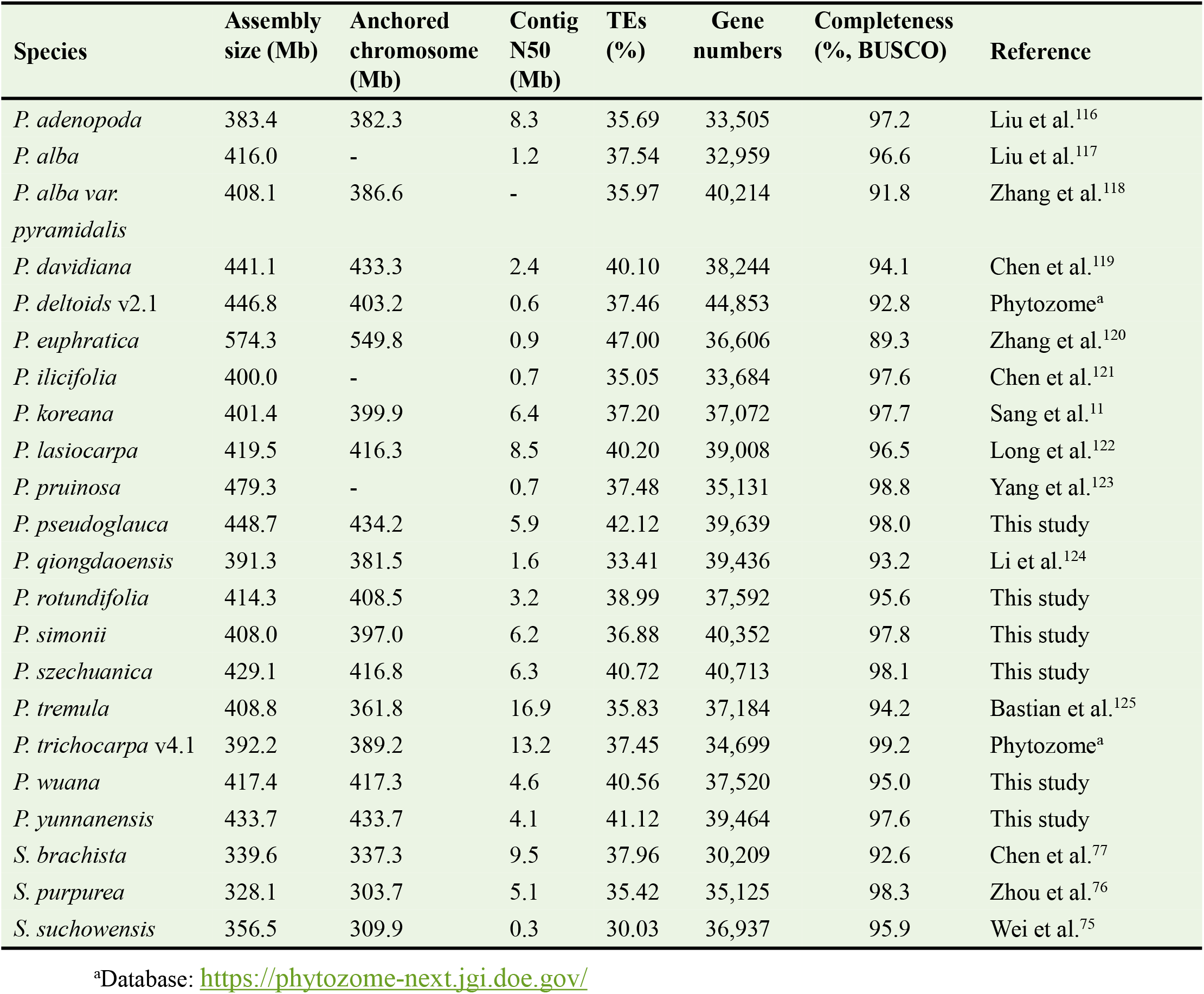
Statistics of the genomic assembly and annotation of 19 *Populus* and 3 *Salix* genomes.

By combining *ab initio*, homology-based and transcriptome-based approaches, 37,520-40,713 protein-coding genes were identified from the newly assembled genomes (Table 1), of which 93.8%-95.3% were functionally annotated through at least one of the databases TrEMBL, Swiss-Prot, NR, Pfam, Interproscan, GO or KEGG (Supplementary Tables 3,4). BUSCO completeness scores of annotated genes from the newly assembled genomes ranged from 95.0 to 98.1% (Table 1 and Supplementary Fig. 2). We observed a high level of chromosomal-scale genetic synteny across the 19 species (Extended Data Fig. 1d and Supplementary Fig. 3), supporting the suggestion that the karyotypes of these species have remained remarkably stable. To minimize methodological artifacts of whole-genome transposable elements (TE) annotation, we used a uniform annotation pipeline for both newly assembled and previously published genomes. We found that repetitive elements made up ∼ 38.5% of genomic sequences with a range from 33.4 to 47.0% across species (Table 1, Fig. 1b and Supplementary Table 5). Variable TE abundance were largely explained in genome size across the species (Supplementary Fig. 4). Among the annotated repeats, long terminal repeat (LTR) retrotransposon elements were the most abundant, accounting for 15.73-41.31% of each genome (Fig. 1c).

### Phylogeny, demography and TE landscape of the genus *Populus*

We constructed the phylogenetic tree using the reference genomes of 19 species/sub-species, based on both the concatenation and coalescent methods, using 2,455 single-copy orthologous genes. However, we observed a conflict with respect to the basal section in trees constructed using the two methods (Extended Data Figs. 2a,b). We thus quantified the amount of incongruence between individual gene trees and the species tree by implementing gene and site concordance factors (gCF and sCF) analysis, which respectively quantify the proportion of informative gene trees (gCF) and sites (sCF) that support a given branch between taxa. Both gCF and sCF were found to be quite low for many branches (Fig. 1b), and the incomplete lineage sorting (ILS) in conjunction with the short internal branches observed are probably the main factors causing the tree discordance, although ancestral gene flow could also play a minor role. To further resolve the phylogenetic challenges, we extracted approximately 48.7 Mb of orthologous regions from the reference-free Cactus^21^ alignments and used a much larger dataset of 11,385 low-copy number genes for phylogenetic analyses. Both methods yielded topologies consistent with the ASTRAL coalescent tree described above (Fig. 1b and Extended Data Fig. 2), implying that the coalescent-based phylogenetic approach is likely to be more robust in the presence of ILS. Three major clades, represented by *P. trichocarpa* (Clade-Ⅰ), *P. davidiana* (Clade-Ⅱ) and *P. euphratica* (Clade-Ⅲ), were highly supported, which also differ distinctly from each other in the phenotypic variation of leaves (Fig. 1). Further, we examined changes in effective population sizes (*N_e_*) of the 19 species in the recent past using the Pairwise Sequentially Markovian Coalescent (PSMC) method^22^. We found that different *Populus* species experienced a highly distinct demographic history (Extended Data Fig. 3), even for species occupying similar ecological niches.

We further compared the genomic distributions of TEs across the genus phylogeny (Figs. 1c,d and Supplementary Fig. 5). TE families tend to reside preferentially within 2kb surrounding genic regions, with the exception of the basal clade (Clade-Ⅲ) comprising *P. euphratica*, *P. pruinosa* and *P. illicifolia* in which *Gypsy* retrotransposons were located mainly in regions more distant from genes. At the superfamily level, different families were found to have varying relationships with gene expression, with expression in general being positively correlated with the distance to the nearest *Gypsy* and *Copia* superfamilies, negatively correlated with Helitrons, and only weakly correlated with other superfamilies (Supplementary Table 6). Therefore, potential *cis*-regulatory influences of TEs on the expression level of neighboring genes may differ between different superfamilies^23^. We also compared the proportion of TEs between shared and species-specific genomic regions. We observed a much higher content of TEs that are largely dominated by *Gypsy* LTR retrotransposons (LTR-RTs) in species-specific sequences compared with shared sequences (Extended Data Figs. 4a-c). We detected an average of 1,370 intact full-length LTRs (fl-LTRs) per species (Supplementary Table 7) and identified relatively recent retrotransposon amplifications (<5 million years ago, Ma) in most species (Supplementary Fig. 6). In particular, the species-specific fl-LTRs were much younger than the shared ones (Extended Data Fig. 4d). After identifying pairwise shared syntenic fl-LTRs between species, we detected 0-39.4% of still syntenic fl-LTRs across species (Supplementary Fig. 7). Differences in pairwise shared numbers were highly consistent with phylogenetic relationships and these results reinforce the notion that TEs, and in particular LTR-RTs, are important drivers of rapid sequence turnover and genome evolution in the genus.

### Evolutionary architecture of the *Populus* super-pangenome

We performed pan-genomic analyses of the 19 species/sub-species at the genus level, which we refer to as a super-pangenome. We annotated 40,606 gene families and 20,928 unassigned genes containing a total of 712,487 genes. The number of pan-genes retained increased with each additional genome added (Fig. 2a). On the basis of their presence in each genome, the gene families were further categorized into four groups: 12,924 gene families that were present across all 19 genomes were defined as core genes; 4,874 families present in 17 to 18 genomes as softcore genes; 19,668 families presented in 2 to 16 genomes as dispensable genes; and the remaining gene families only present in a single genome and the genes were not clustered into families as private genes (Fig. 2b, Extended Data Fig. 5a and Supplementary Table 8). We found that the proportion of each group of genes was highly consistent across species, with an average of 51.3% (SE=0.74%) belonging to the core genes, 22.8% (SE=0.33%) to the softcore genes, 22.3% (SE=0.97%) to the dispensable genes, and 3.6% (SE=0.59%) to the private genes (Supplementary Table 8). Compared to most of the core and softcore genes that are highly syntenic to the sister genus *Salix* (85.6% and 75.7%), dispensable and private genes show much lower syntenic ratios (29.7% and 17.7%) with *Salix* (Extended Data Fig. 5b). When compared with core and soft-core genes, dispensable and private genes had shorter coding sequences and were relatively closer to proximal upstream TEs (Figs. 2c,d). Moreover, dispensable genes exhibited higher nucleotide diversity (*π*) and a higher ratio of nonsynonymous to synonymous substitution (*d*_N_/*d*_S_) than core genes (Fig. 2e and Extended Data Fig. 5c). Expression analysis showed that dispensable and private genes displayed significantly lower expression levels but higher tissue specificity (Tau index) when compared to core and softcore genes (Figs. 2f,g). Nevertheless, caution should be warranted given the potential impact of gene prediction accuracy on the gene gain and/or loss measures across divergent species at the genus level. To minimize these affections, comprehensive annotation are supposed to be build by using the same gene prediction pipelines for all species with high-quality genome assemblies in the future.

**Fig. 2.**
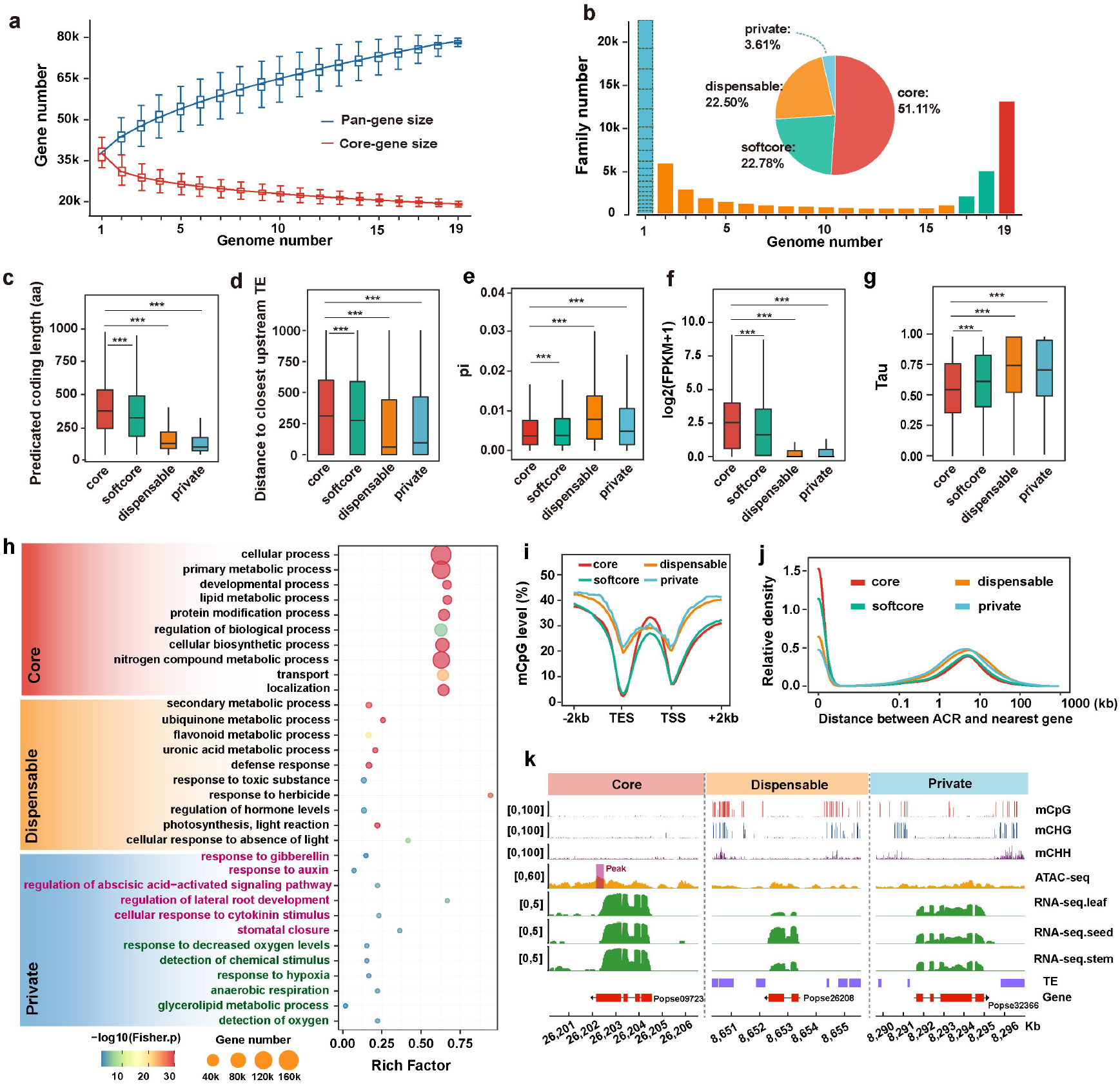
Pan-genome analysis at the gene space. **a,** Number of pan-genes and core-genes for different combinations of species. **b,** Proportion of pan-genes in the core, softcore, dispensable, and private categories. A blue block with a dashed border indicates unclustered genes and genes only clustered in a single genome. **c-g,** The distribution of CDS length (c), distance to the closest upstream TE (d), nucleotide diversity (π) (e), the expression level in leaf tissues (f), the tissue specificity index (Tau index, at least with three tissues available) (g) in core, softcore, dispensable and private genes. **h,** Representative Gene Ontology (GO) enrichment categories of the core, dispensable and private genes (pink fonts: *P. qiongdaoensis*; green fonts: *P. pseudoglauca*). **i,** Regional methylation (CpG) levels across gene-body and flanking regions, which were estimated by appropriately dividing into 30 and 20 equal bins respectively, for genes from various pan-gene categories. **j,** Frequency distribution of accessible chromatin regions (ACRs) and their distance to the nearest genes of genes from various pan-gene categories. **k,** Representative example of differences in gene expression, TEs distribution, methylation levels and ACRs among different pan-gene categories. The coverage tracks were visualized by pyGenomeTracks^135^. The statistical difference between groups was calculated using Wilcoxon ranked sum tests: \*\*\**P* ≤ 0.001. Results for each species are shown in the Supplementary Figs. 8-15.

Functional enrichment analysis showed that core genes were enriched in basic biological and cellular processes, including primary metabolic, developmental and other fundamental metabolic and biosynthetic processes. In contrast, the dispensable and private genes were enriched in areas related to secondary metabolic processes, abiotic and biotic responses, molecule transport and rhythmic processes (Fig. 2h and Supplementary Table 9). In particular, we found that private genes were highly enriched for processes that could be associated with species-specific adaptation to local environments. For instance, the private genes of *P. qiongdaoensis*, which naturally grows in tropical regions, were enriched in multiple processes involved in heat stress response, such as lateral root development, stomatal closure and phytohormone regulation (including response to auxin, cytokinin, abscisic acid and gibberellic acid). In addition, the private genes of *P. pseudoglauca,* which is mainly distributed on the Qinghai-Tibet Plateau and adjacent highlands, were significantly enriched in processes related to hypoxia and rapid temperature change responses, such as response to oxygen levels, anaerobic respiration and glycerolipid metabolic processes (Fig. 2h and Supplementary Table 10).

We further characterized the patterns of DNA methylation and chromatin accessibility across *Populus* species (Supplementary Tables 11-12), in order to compare the epigenetic marks and transcriptional regulatory elements among the different types of pan-genes (Figs. 2i-k and Extended Data Figs. 5d-f). Methylated bases were identified in three sequence contexts: CG, CHG, and CHH (where H is A, T, or C) based on the Bisulfite sequencing data from leaf tissue of 13 species. We found that both CHG and CHH methylation levels, together with CG methylation in the flanking regions of core and softcore genes were substantially lower than in dispensable and private genes (Fig. 2i and Extended Data Figs. 5d,e), which was expected since DNA methylation in these contexts is usually associated with repression of gene expression^24^. In contrast, CG methylation within protein coding sequence regions was similar or even higher in core and softcore genes compared to dispensable and private genes. This pattern mirrors previous studies that have shown that gene body methylation (gbM) in the CG context is positively correlated with levels of gene expression^25–27^, again suggesting that gbM may play an important role in the maintenance of the core genes that are generally enriched for housekeeping functions across species. In addition, as active *cis*-regulatory elements are widely reported to be located within accessible chromatin regions (ACRs), we performed ATAC-seq using leaf tissue from 12 *Populus* species to assess genome-wide chromatin accessibility and identify ACRs within each species (Extended Data Fig. 5f). ACRs were highly enriched upstream of transcription start sites of genes. Interestingly, we observed significantly shorter distances between ACRs and the nearest core and softcore genes relative to dispensable and private genes (Fig. 2j). Overall, these pan-gene results are consistent in all poplar species (Supplementary Figs. 8-15), suggesting that the epigenetic and regulatory architectures may both have pervasive effects on gene evolution as species diverged and adapted to a wide range of ecological niches.

### The evolutionary dynamics of duplicate genes alongside species divergence in the genus *Populus*

*Populus* and its sister genus *Salix* shared a recent WGD event around 58 Ma^8, 9^. In addition to WGD, other modes of gene duplication are also prevalent in various plant genomes^28–30^. We identified different modes of gene duplication across the 19 poplar genomes using DupGen_finder^31^. We found 14,674-22,148 (41.77-57.3%), 2,226-4,757 (5.99-12.06%), 1,015-3,185 (3.01-7.10%), 1,518-5,423 (4.37-15.44%) and 4,630-6,138 (11.55-17.29%) duplicated genes derived from WGD, tandem duplicates (TD), proximal duplicates (PD), transposed duplicates (TRD) and dispersed duplicates (DSD), respectively. In addition, 2,854-6,387 (8.66-17.13%) genes were only present once in the genome-wide landscape (referred to as singletons) (Fig. 3a and Supplementary Table 13). When mapping these duplicated genes and singletons to the pan-genes identified above, we found that core genes were mainly composed of WGD-derived genes, while singletons accounted for the majority of private genes (Fig. 3a). Moreover, the WGD-derived genes were mostly syntenic to *Salix*, whereas this was rarely the case for genes duplicated through other modes and for singletons (Extended Data Fig. 6a). These findings suggest that, compared to other genes, WGD-derived genes are more conserved at both species and genus level.

**Fig. 3.**
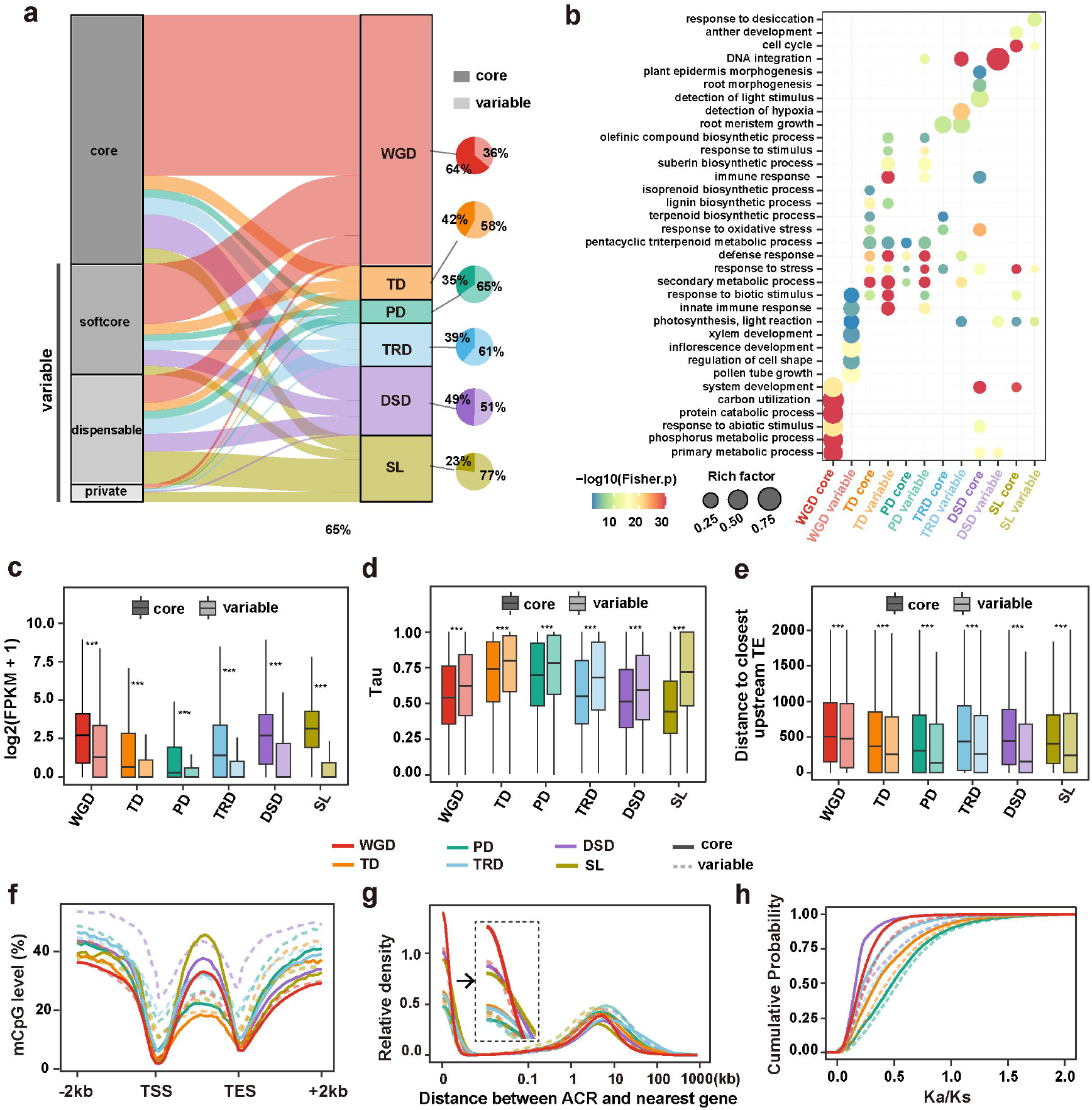
The landscape of gene duplication in poplar genomes. **a,** Overall proportions of different types of duplicated genes (WGD, TD, PD, TRD, DSD and singleton genes) across the four categories of pan-genes, where softcore, dispensable and private genes were all grouped into variable genes. **b**, Representative GO enrichment categories for the different classes of duplicated genes and pan-genes..**c-h,** The comparison of average expression level (log2 FPKM in leaf tissue) (**c**),tissue specificity index (Tau index, at least three tissues) (**d**), the distance to closest upstream TE (**e**), regional methylation (CpG) levels across gene-body and flanking regions (**f**), the frequency distribution of ACRs and their distance to the nearest genes (**g**) and the *K*a/*Ks* distribution (**h**) across different types of duplicate genes intersect with the pan-gene categories (core vs. variable). WGD whole-genome duplication, TD tandem duplication, PD proximal duplication, TRD transposed duplication, DSD dispersed duplication, SL singletons. The statistical analysis was performed using Wilcoxon ranked sum tests: \**P* ≤ 0.05; \*\*\**P* ≤ 0.001. Results for each species are shown in the Supplementary Figs. 16-26.

We also performed integrative genomic analysis, incorporating expression, methylation and chromatin accessibility, to compare these duplicate genes with different origin modes. Regardless of the pan-gene type, the WGD-derived genes had, on average, longer CDS lengths, a greater distance to proximal upstream TEs, lower *K_a_/K_s_* ratios and exhibited higher expression levels and lower expression specificity compared to other, i.e. TD- and PD-derived, duplicate genes (Figs. 3c-e,h and Extended Data Fig. 6b-d). The average methylation levels in flanking regions of WGD-derived genes at CG sites and along the whole gene at CHG and CHH sites were substantially lower than the methylation levels of the other groups of duplicate genes, which are probably associated with fewer TEs near these genes, which may also explain their higher expression levels (Fig. 3f and Extended Data Figs. 6e,f). Moreover, compared with duplicate genes originating in other ways, there was a significantly shorter distance between ACRs and the nearest core-type WGD-derived genes (Fig. 3g). These patterns are consistent in all poplar species (Supplementary Figs. 16-26), further support the suggestion that expression and epigenetic regulatory architectures play key roles in determining the distinct evolutionary trajectories of duplicated genes. To further explore the biased functional roles of duplicate genes with different origin modes, we performed GO enrichment analysis with the results suggesting that TD- and PD-derived duplicate genes exhibited divergent functional roles and were enriched for GO terms related to secondary metabolic processes, response to stress and biotic stimulus when compared to the WGD-derived duplicate genes that were mainly enriched in essential functions (Fig. 3b and Supplementary Table 14).

As WGD is considered as a major driving force in organismal adaptation and species diversification^32^, we thus performed complementary and integrative analysis combining pan-genes and duplicate genes to examine the differential retention and divergent resolution of duplicate genes between species following WGD. We used one-to-one duplicated pairs (i.e., both gene copies appeared only once in all WGD-derived gene pairs) that are originated from WGD for this analysis (Supplementary Table 15). According to the level of synonymous divergence (*K*_s_) between the two paralogs, we divided duplicate genes into three groups and found that WGD pairs with highest *K*_s_ values tend to enrich in dispensable and private gene sets compared to those with lower *K*_s_ values (Figs. 4a,b). Furthermore, the relative gene expression and DNA methylation divergence (in particular of gbM) between the duplicates increased with their evolutionary changes. More strikingly, the duplicate pairs with the different pan-gene types (e.g. one paralog is core gene and the other paralog is dispensable gene) across species exhibit higher expression and methylation divergence than those sharing the same pan-gene types (Figs. 4c,d and Extended Data Fig. 6g). These results imply that potential evolutionary novelties of duplicated genes also accompanied by clade- or species-specific gene retention, losses and functional innovations over the different speciation events (Extended Data Fig. 6h). Additionally, we observed a positive correlation between the expression and methylation divergence of duplicated genes in all three sequence contexts but especially for the CG methylation in gene-body regions (Fig. 4e and Extended Data Fig. 6i). These findings have universality across poplar species (Supplementary Figs. 27-29), suggesting that DNA methylation, particularly gbM, has strong effects in determining the functional divergence and long-term preservation of duplicated genes^33, 34^.

**Fig. 4.**
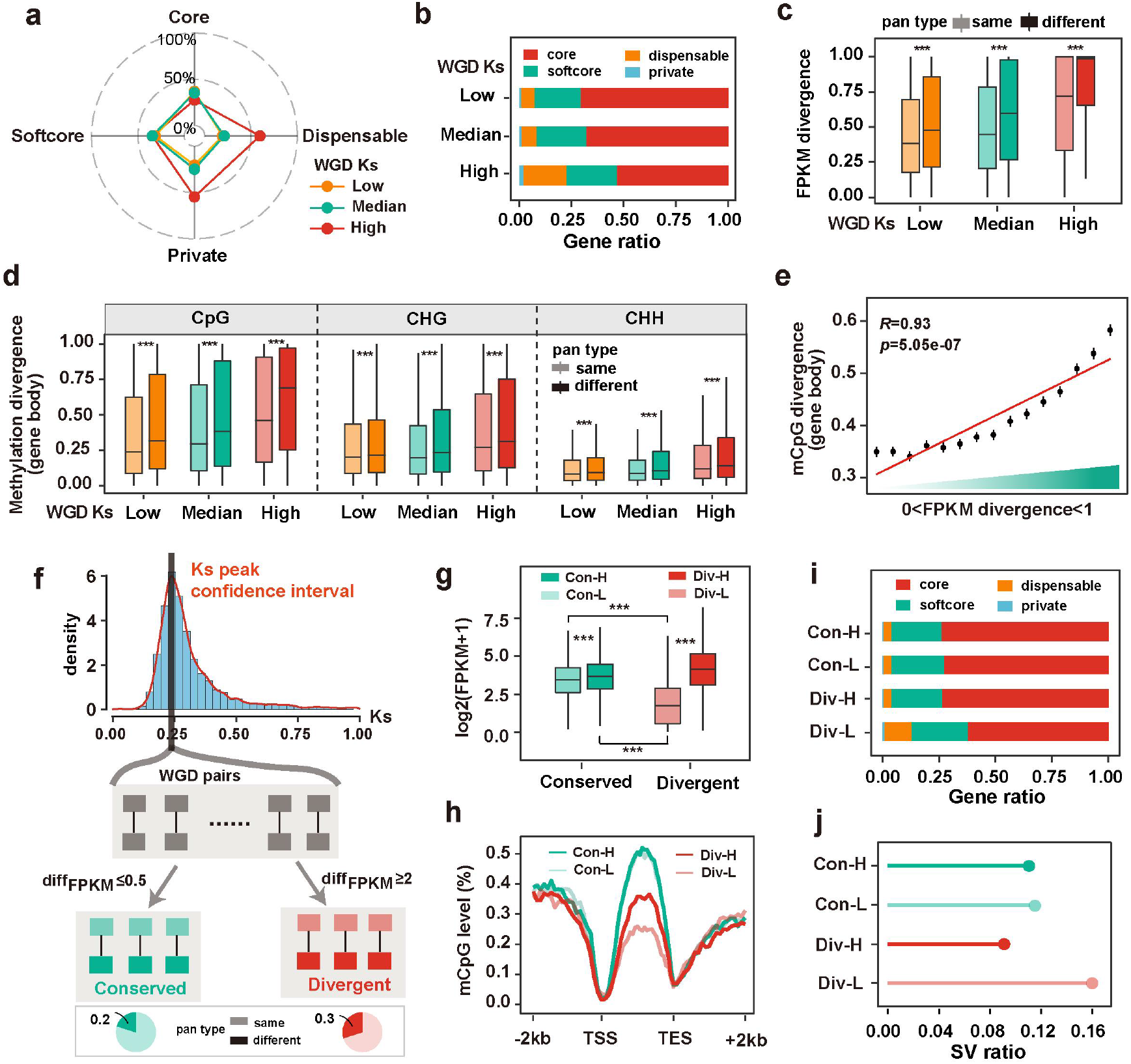
The evolutionary fates and divergent resolution of duplicate genes between species following WGD. **a,b,** Overall proportion of WGD-derived duplicated genes classified by evolutionary age (estimated by calculating *Ks* values between paralogs) across the four pan-gene categories. **c,d,** Association of expression (c) and methylation (d) divergence of duplicated genes with evolutionary age and the consistency pan-gene status between paralogs. **e,** Pearson’s correlation coefficient between methylation divergence (CG) and expression divergence of duplicated genes in gene body regions. **f,** The selection of WGD pairs under the control of evolutionary age effects. diffFPKM was calculated by formula: (FPKMhigh - FPKMlow)/ FPKMlow, where FPKMHigh and FPKMlow denote the genes with relative higher and lower expression in the duplicated pairs, respectively. Gene pairs with diffFPKM ≤ 0.5 were considered as conserved whereas those ≥ 2 were assumed to be divergent. The pie chart denotes the proportion of paralogs with same or different pan-gene type within conserved and divergent duplicate gene pairs respectively **g**, The expression level (log2 FPKM in leaf tissue) of genes in divergent and conserved gene pairs. The legends “Con-L”and “Con-H” respectively represent genes with low and high expression in the conserved gene pairs, while “Div-L”and “Div-H” respectively represent genes with low and high expression in the divergent gene pairs (same in h-j). **h,** Regional methylation (CG) levels across gene-body and flanking regions of genes in divergent and conserved gene pairs. **i,** Overall proportion of genes from the four pan-gene categories in distinct WGD-derived genes based on expression divergence and expression level. **j,** The ratio of structural variation (SV) identified across the 16 *Populus* species by SyRI among different categories of WGD-derived genes divided according the expression divergence and level. The statistical analysis was performed using Wilcoxon ranked sum tests: ns.*P* > 0.05; \*\*\**P* ≤ 0.001. Results for each species are shown in the Supplementary Figs. 27-29

In support of this, we further selected only duplicate gene pairs located within the confidence interval of corrected *K_s_* peaks for analysis to corrected for the sequence divergence(Fig. 4f). Among duplicate gene pairs that share the greatest sequence similarity, we still found extensive variation in expression divergence (Extended Data Figs. 7a,b). To investigate the factors driving the differences in gene expression divergence, we classified duplicate genes into conserved and diverged pairs by assessing the similarity in expression profiles of partners (Figs. 4f,g). Despite divergent duplicate gene pairs showing similar overall expression levels with conserved pair, they still exhibited significantly higher *K_a_/K_s_* ratios and tissue expression specificities (Extended Data Figs. 7c-g), suggesting that functional divergence probably occurred at both expression and protein level for these divergent duplicate pairs. However, compared to the weak differences in nearby TE, chromatin accessibility distribution and methylation in the CHG and CHH sequence contexts between divergent and conserved duplicate gene pairs (Extended Data Figs. 7h-k), we observed remarkable divergence of CG methylation in the gene-body regions (Fig. 4h), with divergent pairs displaying much lower and divergent levels of gbM when compared with the similar levels of gbM observed for conserved pairs. Moreover, levels of duplicate-gene retention and sequence divergence across species also differed between conserved and divergent duplicate pairs, with the latter having higher tendency of containing different pan-gene categories and structural variations (SVs) across species (Figs. 4f,i-j). Together, these findings suggest that the loss of gbM is likely a key epigenetic precursor for sub- or neofunctionalization of duplicates following WGD, particularly given that rapid reduction of gbM may result in aberrant transcription, splicing and TE insertion^35^, which extends the window of opportunity for subsequent differential expression and sequence divergence via mutation^36, 37^. Nevertheless, some caution is warranted given the non-independence of duplicated genes across species and also more future work needs to dissect the precise cause or consequence roles of epigenetic modification underlying retention and functional divergence of duplicated genes.

### The landscape of structural variations (SVs) and hemizygous genes in *Populus*

We identified SVs based on 16 chromosome-level genomes by comparing 15 species against the *P. trichocarpa* reference genome to identify syntenic and rearranged regions using SyRI^38^. We found highly consistent patterns when using an alternative species, *P. adenopoda*, as the reference (Supplementary Figs. 30-38). In general, species that have a closer phylogenetic relationship to the reference had more syntenic regions and lower sequence divergence (Extended Data Fig. 8a). Five types of representative SVs (>50 bp) were extracted, including insertions (INS), deletions (DEL), inversions (INV), translocations (TRANS) and duplications (DUP). The total sizes of SVs varied between 49.4 Mb and 184.6 Mb across the species compared (Supplementary Table 16), with the length per SV ranging from a few dozen bp to hundreds of Kb or even over Mb scales (Extended Data Fig. 8b). To verify the accuracy and consistency of the SVs when called based on different reference genomes, we randomly sampled 1000 different SVs detected by using *P. trichocarpa* as the reference and then looked at how well those SVs were called when using *P. adenopoda* as the reference. The results showed that only 9 SVs (9/1000) were not cross-validated (Supplementary Table 17). Furthermore, we selected 50 SVs and manually checked by mapping Nanopore long reads to the genome assemblies. We found that only one border (1/100) could not verified in the mapping results (Supplementary Table 18 and Extended Data Fig. 8c). We next merged all SVs into a set of 142,202 nonredundant SVs, comprising 34,372 insertions, 40,811 deletions, 24,623 translocations, 1,107 inversions and 41,289 duplications. When using the total number of genes in *P. trichocarpa* as a reference, 77.6% of which (26,933) harbored at least one SV across different species. We observed a remarkable uneven distribution of SVs across the genome, with some regions being highly collinear among species whereas other regions act as SV hotspots, harboring more TEs and fewer collinear genes (Fig. 5a and Extended Data Figs. 8d,e). Many functional genes were identified to be overlapped with at least one SV across species (Supplementary Table 19). GO analyses of these genes within SV hotspot regions indicated that defense response, secondary metabolite biosynthesis and signal transduction in both adaptation and development were enriched (Supplementary Table 20), suggesting an important role of SVs in adaptation of the sampled *Populus* species to biotic and abiotic stresses, and the result of significantly higher ratios of disease resistance genes (or *R* genes) overlapping with SVs also supports this (Extended Data Fig. 8f).

**Fig. 5.**
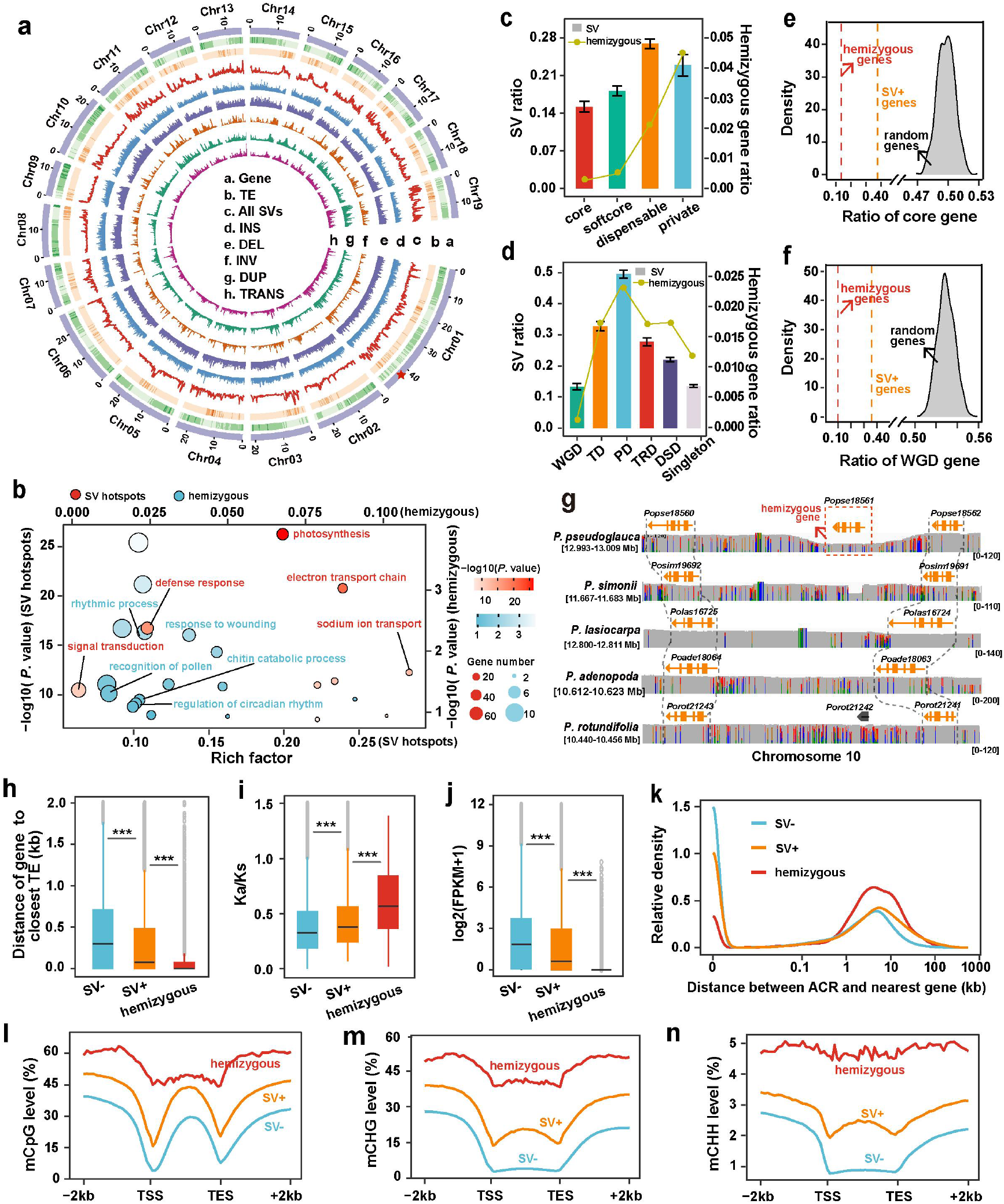
The landscape and genomic features of structural variations (SVs) and hemizygous genes. **a,** Circos plot of the densities of genes, TEs, all SVs, insertions (INS), deletions (DEL), inversions (INV), duplications (DUP) and translocations (TRANS) across different *Populus* species when using the *P. trichocarpa* genome as the reference. The position of the red star represents an fixed inversion between species from Clade-Ⅰ and Clade-Ⅱ detained in Fig.6. **b,** Representative GO enrichment categories of the genes overlapped with SV hotspots regions and hemizyous regions. **c,** The ratio of SV-related and hemizyous genes in core, softcore, dispensable and private genes. **d,** The ratio of SV-related and hemizyous genes in different classes of duplicated genes. **e,f,** The density of ratios of core-gene (e) and WGD gene (f) overlapped with SV and hemizyous genes (dashed lines indicate the empirical observation) compared with the distribution resulting from the 1,000 randomizations. **g.** Integrative Genomics Viewer (IGV) plot showing the coverage of reads and the structure of a specific hemizygous gene in *P. pseudoglauca*. In each diagram, the arrow denotes the gene, and the connected grey lines indicate the homologous genes. The red box indicates the hemizyous gene. The height represents the read coverage. **h-n,** Comparison of the distance to closest TE (h), selective constraint (i), expression level (j), ACR distributions (k), methylation levels at CpG (l), CHG (m) and CHH (n) context of hemizygous genes as well as genes with SVs (SV+) and without SVs (SV-). SV+ and SV-genes respectively denote the gene-body regions overlapping and not overlapping the SVs. The statistical analysis was performed using Wilcoxon ranked sum tests: \*\*\**P* ≤ 0.001. Results for each species are shown in the Supplementary Figs. 30-38.

An extreme pattern of SVs within a given genome is that cause hemizygous genes, which can be inferred and evidenced from long-read mapping^39^. Based on remapping Nanopore long reads to the corresponding reference genome, we estimate that 0.63-1.42% (211-559) of genes are hemizygous (i.e., gene presents in only one of the two homologous chromosomes) across 10 *Populus* species with long-reads datasets available (Supplementary Table 21), which is slightly higher than domesticated selfing rice (0.35%–0.73%)^40^, but much lower than the clonal propagated grapevine (∼15.5%)^39^ and apomictic sweet orange (∼11.2%)^41^. The relatively lower proportion of hemizygous genes is not surprising given that the features shared by most *Populus* species, such as outcrossing mating system, widespread geographic distribution and higher efficacy of purifying selection acting against deleterious variations^42^. In particular, we found that hemizygous genes were significantly concentrated in SV hotspot regions than expected at random (Extended Data Fig. 8g), suggesting similar mechanisms driving the formation of SVs within and across genomes in specific regions. Functional enrichment analysis on these hemizygous genes are linked to GO terms such as “rhythmic process”, “regulation of circadian rhythm” and ‘response to wounding’, implying their possible adaptive roles in immunity and stress response (Fig. 5b and Supplementary Table 22). For instance, the gene *Popse18561*, the ortholog in *Arabidopsis PCR2* that was known to be associated with the response to oxidative stress^43^, was specifically expanded hemizygously in high altitude *P. pseudoglauce* (Fig. 5g).

When comparing the ratio of genes associated with SVs and hemizygous genes among the pan-genes and duplicated genes categories, we observed that core-genes and WGD-derived genes were significantly depleted in genes overlapping with SVs and hemizygous state when compared to dispensable-genes and the small-scale duplicated genes that are more likely to affected by structural variants than expected by chance (Figs. 5c-f and Extended Data Figs. 8h,i). Moreover, genes associated with SVs and particularly the hemizygous genes were found to be much closer to TEs than other genes (Fig. 5h), again supporting the idea that TE mobilization is likely to be a constant and dominant mechanism for the formation of SVs, finally leading to the massive gene presence-absence variation within and between species^44^. We also found that genes with SVs, especially those hemizygous genes, had significantly higher molecular evolutionary rates (*K_a_/K_s_*), lower expression level, longer distance to nearest ACRs and higher methylation level within both gene-body and flanking regions when compared to genes without SVs (Figs. 5i-n). As such, these results suggest that the transposon-dominated formation of SVs has a broad influence on gene function by affecting the expression of nearby genes through altering both coding and flanking regulatory sequences, as well as triggering changes of local chromatin and the epigenome^45–48^.

### Identification of a *cis*-regulatory SV potentially underlying leaf margin differences across species in *Populus*

SVs have been reported to have widespread impacts on gene expression and functional traits^49–51^. Among the SVs detected across species, we detected an inversion (∼104 kb) on chromosome 1 that occurred between species of Clade-Ⅰ (mainly belonging to sect. *Tacamahaca*) and Clade-Ⅱ (mainly belonging to sect. *Populus*) (Fig. 5a). Interestingly, species in these two clades have contrasting leaf margins (Fig. 6a). This inverted region was also confirmed by mapping the Nanopore long reads of the representative species from the two clades to the *P. trichocarpa* genome (Extended Data Fig. 9a). We further measured and compared the synteny diversity, which quantified the degree of genomic collinearity, among species within and between the two clades. Compared to the flanking regions, we observed high inter-clade, but not intra-clade, synteny diversity across the entire region, which is consistent with the presence of an inversion that suppressed recombination and contributed to divergence (Fig. 6b).

**Fig. 6.**
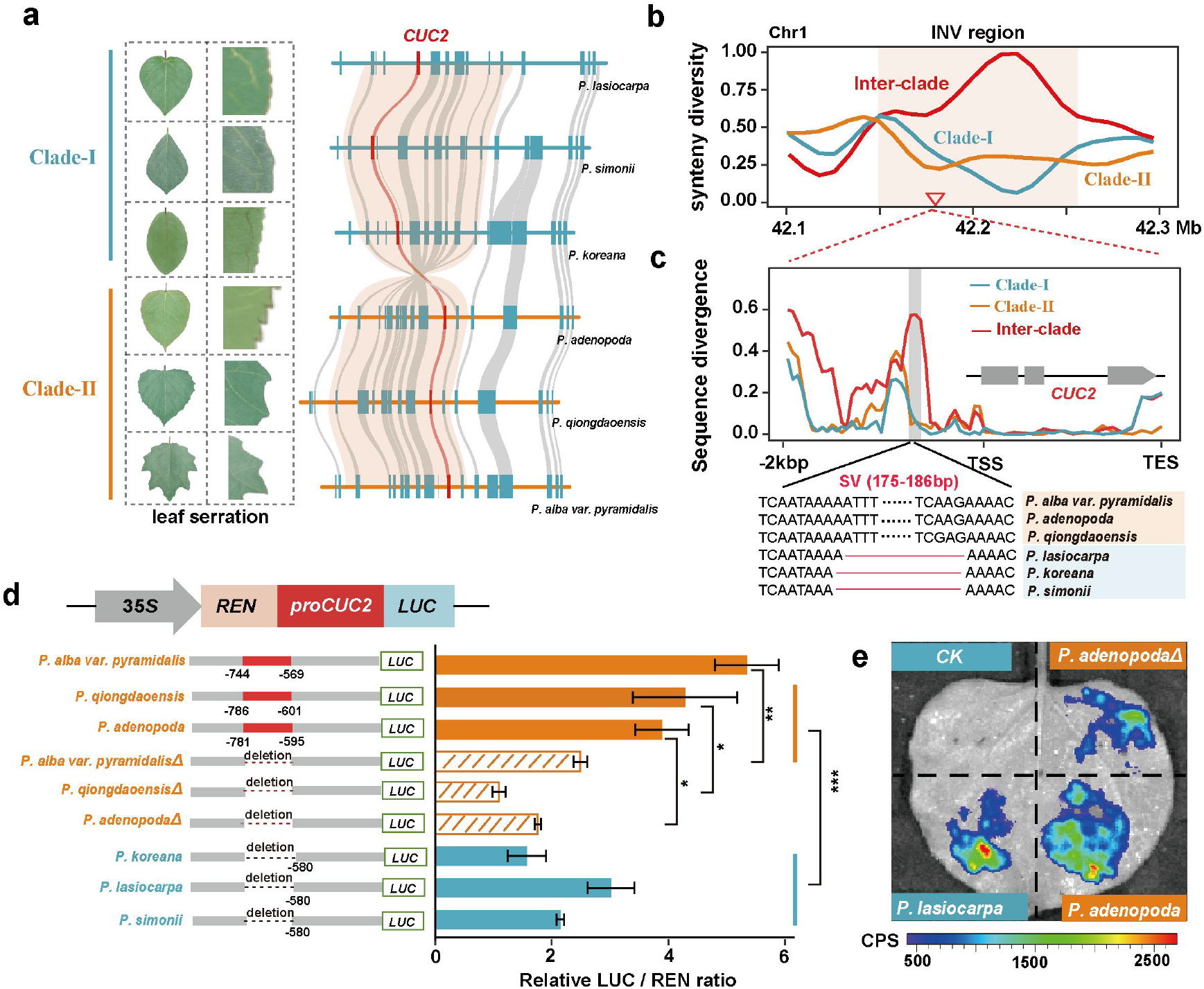
Identification of a *cis*-regulatory SV potentially underlying leaf margin differences across species in *Populus*. **a,** Identification of a fixed inversion (the location was shown as red star in Fig.5a) that includes the *CUC2* gene between species from Clade-Ⅰ and Clade-Ⅱ as shown in Fig. 1b that have contrasting leaf margin phenotypes. **b,** The synteny diversity within and between clades around the inverted region. **c,** The divergence in the promoter and gene-body regions of *CUC2* highlight a ∼180bp presence/absence variant closet to the transcription start site (TSS) being fixed between species from the two clades. **d,** Validation of function of the ∼180bp presence upstream of *CUC2*. Schematic overview of luciferase constructs with and without the ∼180bp *CUC2* promoter, with corresponding luciferase activity in transfected *N. benthamiana*. Each bar represents the mean of three independent experiments ± SD (\**P* ≤ 0.05, \*\**P* ≤ 0.01, Student’s t test). **e,** Transient expression assay show that the ∼180bp deletion directly decreased the regulatory potential of the promoter on the *CUC2* gene. Representative images of *N. benthamiana* leaves 72 h after infiltration were shown.

Among the ten genes contained in the inverted region, we identified an orthologous gene of the transcription factor *CUP-SHAPED COTYLEDON2* (*CUC2)*, which has been repeatedly shown to play an essential role in promoting leaf margin outgrowth and the formation of serration in plants^52–54^. As noted above, there are striking morphological differences in leaf margin serrations between species from the two clades. Most species of the Clade-Ⅰ have smooth and entire margins whereas those Clade-II ones have more serrated leaves with sparse or dense sinuous teeth (Fig. 6a), which may reflect their long-term adaptation to different natural environments and habitats. The divergent patterning of leaf margin between the two clades could likely be governed by the gene *CUC2*. Further testing *CUC2* expression along the leaf margin of emerging young leaves revealed significantly higher expression of *CUC2* in species from the Clade-Ⅱ than those from Clade-Ⅰ (Extended Data Figs. 9c,d). This is consistent with the previous finding that the elevated expression of the *CUC2* probably further triggers the formation of more serrated leaf margins in *Arabidopsis*^53, 54^. Notably, compared to small sequence variations observed between the two clades within *CUC2* genic regions, the divergence was much higher in the promoter region, especially a ∼180bp presence/absence variant close to the transcription start site (TSS) was identified between species from the two clades (Fig. 6c). We therefore speculated that the ∼180bp presence/absence variant might have regulated the inter-species *CUC2* expression differences. To test this hypothesis, we cloned the promoter fragments (Supplementary Fig. 39) from three species belonging to Clade-Ⅰ and three species belonging to Clade-Ⅱ and conducted a dual luciferase assay (LUC) via transiently transformed tobacoo (Figs. 6d,e and Extended Data Fig. 9e). LUC activity driven by the Clade-Ⅱ promoters was significantly higher than that from the Clade-Ⅰ promoters (Figs. 6d,e and Extended Data Fig. 9e). Importantly, the modified Clade-Ⅱ sequences components excluding the ∼180bp fragment (Clade-Ⅱ*Δ*) drove significantly lower luciferase reporter gene expression compared to constructs containing the fragment (Figs. 6d,e and Extended Data Fig. 9e). Therefore, these data suggest that the ∼180bp insertion that contain multiple transcriptional factor binding motifs in species from Clade-Ⅱ (Extended Data Fig. 9b) resulted in the elevated transcriptional activation of *CUC2*, thereby likely causing the more serrations along the leaf margin.

## DISCUSSION

The genus-level super-pangenome dataset of 19 *Populus* species/sub-species covering the major taxonomic clades from this study represents, to our knowledge, the first super-pangenome reconstructed based on whole-genome assemblies for forest trees (Fig. 7). The availability of high-quality genome assemblies of these species enabled us to precisely examine the evolution of the genomic landscape underlying the widespread adaptation and high phenotypic diversity of this genus. We revealed substantial variations in the number of genes and the content of repetitive regions between these *Populus* species. Variable TEs abundance largely explained variation in genome size and further promoted gene and genome evolution across different species. The constructed super-pangenome greatly expands the gene repertoires of the total genus, which can now be used in further diverse research areas. For example, many dispensable and private genes across different species were found to play critical roles in specific adaptation to abiotic and biotic stresses; these may be valuable for genetic breeding of poplars and for revealing novel adaptive mechanisms that can be translated to other species.

**Fig. 7.**
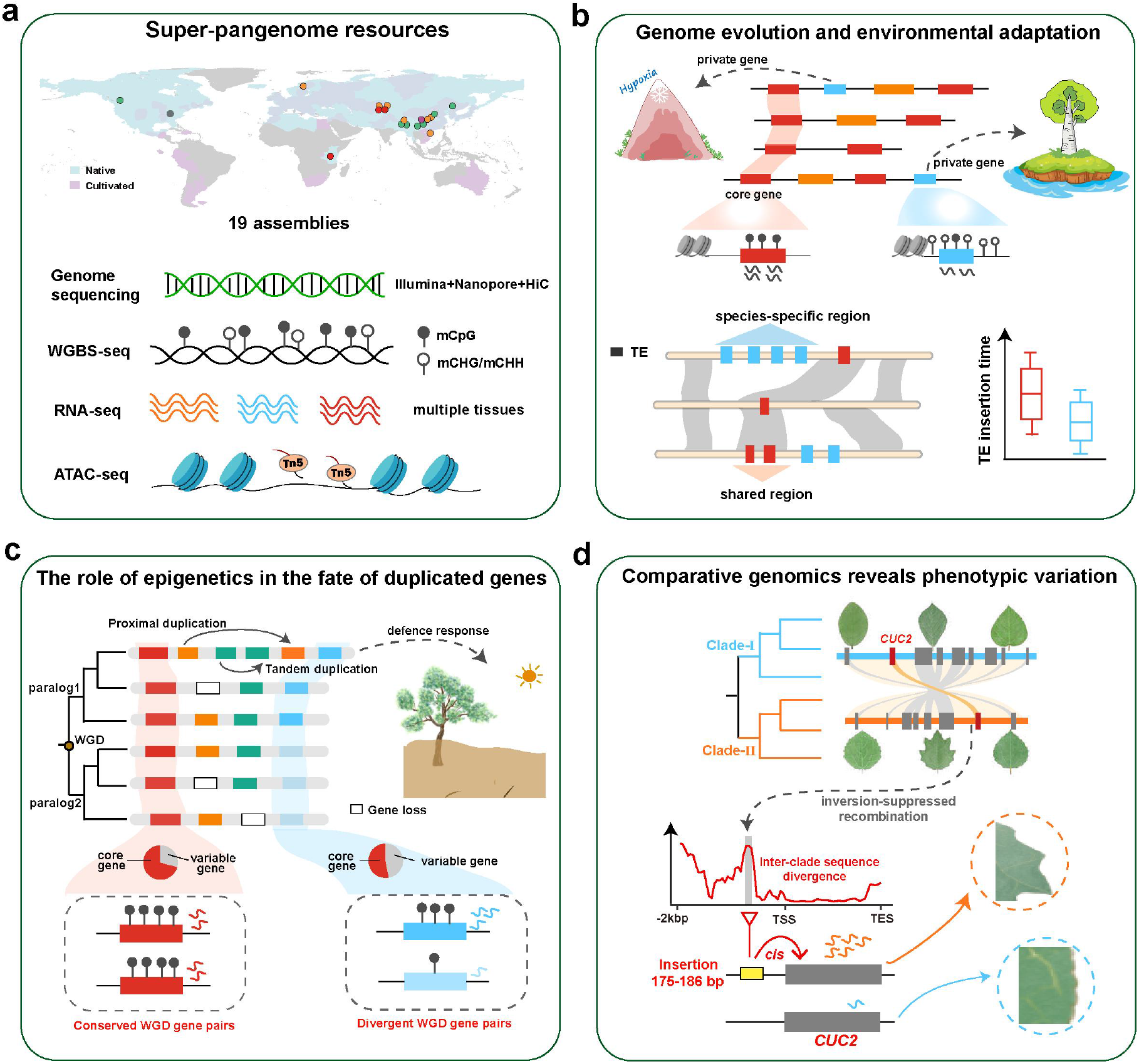
Summary of the four key findings in this study. **a,** The super-pangenome resources combined with the multi-omics datasets across the 19 *Populus* genomes generated. **b,** Integrative analysis associating pan-genomes with epigenetic and regulatory architecture to understand genome evolution and species adaptation. **c,** Complementary analysis of pan-genes and duplicate genes provides novel insight into understanding the genetic mechanisms that create functional divergence of duplicates retained alongside species divergence following whole-genome duplications (WGDs). **d,** The large-scale comparative genomics at the genus level opens vital opportunities for exploring how genomes evolved and diverged as species adapted to a wide range of ecological niches, and also enable better identifying previously hidden structural variants that affect phenotypic and functional divergence across species.

All *Populus* members shared a WGD event and we combined the analysis of transcriptomes and methylomes as well as chromatin accessibility mapping to illustrate the evolutionary trajectories of these duplicated genes after WGD across different species. We find that sequence evolution and further functional divergences of the WGD-derived duplicate genes are largely affected, regulated and maintained by the levels of gene-body methylation. The evolution and maintenance of these WGD-derived duplicate genes contribute to variable gene repertoires between species that may lead to their divergence with respect to both adaptation and phenotype. The evolutionary mechanism of these duplicated genes that we revealed deepens our understanding of the critical roles of the frequent WGD during plant diversification. We further identified 142,202 SVs across the sampled species, which overlap with substantial genes in the genus and are involved in diverse functions. Among which, 0.63-1.42% of genes were found be hemizygous. SVs tend to occur more frequently and in greater numbers within dispensable and small-scale duplicated genes, which are mainly involved in defense response, secondary metabolite biosynthesis and signal transduction. This further reveals the potential important regulatory roles of SVs in modulating species adaptation and resilience to environmental changes. One notable finding in our study was the identification of a large inversion-mediated *cis*-regulatory divergence of *CUC2* gene likely act as an essential regulator in driving divergent patterning of leaf margin serration across *Populus* species. Overall, the super-pangenome that we constructed here for this model tree genus provides insights into the evolution of the genomic landscape across different species, mainly involving variations in genes, SVs and TEs. These genetic resources will facilitate the genotyping of agronomically important traits at both population- and species-level for this genus, something that has rarely been explored in forest trees.

## ACKNOWLEDGMENTS

This work was supported by the National Key Research and Development Program of China (2022YFD2201200 to J.W. and 2021YFD2200202 to T.Y. and J.L.). National Natural Science Foundation of China (31971567 to J.W.) and Fundamental Research Funds for the Central Universities (SCU2022D003, SCU2021D006, SCU2019D013 and 2020SCUNL207 to J.W. and J.L.).

## AUTHOR CONTRIBUTIONS

J.W., T.Y., J.L. conceived the research. J.W. supervised the study. T.S., X.Z., C.J., Q.L., Z.L., Y.Z., J.Z. J.W. conducted genome assembly, annotation and bioinformatics analyses. S.L. J.F. and L.Z. collected samples of different species. Y. H., Y. J., X.D., J.F., L.C. carried out experiments. T.S. and J.W.wrote the manuscript, with the input from N.R.S., P.K.I. and J.L.. All authors proofread and approved the final manuscript.

## Methods

### Plant materials and genome sequencing

Nineteen *Populus* species were selected for the construction of the genus-level super-pangenome. In addition to 13 published assemblies, genomes of 6 more species were newly sequenced and assembled (Extended Data Fig.1 and Supplementary Table 1). Fresh young leaves were harvested and immediately frozen in liquid nitrogen, followed by preservation at -80 °C until DNA extraction for the newly sequenced species. High-quality genomic DNA was extracted from leaves using the CTAB method. For the short-read sequencing, 150-bp paired-end libraries with an insert size of 350 bp were generated according to the standard protocol (Illumina) and sequenced on an Illumina HiSeq X Ten platform. All raw reads were filtered using fastp (v0.20.0)^55^ with the following criteria: (1) with adapter sequences, (2) containing N base and (3) more than 20% of bases with a quality score <20.

For the long-read sequencing, libraries for Nanopore long reads sequencing were built using large (>20 kb) DNA fragments with the Ligation Sequencing Kit 1D (SQK-LSK109), and sequenced using the PromethION platform (Oxford Nanopore Technologies). Low-quality nucleotides that a mean quality score <7 were removed.

For the Hi-C experiment, the libraries were constructed from about 3g of fresh and young leaves and prepared with DpnII restriction enzyme, followed by sequencing on the Illumina NovaSeq platform. All clean Hi-C reads were obtained using fastp (v0.20.0)^55^ according to the following criteria: (1) with adapter sequences, (2) the number of N base ≥5 and (3) more than 40% of bases with a quality score <15. Total RNA was extracted from multiple tissues (leaf, root, stem, etc., the detailed information was described in Supplementary Table 1) using the CTAB method. RNA-seq libraries were constructed using a NEB Next Ultra Directional RNA Library Prep Kit and sequenced on the Illumina HiSeq 2500 platform with a read length of 2 × 150 bp.

### Genome assembly and pseudo-chromosome construction

The quality-filtered Illumina short reads were first used to estimate the genome size of each *Populus* species via a 17-bp k-mer frequency analysis with Jellyfish (v2.3.0)^56^. NextDenovo (v2.0-beta.1, https://github.com/Nextomics/NextDenovo), which contain two core modules (NextCorrect and NextGraph), was then used for the preliminary sequence assembly based on the Nanopore long reads. The raw long reads were first error-corrected via NextCorrect with parameters “reads_cutoff=1k, seed_cutoff=30k”, and then assembled via NextGraph with default parameters. To further improve single-base accuracy and obtain high-quality consensus sequences; assembled contigs were polished using Racon (v1.3.1)^57^ with long reads for three rounds, and further error-corrected using NextPolish (v1.0.5)^58^ with the Illumina short reads for four rounds. The redundant sequences were subsequently removed by using perge_haplotigs (v1.1.1)^59^ and the obtained genome assemblies were checked for DNA contamination by searching against the NCBI non-redundant nucleotide database (Nt) using BLASTN, with an E-value cutoff of 1e-5. Then, BUSCO (v4.0.5, embryophyta_odb10 download at 16-Oct-2020)^19^ and Merqury (v1.3)^20^ with default settings were both applied to assess the integrity of assemblies.

To generate pseudochromosome-level genomes, the clean Hi-C paired-end reads were first mapped to the draft assembled sequences using bowtie2 (v2.3.2)^60^ (parameters: -end-to-end --very-sensitive -L 30) to obtain the unique mapped paired-end reads. Then, valid interaction-paired reads identified by HiC-Pro (v2.11.4)^61^ were further used to cluster, order, and orient scaffolds onto 19 pseudochromosomes using LACHESIS^62^, with parameters CLUSTER MIN RE SITES=100, CLUSTER MAX LINK DENSITY=2.5, CLUSTER NONINFORMATIVE RATIO=1.4, ORDER MIN N RES IN TRUNK=60, ORDER MIN N RES IN SHREDS=60. Finally, placement and orientation errors exhibiting obvious discrete chromatin interaction patterns were manually adjusted.

### Repeat and gene annotation

Considering that the methods for TE annotation are highly variable across studies, we used a uniform workflow to perform whole-genome TE annotation for both the newly assembled and previously reported genomes, in order to minimize methodological artifacts for TE discovery. TE annotations were first derived using the Extensive de-novo TE Annotator (EDTA, v1.9.3) pipeline^63^, which combines well performing structure- and homology-based programs and subsequent filtering methods to create a high quality non-redundant TE library. TEsorter (v1.2.5)^64^ was then used to reclassify the TEs that were annotated as “LTR/unknown” by EDTA.

For gene prediction of newly assembled genomes, we first used RepeatMasker (v4.10)^65^ to mask the whole genome sequences with the TE libraries constructed by EDTA. An integrated strategy that combined *ab initio* prediction, homology-based prediction and transcriptome-based prediction was then used to predict the protein-coding genes based on the masked genomic sequences. For homology-based gene prediction, the published protein sequences of six species, including *Populus trichocarpa*, *Populus euphratica*, *Salix brachista*, *Salix purpurea*, *Arabidopsis thaliana* and *Vitis vinifera* were used. TBLASTN (ncbi-BLAST v2.2.28)^66^ with e-value less than 1e-5 was employed to align these protein sequences against the genome assemblies, and GeneWise (v2.4.1)^67^ was used with default settings to predict the exact gene models. For transcriptome-based prediction, trimmed RNA-seq reads from multiple tissues were mapped to the respective reference genome using HISAT (v2.2.1)^68^ with parameters “--max-intronlen 20000 --dta --score-min L,0.0,-0.4”, and Trinity (v2.8.4)^69^ was used for transcripts assembly with default parameters. These assembled transcripts were subsequently aligned to the corresponding genome to predict gene structure using PASA (v2.4.1)^70^. For *ab initio* gene prediction, Augustus was used with default parameters incorporating the homology- and transcripts-based evidence for gene model training. Finally, all the gene models generated by the above three approaches were integrated into a comprehensive gene set using EvidenceModeler (v1.1.1)^71^, and the resulting gene models were further updated using PASA for three rounds of iteration. Furthermore, BUSCO was used to evaluate the protein-coding annotations.

Functional annotation of predicted genes was conducted based on comparisons with the NCBI nonredundant protein database (NR), SwissProt, TrEMBLE, COG and KOG protein databases by using BLASTp with 1e-5 E-value cutoff. InterProScan (release 5.32-71.0)^72^ was used to identify functional domains and motifs. Gene ontology (GO) terms and KEGG pathways for each gene were assigned by InterProScan and KEGG Automatic Annotation Server, respectively. The topGO^73^ R package was used for downstream gene set enrichment analysis in this study.

To evaluate the genomic collinearity among species, we used MCScanX^74^ to detect syntenic gene blocks between each of the newly sequenced genomes and *P. trichocarpa* genome, as well as the collinearity between the 19 poplar genomes and *S. suchowensis*, with default parameters.

### Phylogenetic reconstruction

Three different datasets were used to retrieve the phylogenetic relationships of 19 poplar species, along with three willow species (*S. suchowensis*^75^*, S. purpurea*^76^ and *S. brachista*^77^) with published genome sequences available. First, amino acid sequences of the single-copy gene families (SCGs, n= 2,455), identified using the OrthoFinder pipeline (v2.5.2)^78^ under the default parameters after removing genes with early stop codons or open reading frames shorter than 50 amino acids, were aligned using MAFFT (v7.475)^79^. PAL2NAL (v14)^80^ was then used to convert the protein sequence alignments into the corresponding codon alignments and trimmed with trimAl (v1.4)^81^. The maximum likelihood (ML) concatenated tree was constructed using IQ-TREE (v2.0.3)^82^ with 1,000 replicates of the ultrafast bootstrap approximation^83^ for datasets including all three codon positions (CDS) and including only the first and second codon positions (Codon12) separately, and the best-fitting substitution model was automatically selected by internal program ModelFinder^84^ (-bb 1000 -m MFP). In addition, coalescent-based analysis was also used to infer phylogenetic relationship. Initially, un-rooted gene trees from each of the two datasets (CDS and conden12) were individually estimated using IQ-TREE, and then the quartet-based species tree was reconstructed based on these gene trees by ASTRAL (v5.6.1)^85^. To quantify phylogenetic discordance among loci, the gene concordance factor (gCF) and the site concordance factor (sCF)^86^ for every node in the species trees were calculated.

Given that the whole-genome sequences contain more genetic information than protein-coding genes, we further constructed a phylogenetic tree based on multiple whole-genome sequence alignment (MSA) that was generated across 19 poplar and three willow genome sequences using the progressive mode in Cactus^21^. To do so, the hierarchical alignment format (HAL) was converted into multiple alignment format (MAF) using the HAL tools command hal2maf^87^ with parameters “--noAncestors, --onlyOrthologs and --noDupes” to ignore paralogy edges. To reduce errors in phylogenetic inferences, only blocks longer than 100bp and containing at least one species in sect. *Populus*, sect. *Turanga*, ATL clade and willows were retained. All realigned individual segments were concatenated to obtain final whole genome alignments of 48.68 Mb and were then used to infer the topology by IQ-TREE with the ModelFinder function (-bb 1000 -m MFP). Moreover, low-copy genes (LCGs, n=11,385), which ranged between one and five gene copies per species in each orthologous group, were also employed to infer species trees by ASTRAL-pro^88^.

Divergence times among species were estimated using r8s (v1.81)^89^ software with selected parameter settings as follows: “blformat lengths=persite nsites=3597653 ulrametric=no; set smoothing=100; divtime method=PL algorithm=TN” and the others as defaults. The root age of the tree was calibrated to 48–52 Mya obtained from the TimeTree database (http://www.timetree.org/). Finally, with the gene family data and ultrametric phylogeny as input, CAFÉ (v4.2)^90^ was used to identify expansion and contraction of gene families, with the parameter p-value threshold was set to 0.01 and auto searching for the λ value. The lineage/species-specific gene families that had undergone expansion and contraction were further subjected to functional analysis using GO enrichment.

### Nucleotide diversity and demographic history analysis

For each species, whole-genome resequencing of two to three individuals with an average sequencing depth of 32× were obtained. After filtering the raw sequencing reads using Trimmomatic (v0.36)^91^, the clean reads were mapped to the respective reference genome with BWA-MEM (v0.7.17)^92^ and sorted with SAMtools (v1.9)^93^. The MarkDuplicates tool in Picard (v2.18.21, http://broadinstitute.github.io/picard/) was then used to remove duplicates. To estimate the nucleotide diversity (π) of different poplar species, variant calling for each individual was carried out using the HaplotypeCaller tool in GATK (v3.8.1)^94^ and the resulting gVCF files were merged to a single-variant calling file using CombineGVCFs and GenotypeGVCFs with the option “-all-sites”. The program pixy^95^ was then used to calculate the pairwise nucleotide diversity for each species.

To further investigate the demographic history of each species, the pairwise sequentially Markovian coalescence model (PSMC)^22^ was used to infer historical dynamics of effective population sizes (*N_e_*) with parameters -N25 -t15 -r5 -p “4+25*2+4+6”. Assuming 15 years as the generation time and a mutation rate of 3.75×10^-8^ per site per-generation^96^, we converted scaled population parameters into years and *N_e_*. Bootstrapping was performed 50 replicates per individual to examine the variance in *N_e_*.

### Transposable element analysis

To examine the role of transposons in driving genome evolution, we first calculated the distance between TEs and the nearest gene as well as the distance between genes and the nearest TE using the BEDTools (v 2.29.2)^97^ closest function. Furthermore, we compared the proportion of TEs in shared (homologous) vs. species-specific (non-homologous) sequences across genomes of various species. With the MAF files generated by multiple whole-genome sequence alignment, we identified species-specific or shared sequence between species and the sequence coordinates of each genome were saved as bed files using a custom Python script. Next, species-specific and shared sequences were intersected with TE annotations, respectively, using BEDTools intersect function.

For the insertion time (T) estimation of TEs, we extracted the full-length LTR-retrotransposons (fl-LTRs) and calculated the insertion time according to the formula T=*K*/2μ, where *K* is the sequence divergence rate between its 5’ and 3’ -LTRs and using the neutral mutation rate of μ = 2.5 × 10^−9^ mutations per bp per year.

To quantify transposon dynamics between species, the syntentic fl-LTRs among genomes of the 19 species were identified by sequence clustering of TE junctions from the annotated fl-LTR locations using the program Vmatch (http://www.vmatch.de) with the following parameters: -dbcluster 90 90 -identity 90 -exdrop 4 -seedlength 20 -d. The junctions consisted of 2×100-bp sequence signatures spanning the upstream and downstream insertion sites, with each 50 bp inside and 50 bp outside of the fl-LTR element.

### Construction of the super-pangenome

Ortholog groups among the 19 *Populus* genomes were identified using OrthoFinder (v2.5.2)^78^ with default parameters. To improve the accuracy of the analysis, genes with early stop codons or open reading frames shorter than 50 amino acids were removed and only the longest transcript of each gene was selected for gene family clustering. The resulting gene families were divided into core, softcore, dispensable compared protein coding length, distance to proximal upstream TEs, nucleotide diversity, *d*_N_/*d*_S,_ expression level and tissue specificity among different types of pan-genes. Besides, the epigenetic marks and transcriptional regulatory elements distributions were also compared by characterizing the patterns of DNA methylation and chromatin accessibility.

For *d*_N_/*d*_S_ analysis, only single-copy ortholog groups (scOGs) containing more than three species were used to avoid biases related to duplication among lineages and out-paralog genes. As described above, protein sequences of scOGs were first aligned by MAFFT and then converted to DNA codon alignments using PAL2NAL. Next, trimAl was used to trim the aligned CDSs. Maximum Likelihood trees for each of the scOGs were constructed using IQ-TREE (-m MFP -bb 1000) based on the trimmed alignments. The Codeml program of the PAML (v4.9i)^98^ was used to estimate the *d*_N_/*d*_S_ ratio for each ortholog group using its corresponding phylogenetic tree, with “model=0, NSsites=0, ncatG=1” choices.

### Expression analysis

For both the newly generated RNA-seq datasets and published RNA-seq datasets (Supplementary Table 1), Trimmomatic (v0.36)^91^ with default parameters was used to remove adapters and low-quality reads form the raw RNA-seq reads. The clean RNA-seq reads were mapped to the corresponding reference genome using Tophat (v2.1.1)^99^ with default settings (Supplementary Table 11). Gene expression abundance was estimated using FPKM (fragments per kilobase of exon per million fragments mapped) calculated by Cufflinks program (v2.2.1)^100^ for each transcript. Tissue-specific expression was assessed by the Tau index^101^, which is calculated as follows:

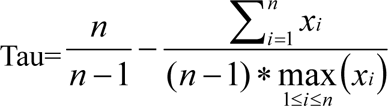

where *i* is a tissue, *xi* is the expression for tissue *i*, *n* is the total number of tissues.

### Whole-genome bisulfite sequencing and methylation calling

For whole-genome bisulfite sequencing (WGBS) of the 6 species sampled in the present study, genomic DNA of fresh leaf was isolated by DNeasy Plant Mini Kit (Qiagen) and sonicated into 200-300 bp using Covaris S220. Bisulfite treatment was performed using EZ DNA Methylation GoldTM Kit (Zymo Research) according to the manufacturer’s instructions. WGBS libraries were sequenced on the NovaSeq platform (Illumina) for 150 bp paired-end reads after evaluation by Agilent Bioanalyzer 2100. Together with the 7 published methylomic data (Supplementary Table 1), the raw sequencing reads of 13 species were trimmed using Trimmomatic (v0.36)^91^ with parameters “TruSeq3-PE.fa:2:30:10 LEADING:20 TRAILING:20 SLIDINGWINDOW:4:20 MINLEN:50”. The trimmed reads were then mapped to the respective reference genome using bowtie2 (v2.3.2)^60^ via Bismark (v0.22.3)^102^ with options “-N 1 -L 20”, followed by removal of PCR artifacts using deduplicate_bismark and only the unique mapped reads were retained for methylation calling using bismark_methylation_extractor (Supplementary Table 12). To reduce possible sequencing errors, methylated cytosine (mC) sites with less than four read coverages were finally discarded. The methylation level of each mC site was determined by the percentage of reads supporting mCs to all Cs at the site. The methylation level in genes bodies and the flanking regions was determined by evenly divided the region into 30 and 20 bins, respectively, and evaluated as a weighted methylation level.

### ATAC sequencing and accessible chromatin regions (ACRs) identification

Approximately 500mg of flash-frozen leaves were immediately chopped with approximately 1ml of prechilled lysis buffer (20mM Tris pH8.0, 40mM NaCl, 90mM KCl, 0.5 mM spermidine, 0.2 mM spermine, 5mM 2-mercaptoethanol and 0.1% Triton X-100). The suspension was then filtered through a series of cell strainers, and then the samples were disrupted through a Dounce homogenizer, and washed four times with ice-cold isolation buffer. In order to continue with 50,000 nuclei, the crude nuclei were stained with 4,6-Diamidino-2-Phenylindole (DAPI), and counted under a microscope using a hematocytometer. The sorted nuclei were incubated with 2μl Tn5 transposomes in 40μl tagmentation buffer (10mM TAPS-NaOH pH8.0, 5mM MgCl2) at 37 °C for 30min without rotation. The integration products were purified and transposed DNA using a Qiagen MinElute PCR Purification Kit according the manufacturer’s instructions. Then amplified using Phusion DNA polymerase for 10-13 cycles. Amplified libraries were purified using AMPure XP beads to remove primers. The purified libraries were stored at -20 ℃ for future use. Finally, NGS libraries were sequenced on an Illumina HiSeq platformm (Illumina, CA, USA) for twelve *Populus* species (Extended Data Fig. 5f and Supplementary Table 1). After sequencing, raw reads were trimmed using Trimmomatic (v0.36)^91^ with parameters “LEADING:20 TRAILING:20 SLIDINGWINDOW:4:20 MINLEN:36”. The trimmed reads were then aligned to their corresponding reference genomes using Bowtie (v2.3.2)^60^ with the following parameters: ‘bowtie2 --very-sensitive -N 1 -p 4 -X 2000 -q’. Aligned reads were sorted using SAMtools (v1.9)^93^ and clonal duplicates were removed using Picard (v2.18.21, http://broadinstitute.github.io/picard/). Finally, only high quality properly paired reads were retained for further analysis by SAMtools (view -b -h -f 3 -F 4 -F 8 -F 256 -F 1024 -F 2048 -F 1804 -q 30) (Supplementary Table 11). MACS2^103^ was used to define ACRs with the “--keep-dup all” function. The distribution of ACR peaks annotation for each species was visualized by ChIPseeker (v.1.28.3)^104^.

### Gene duplication classification and evolution analysis

DupGen_finder^31^ with default parameters was used to identify different modes of gene duplication: whole-genome duplication (WGD), tandem duplication (TD), proximal duplication (fewer than 10 gene distance on the same chromosome: PD), transposed duplication (TRD) and dispersed duplication (DSD). *S. suchowensis* was adopted as an outgroup to infer transposed duplicated gene pairs, all the poplars were used in a self-BLASTp search and a BLASTp search against *S. suchowensis*. Moreover, to eliminate redundant duplicate genes among different modes, we assigned each duplicated gene to a unique mode according to the priority: WGD >TD > PD > TRD > DSD. For each gene pair derived from different duplication modes, the *K_a_, K_s_*, and *K_a_/K_s_* values were calculated based on the YN model incorporated in KaKs_Calculator (v2.0)^105^. MAFFT and PAL2NAL were used for amino acid alignments and codon alignments conversion, respectively. Only gene pairs with *K_s_* values <5.0 were retained for further analysis. Genomic traits (CDS length, distance to proximal upstream TEs) and evolutionary parameters (*K_a_, K_s_*, and *K_a_/K_s_*) were further summarized and compared among genes originating from different modes of duplication.

We further explored the evolutionary fates of duplicate genes following WGD, and the duplicate gene pairs in which any partner was present in more than one WGD pairs was removed in view of the fact that only one recent whole-genome duplication event occurred in *Populus.* WGD gene pairs were then classified into three groups according to *K_s_* values. Briefly, all duplicates were first sorted in ascending order based on their *K_s_* values and then divided into three groups with an equal number of pairs by tertiles, defined as low *K_s_*, median *K_s_* pair and high *K_s_* groups. Gene pairs with no detectable expression level (FPKM=0) in both partners were further filtered out. For each duplicate pair, gene expression (E) and methylation (M) divergence between the two partners were calculated as (E1-E2)/(E1+E2) and (M1-M2)/(M1+M2), respectively, as described previously^106^. To further investigate the association between expression divergence and methylation divergence of duplicated genes, we also sorted all gene pairs in ascending order by their expression divergence and then divided them equally into 15 groups. We examined the correlations between expression divergence and methylation divergence (CG, CHG and CHH) both in genic and flanking regions (2 kb of upstream or downstream region) based on these 15 groups.

To detect the lineage and/or species-specific divergent clusters of genes retained following WGD, we first corrected the *K_s_* values of the above one-to-one WGD pairs using a method reported previously^107^ in order to avoid the resulting errors caused by inconsistent evolution rates among different poplar species. Then, hierarchal clustering (Euclidean’s distances and complete-linkage clustering) based on the corrected-*K_s_* values was constucted based on pheatmap (v1.0.12) for these one-to-one WGD pairs, in which both partners were retained in all poplar genomes (i.e., both belonging to core genes).

Furthermore, we performed the following steps for each genome to further control for the evolutionary age effects: (1) randomly sampling the duplicated pairs 1,000 times; (2) estimating the peak value of the corrected-*K_s_* distribution for each of the 1,000 samples, and then calculating the average and 95% confidence interval of all 1,000 corrected-*K_s_* peak values; (3) retaining the gene pairs with the corrected-*K_s_* value that were within the confidence interval. Finally, we obtained an average of 346 gene pairs (ranged from 232 in *P. alba var. pyramidalis* to 448 in *P. koreana*) per genome. According to the expression difference between duplicated genes, we further classified these pairs into conserved and divergent. The expression difference was calculated by the formula: diff_FPKM_ = (FPKM_High_ - FPKM_Low_)/ FPKM_Low_, where FPKM_High_ and FPKM_Low_ denote the genes with relative higher expression and lower expression in the duplicated pairs, respectively. The gene pairs with diff_FPKM_ ≥ 2 were assumed to be divergent, whereas those ≤ 0.5 were considered to be conserved. The CDS length, distance to closest upstream TE, expression level, tissue specificity and epigenetic regulation (methylation levels and ACRs distributions) were compared between the gene partners among the conserved and divergent pairs.

### SV detection and validation

To overcome the results bias caused by a single reference genome, we used two reference genomes (*P. trichocarpa* and *P. adenopoda*) to detect structural variations (SVs). All the *Populus* assemblies were aligned to the reference genome using nucmer in the MUMmer (v4)^108^ package with parameters “-l 40 -g 90 -b 200 -c 100 -maxmatch”, and the resulting alignments were further filtered using delta-filter with parameters “-m -i 90 -l 100”. Then, SyRI pipeline^38^ was employed to detect structural variations with default parameters and five types of representative SVs (>50 bp) were extracted for further analysis, including insertions (INS), deletions (DEL), inversions (INV), translocations (TRANS/INVTR) and duplications (DUP/INVDP).

A total of 50 SVs including insertions and deletions were randomly selected from SVs between the *P. trichocarpa* and *P. adenopoda*, and/or between the *P. pseudoglauca* and *P. adenopoda* assemblies, respectively. The raw long-reads from *P. adenopoda* were aligned to the *P. trichocarpa* genome using minimap2 (v2.17)^109^ to verify the DEL SV (relative to *P. trichocarpa*) in the *P. adenopoda* assembly. Similarly, the reads from *P. pseudoglauca* were aligned to the *P. adenopoda* genome to verify the DEL (relative to *P. adenopoda*) in *P. pseudoglauca*. On the other hand, the raw long-reads from *P. adenopoda* were aligned to the *P. pseudoglauca* genome to verify the INS SV (relative to *P. adenopoda*) in the *P. pseudoglauca* assembly. Lastly, the alignments were manually inspected using the Integrative Genomics Viewer^110^.

### Identification of hemizygous genes

To identify hemizygous genes, Nanopore long reads were mapped onto respective reference assembly using minimap2 (v2.17)^109^, and variant calling was then performed with Sniffles (v2.0.7)^111^. As the aim was to identify regions of the genome assemblies that were both hemizygous and contained previously annotated genes, the type of structural variants used for subsequent analyses were limited to deletions and the genotypes of ‘0/1’ which represented hemizygosity were extracted. Genes were extracted from hemizygous regions of the genome using BEDTools intersect, and only those genes falling fully within hemizygous regions were considered hemizygous genes and were extracted for analysis (using the -F 1 option).

### Characteristic analysis of SVs and hemizygous genes

The continuous or overlapping SVs of identical type were merged as a single SV. Further, we calculated the distribution of SVs for each 200 kb sliding window with a 100 kb step-size along the entire genome. The windows with number of SVs in the top 5% were defined as SV hotspots, and the continuous windows were merged into one region. First, we compared the ratio of SVs in putative disease resistance genes that contain at least an NB-ARC PFAM protein domain (PF00931) predicted using Pfam HMMs (InterProScan, v5.32-71.0) with other genes. Then, to assess hemizygous genes enrichment in SV hotspots, random genomic regions with the same length distribution to hotspots were generated as a control using the ‘shuffle’ command in BEDTools, and the ratio of hemizygous gene of these two sets (SV hotspot regions and random regions) was compared by using the intersectBed utility from BEDTools. Furthermore, we identified the genes overlapping with SVs (SV+) and with hemizygous states respectively. We further compared their ratios and enrichment in different categories of pan-genes and duplicated genes through comparing with random sets of genes (Generate 1000 samples, each time randomly selecting genes with the same number of SV+ genes and hemizygous genes). Lastly, we compared selection pressure (*K_a_, K_s_*, and *K_a_/K_s_*), expression levels, methylation levels and chromatin accessibility among hemizygous genes, genes with (SV+) and without SVs (SV-).

### Analysis and RT-qPCR for the *CUC2* gene

To quantify collinearity along the inverted region that contained *CUC2* gene, we measured and compared the synteny diversity across all pairwise genome comparisons within Clade-Ⅰ and Clade-Ⅱ separately as well as between these two clades, following the method reported by^112^. For validation of *CUC2*-associated inversion, the long reads of query genome were mapped to the reference genome (*P. trichocarpa*) using minimap2 (v2.17)^109^ and further visualized through samplot (v1.3.0)^113^ Motif calling was analyzed on the local region of insertion sequence of *CUC2* promoter using PlantCARE^114^.

Reverse transcription quantitative PCR (RT-qPCR) was used to investigate the expression levels of *CUC2* genes in species from different sections. Total RNA was extracted from the leaf margin areas of the emerging young leaves at the same developmental stage containing only serrations, and the HiScript® III RT SuperMix for qPCR (+gDNA wiper) (Vazyme #R323) was used to obtain cDNA. qPCR was performed with gene-specific primers using the Taq Pro Universal SYBR qPCR Master Mix (Vazyme #Q712) reaction system on the CFX96 Real-Time detection system (Bio-Rad). Each experiment was performed with three reaction replicates and UBQ10 was used as the internal control for data analysis. Primer sequences are available in Supplementary Table 23.

### Dual-luciferase assays

Three species (*P. koreana, P. simonii* and *P. lasiocarpa;* absence of the ∼180bp upstream of *CUC2*) in Clade-Ⅰ and three species (*P. alba var. pyramidalis, P. adenopoda* and *P. qiongdaoensis*; presence of the ∼180bp upstream of *CUC2*) in Clade-Ⅱ were selected for experiment and the dual-luciferases (LUC) assay was performed using young *N. benthamiana* leaves^115^. For these six species, 1.2-1.4 kbp of the *CUC2* promoter containing the entire SV were isolated from the genomic DNA with specific primers (Supplementary Table 24 and Supplementary Fig. 39) followed by a 35S mini promoter and ligated with the BamHI-digested pGreen II 0800-LUC vector as reporters, respectively. To further validate the regulatory potential of this SV, we artificially designed a deletion in three species belonging to Clade-Ⅱ, i.e., the corresponding upstream and downstream sequences of the variant were extracted and then connected using ClonExpress MultiS One Step Cloning Kit (Vazyme Code: C112-02). These connected sequences were inserted into the BamHI-digested pGreen II 0800-LUC vector to use as reporters, too. Equal amounts of *A. tumefaciens* strain GV3101 carrying different constructs were injected into different regions of 3-week-old *N. benthamiana* leaves. Luciferase and renilla activities were measured with a dual-LUC Reporter Assay System (Promega, Madison, WI, USA) after 2 days of incubation in the dark and 1 day of cultivation under normal growth conditions. Each experiment was independently performed three times. In addition, the injected leaves were sprayed with sodium luciferin (Gold Biotech, China) and the luciferase luminescence from the infiltrated area was imaged using PerkinElmer IVIS Lumina III (PerkinElmer, USA).

### Data availability

All raw sequencing data as well as genome assembly and annotation generated in this study have been submitted to the National Genomics Data Center (NGDC; https://bigd.big.ac.cn/bioproject) under BioProject accession number PRJCA010101 and will be available upon publication. All scripts used in this study will be available at https://github.com/jingwanglab/Populus_superpangenome upon publication.

## Extended Data Figures

**Extended Data Fig. 1.**
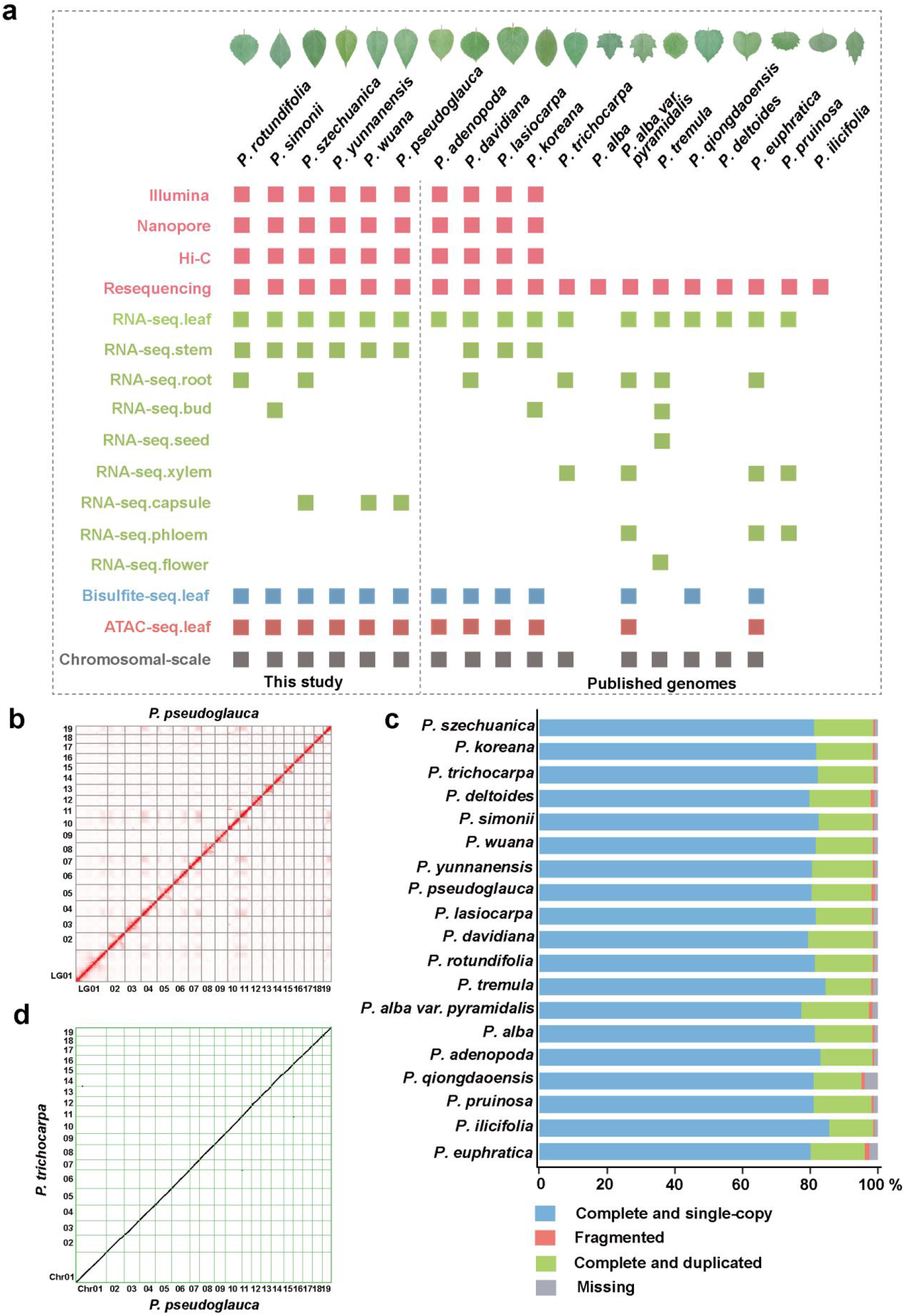
General information about the poplar species and characteristics of genome assemblies in this study. **a,** General information about the 19 genomes used in this study. The colored rectangles represent the corresponding data available for each dataset. Detailed data sources are listed in the Supplementary Table 1. **b,** Hi-C heatmap of *P. pseudoglauce* shown with a resolution window of 100 kbp. Darker red indicates stronger interactions. **c,** BUSCO assessment for the poplar genome assemblies. **d,** Synteny between *P. trichocarpa* and *P. pseudoglauce*. The scaffolds nomenclature was adopted for the chromosome numbering on the basis of their collinearity with 19 chromosomes of *P. trichocarpa* genome.

**Extended Data Fig. 2.**
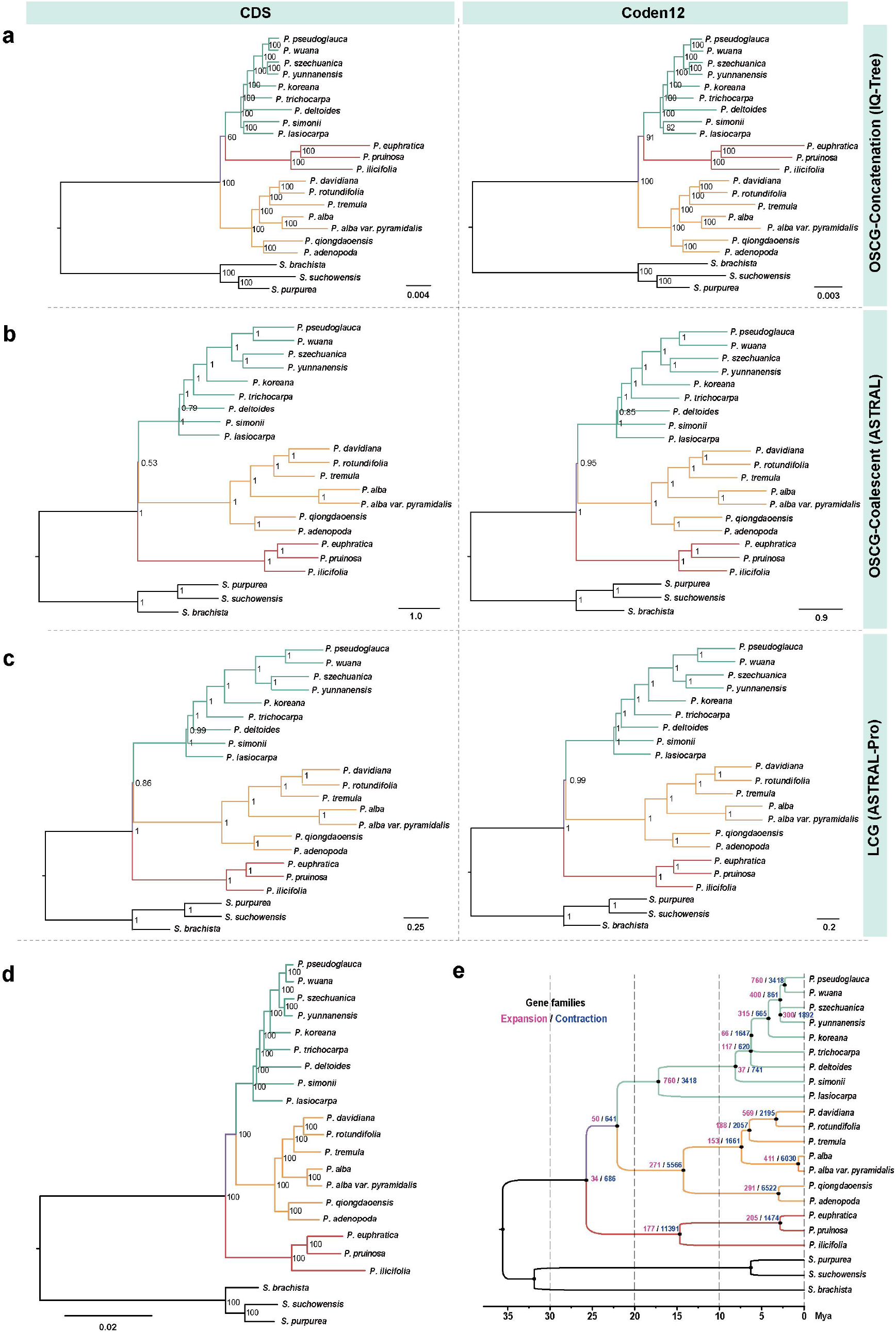
Phylogeny of the genus *Populus*. **a,b,c,** Phylogenetic relationships inferred with the 2,455 single-copy genes (CDS and conden12 respectively) using IQ-TREE (a) and ASTRAL (b), 11,385 low-copy genes (CDS and conden12 respectively) using ASTRAL-Pro (c). **d,** Phylogenetic relationships inferred using the orthologous regions (∼48.7Mb) from the multiple sequence alignments (generated by Cactus) using IQ-TREE. **e,** Estimation of divergence time and dynamic evolution of orthologous gene families. Gene family expansion events are shown in pink and gene family contraction events in blue. The calibration time for divergence between *P. trichocarpa* and *S. suchowensis* (12-48 Mya) was obtained from the TimeTree database (http://www.timetree.org/).

**Extended Data Fig. 3.**
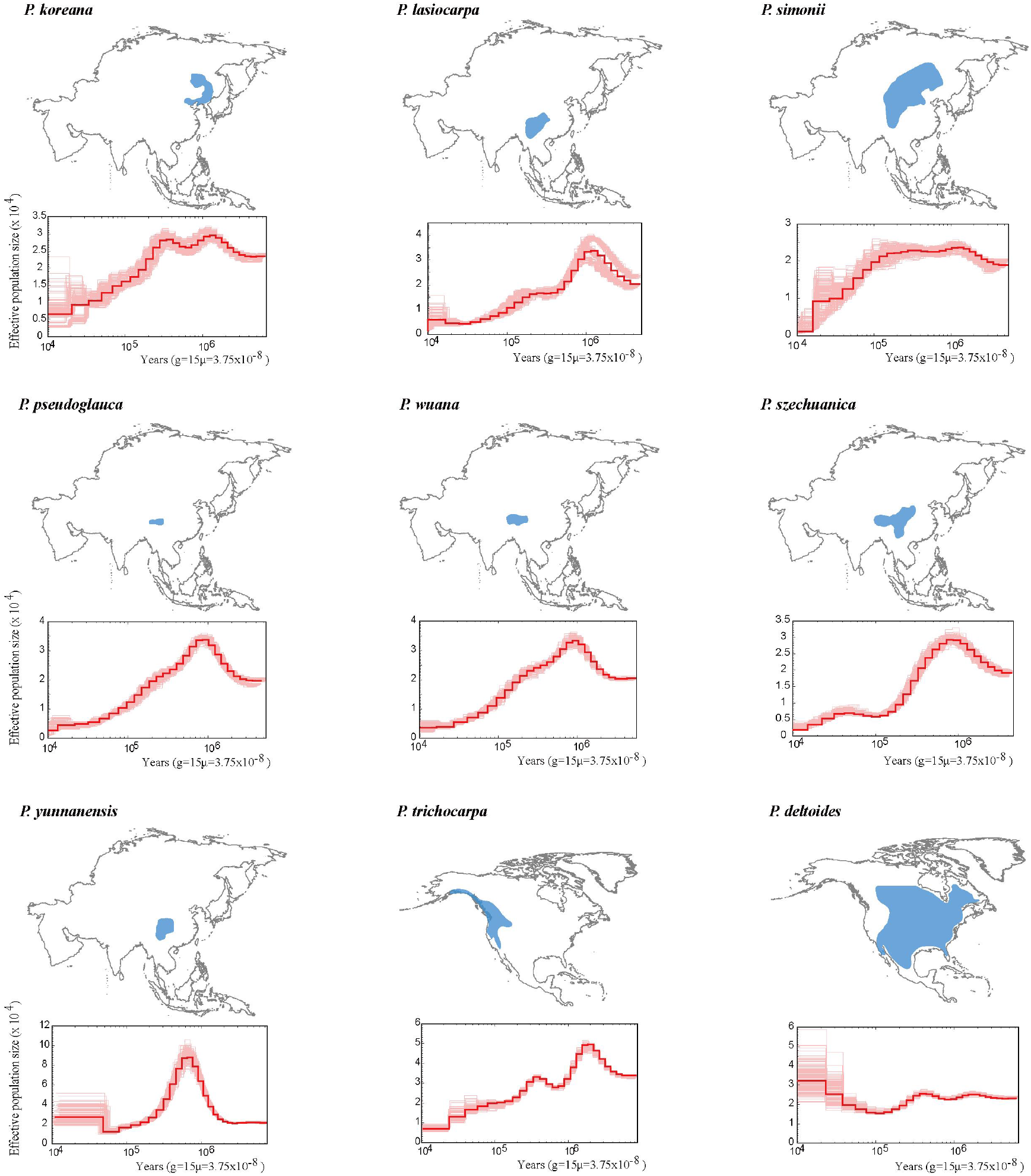
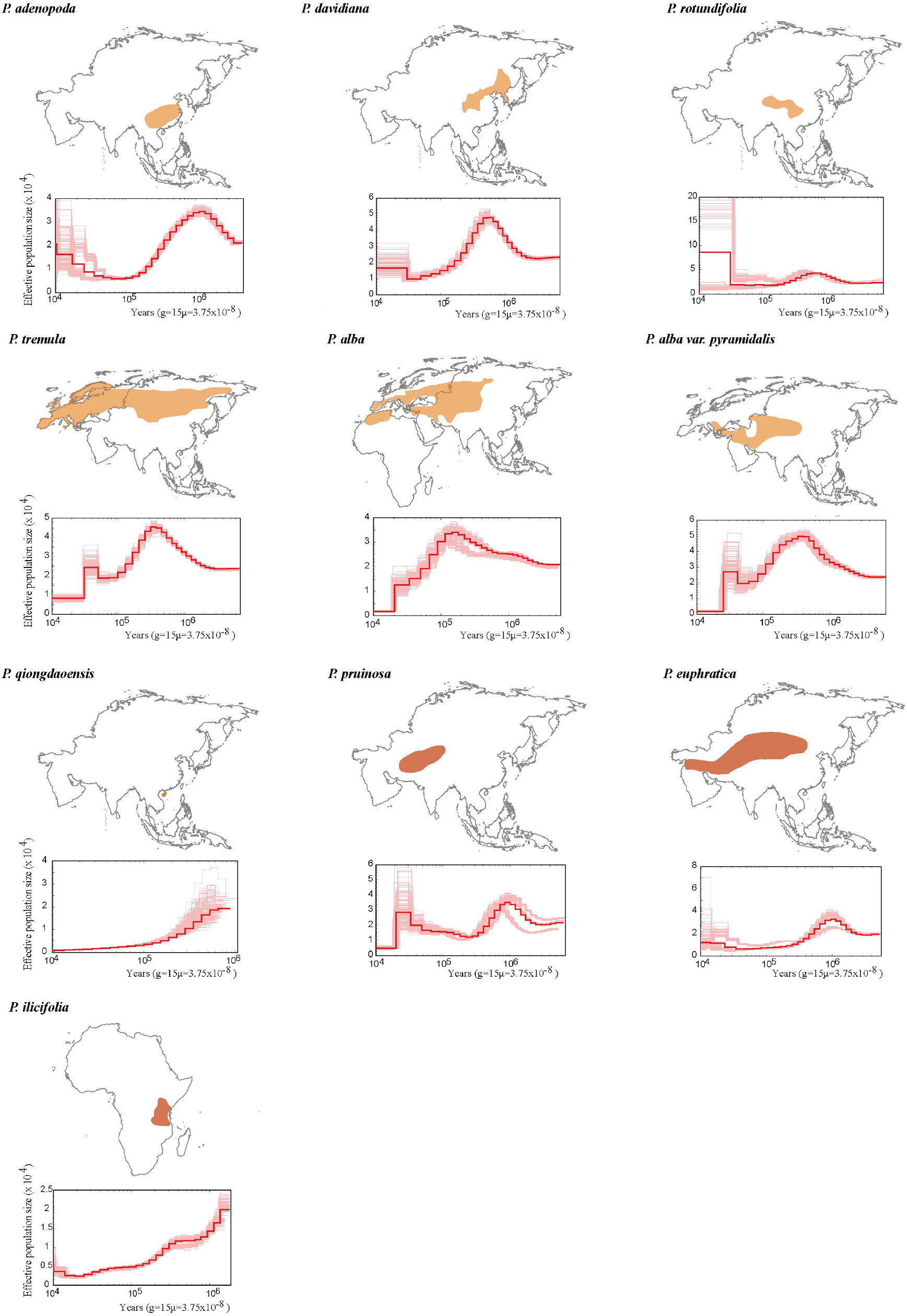
Demographic history of different poplars. Changes in effective population size (*Ne*) through time inferred by the Pairwise Sequentially Markovian Coalescent model (PSMC). Bold lines are the mean estimate for all resequenced samples, whereas faint lines are for 50 replicates per individual.

**Extended Data Fig. 4.**
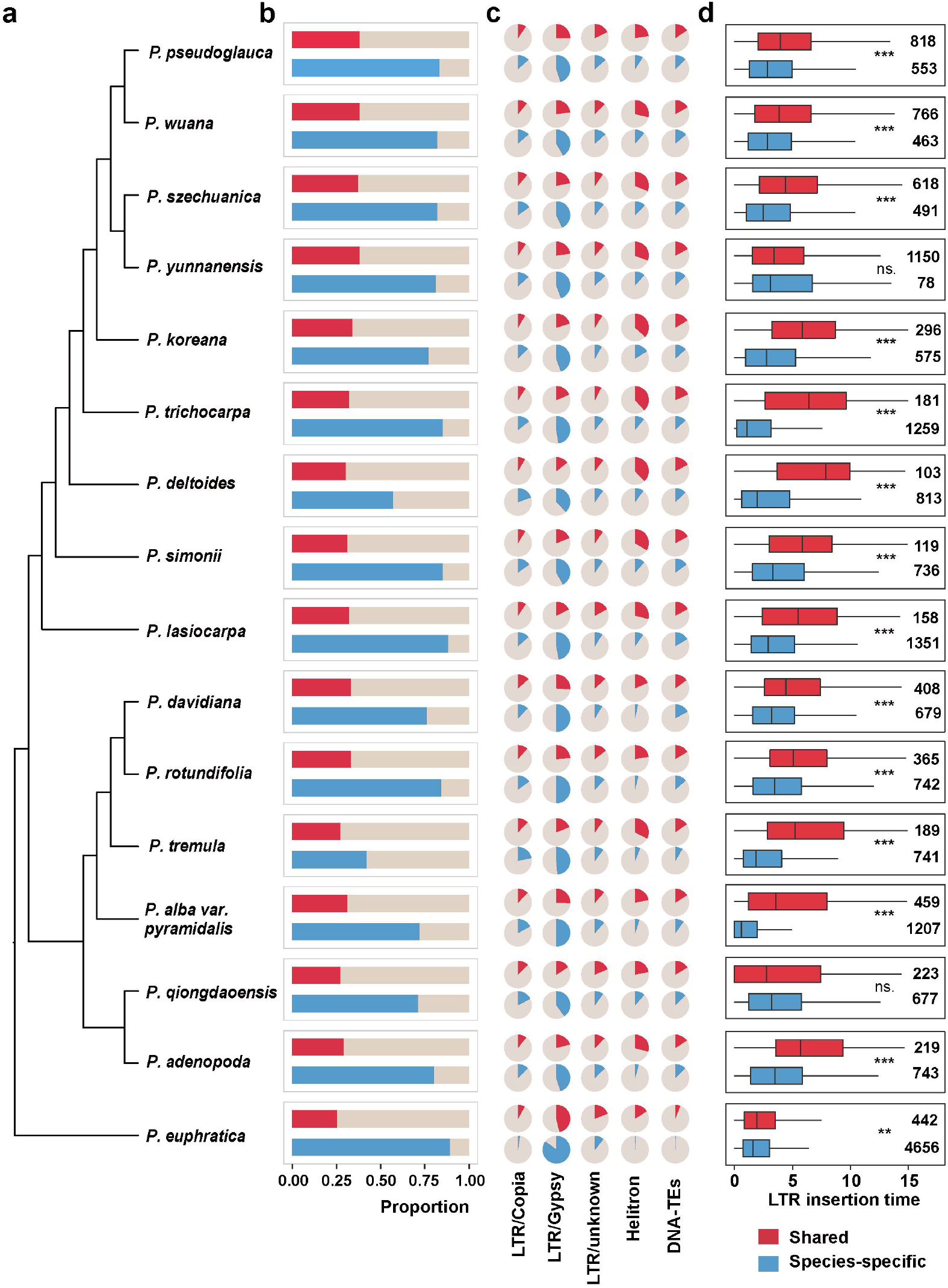
TE dynamics in the shared and species-specific genomic regions across the 19 poplar species/sub-species. **a,** Phylogenetic relationship between the poplar species. **b,** TE proportion in the shared and species-specific genomic regions, respectively. The shared and species-specific genomic regions were extracted from the multiple sequence alignments generated by Cactus. **c,** Proportion of each TE superfamily in shared and species-specific genomic regions. **d,** Age distribution of LTR-RT insertions belonging to shared and species-specific genomic regions. The numbers indicate the sample size used in the analysis. The statistical difference between groups was calculated using Wilcoxon ranked sum tests: ns. *P* > 0.05; \**P* ≤ 0.05; \*\**P* ≤ 0.01; \*\*\**P* ≤ 0.001.

**Extended Data Fig. 5.**
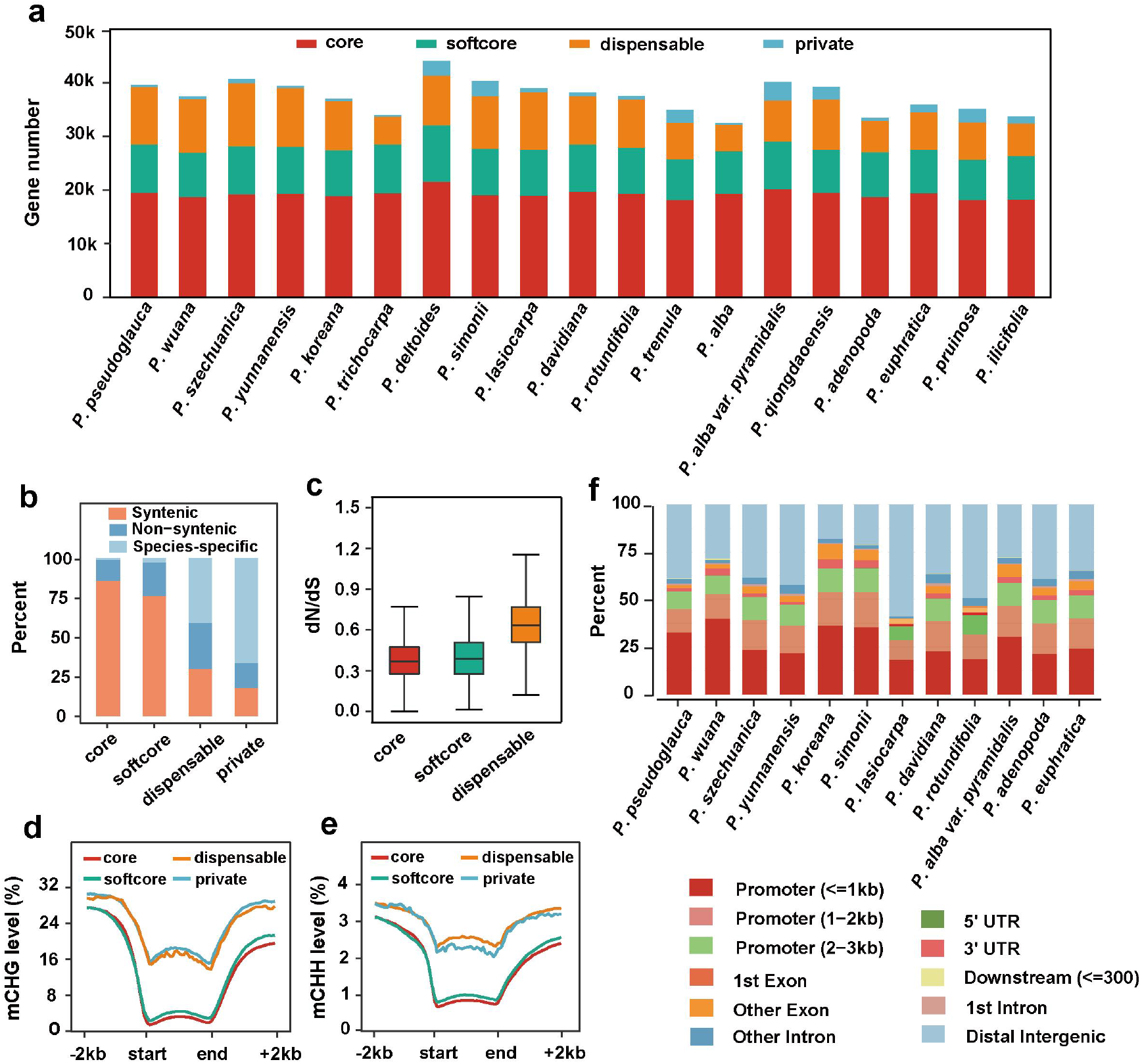
Pan-genome analysis of the gene space. **a,** Number of genes in the core, softcore, dispensable, and private fractions in each genome. **b,** Overall proportions of syntenic, non-syntenic, and species-specific loci in each genome based on their pan-genome classification. Syntenic genes in each genome were calculated using *S. suchowensis* as the query genome with MCScanX. **c,** The ratio of non-synonymous to synonymous mutations (*dN/dS*) in core, softcore and dispensable genes. Only single-copy ortholog groups that contained more than three species were used to calculate dN/dS ratio using PAML. **d,e,** Differences in average CHG (d), and CHH (e) methylation level along the gene and flanking regions among core, softcore, dispensable and private genes. The color of the line is consistent with the classification of (a). **f,** Genome-wide distribution of annotated ATAC-seq peaks in 12 poplar species.

**Extended Data Fig. 6.**
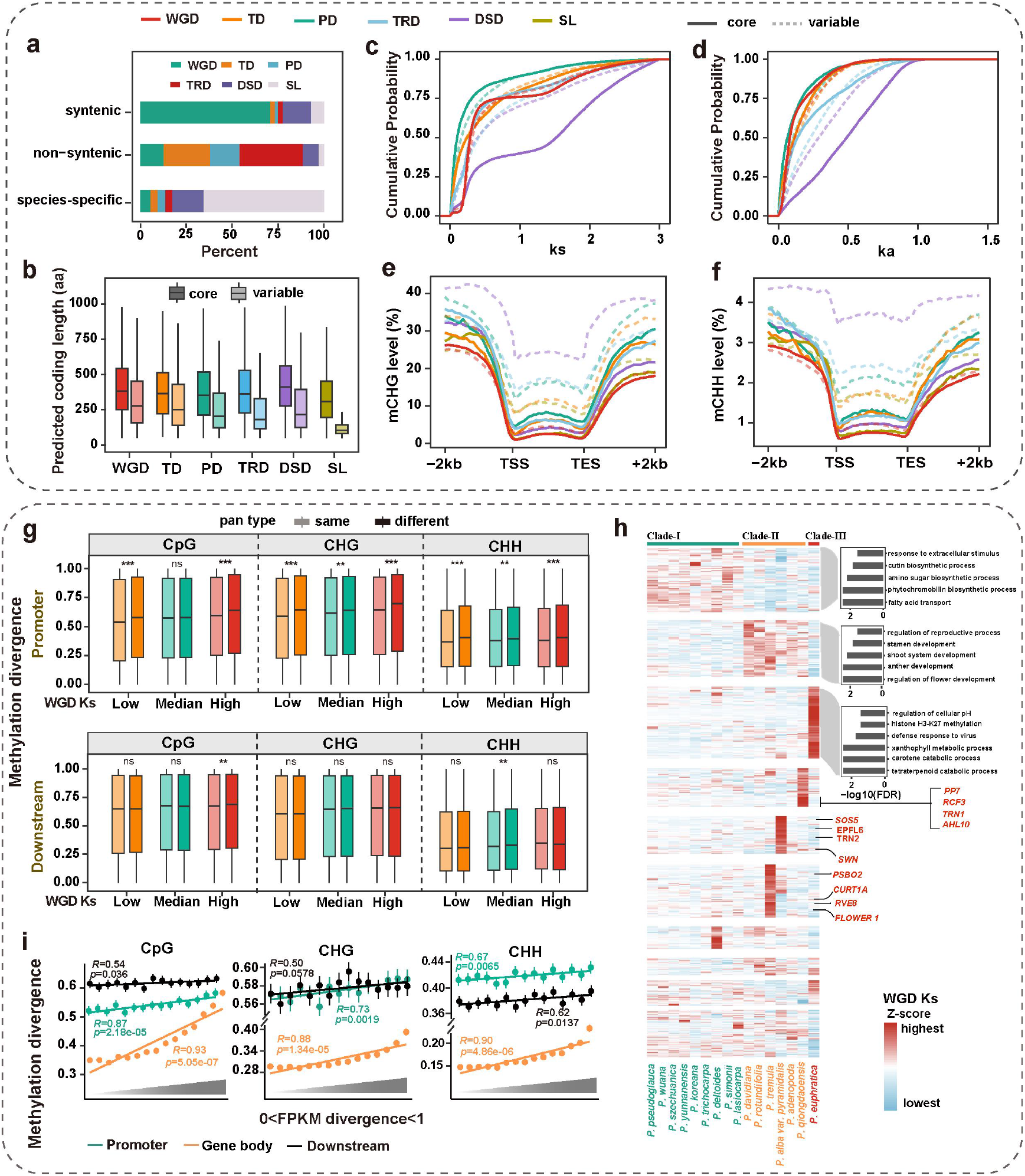
The landscape of gene duplication and the evolutionary fates of WGD-derived genes over a long evolutionary time period in poplar genomes. **a,** Overall proportions of WGD, TD, PD, TRD, DSD and singleton loci based on their syntenic blocks with *S. suchowensis*. Syntenic genes in each genome were calculated using MCScanX. **b,** CDS length of different duplicated genes that in distinct pan-gene categories. Variable represent softcore, dispensable and private genes, which correspond those shown in Fig.3. **c,d,** The *K*s (c) and *Ka* (d) distributions of different duplicated gene pairs based on different pan-gene categories**. e,f,** Differences in average CHG (e) and CHH (f) methylation level along the gene and flanking regions among different gene groups based on the duplication status and pan-gene type. **g**, methylation divergence (upstream and downstream) of duplicated genes with evolutionary age and the consistency pan-gene status between paralogs. **h**, Clade- and/or species-specific functional divergence of duplicated genes retained following WGD. Unsupervised hierarchical clustering of corrected-*K*s values (Z-score) of WGD gene pairs belonging to core gene families (both paralogs retained in all species). Right: GO enrichment and gene examples for the WGD-derived genes from the clade-specific and species-specific functional divergence cluster, respectively. **i**, Pearson’s correlation coefficient between the expression divergence and methylation divergence of WGD gene pairs at CG, CHG, and CHH contexts. The different colors indicate the methylation divergence of duplicated genes in the promoter, gene body, and downstream regions, respectively. WGD whole-genome duplication, TD tandem duplication, PD proximal duplication, TRD transposed duplication, DSD dispersed duplication, SL singletons. The statistical analysis was performed using Wilcoxon ranked sum tests: ns. *P* > 0.05; \**P* ≤ 0.05; \*\**P* ≤ 0.01; \*\*\**P* ≤ 0.001. Results for each species are shown in the Supplementary Figs. 16-29.

**Extended Data Fig. 7.**
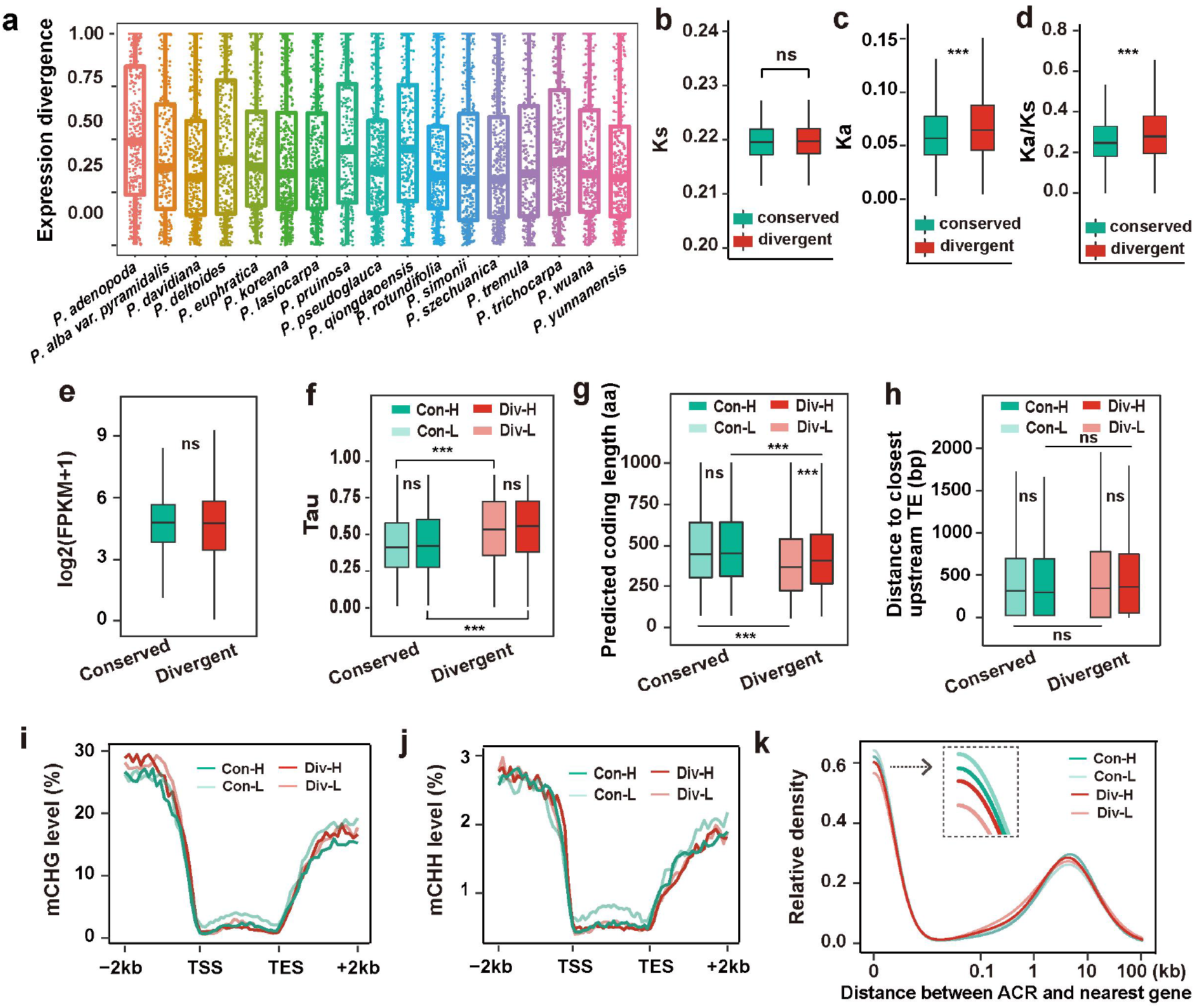
The diverse fates of WGD-derived genes under the same evolutionary age effects. **a,** Expression divergences of WGD-derived duplicates under the same sequence divergence. **b-d,** *Ks* (b), *Ka* (c) and *Ka*/*K*s (d) distributions of divergent and conserved gene pairs, as classified shown in Fig.4. **e,** Overall expression levels (log2 FPKM in leaf tissue) of conserved and divergent duplicate genes. **f,** Tissue specificity index (Tau index, at least three tissues) in divergent and conserved gene pairs. The legends “Con-L”and “Con-H” respectively represent genes with relative lower and higher expression in the conserved gene pairs, while “Div-L”and “Div-H” respectively represent genes with relative lower and higher expression in the divergent gene pairs (same in g-k). **g-k,** The CDS length (g), distance to neighboring TE (h), methylation in the CHG (i) and CHH (j) sequence contexts and the frequency distribution of ACRs and their distance to the nearest genes (k) for each of the two partners (divided into low and high according their expression) in conserved and divergent gene pairs. The statistical analysis was performed using Wilcoxon ranked sum tests: ns. *P* > 0.05; \*\*\**P* ≤ 0.001.

**Extended Data Fig. 8.**
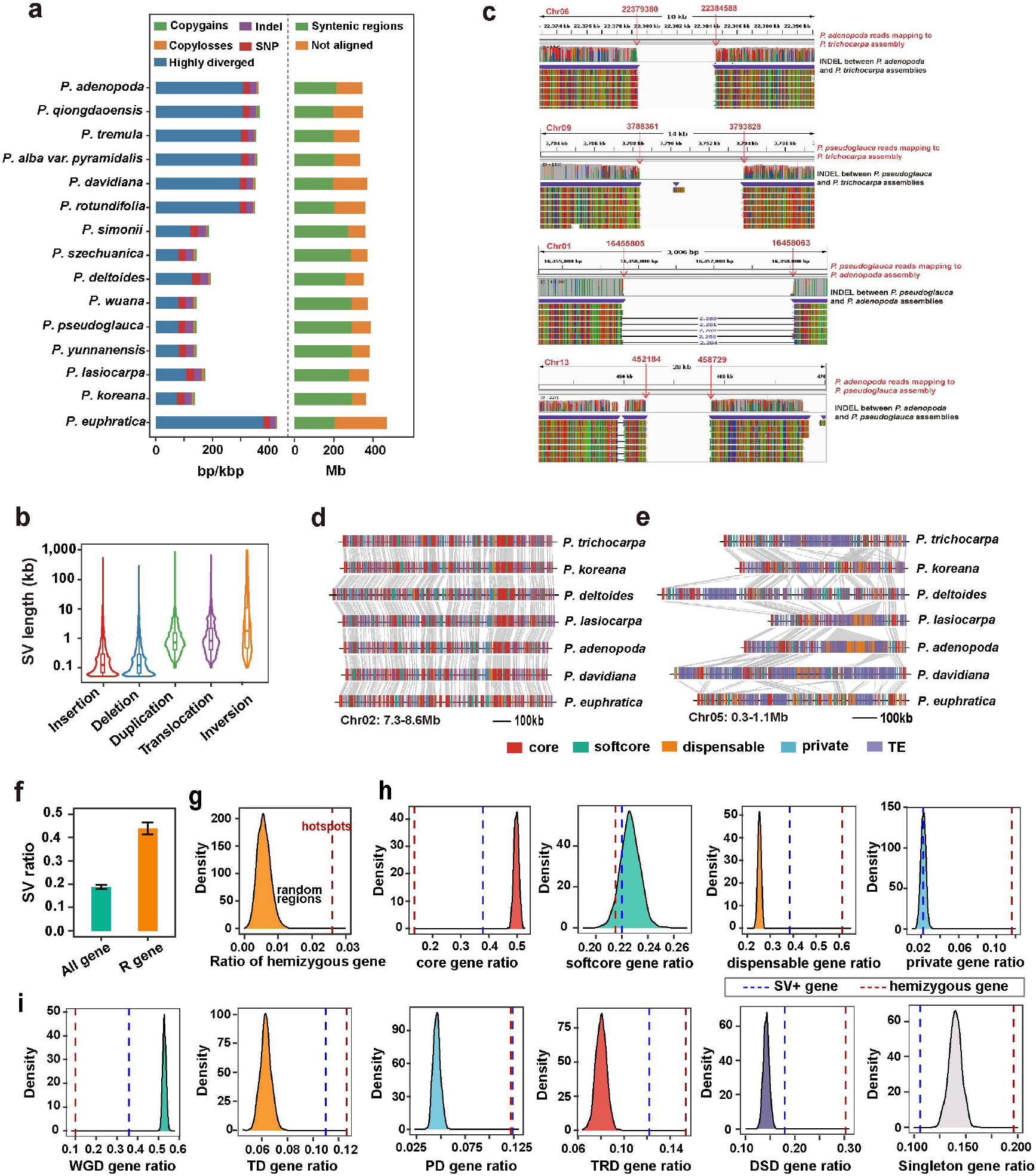
SVs landscapes in the genus *Populus*. **a,** Schematic of sequence variation detected in poplar genomes. **b,** Size distributions of different types of SV. **c,** Example manual validations of 50 randomly selected SVs (also see Supplementary Table 18), based on mapping Nanopore long-reads to the genome assemblies. **,d, e,** Examples of pan-genes and TEs enriched in SV hotspots (d) and desert (e) regions. **f,** Comparision of the ratio of SVs overlapped with NLR genes and all other genes. **g,** The density of ratios of hemizyous genes in SV hotspot region (dashed lines indicate the empirical observation) compared with genomic random regions with same sizes resulting from the 1,000 randomizations. **h,i,** Comparison of ratios of different categoreis of pan-gene (h) and duplicated gene (i) in SV-related (SV+) genes (blue dashed lines), hemizyous genes (red dashed lines) and genes selected randomly.

**Extended Data Fig. 9.**
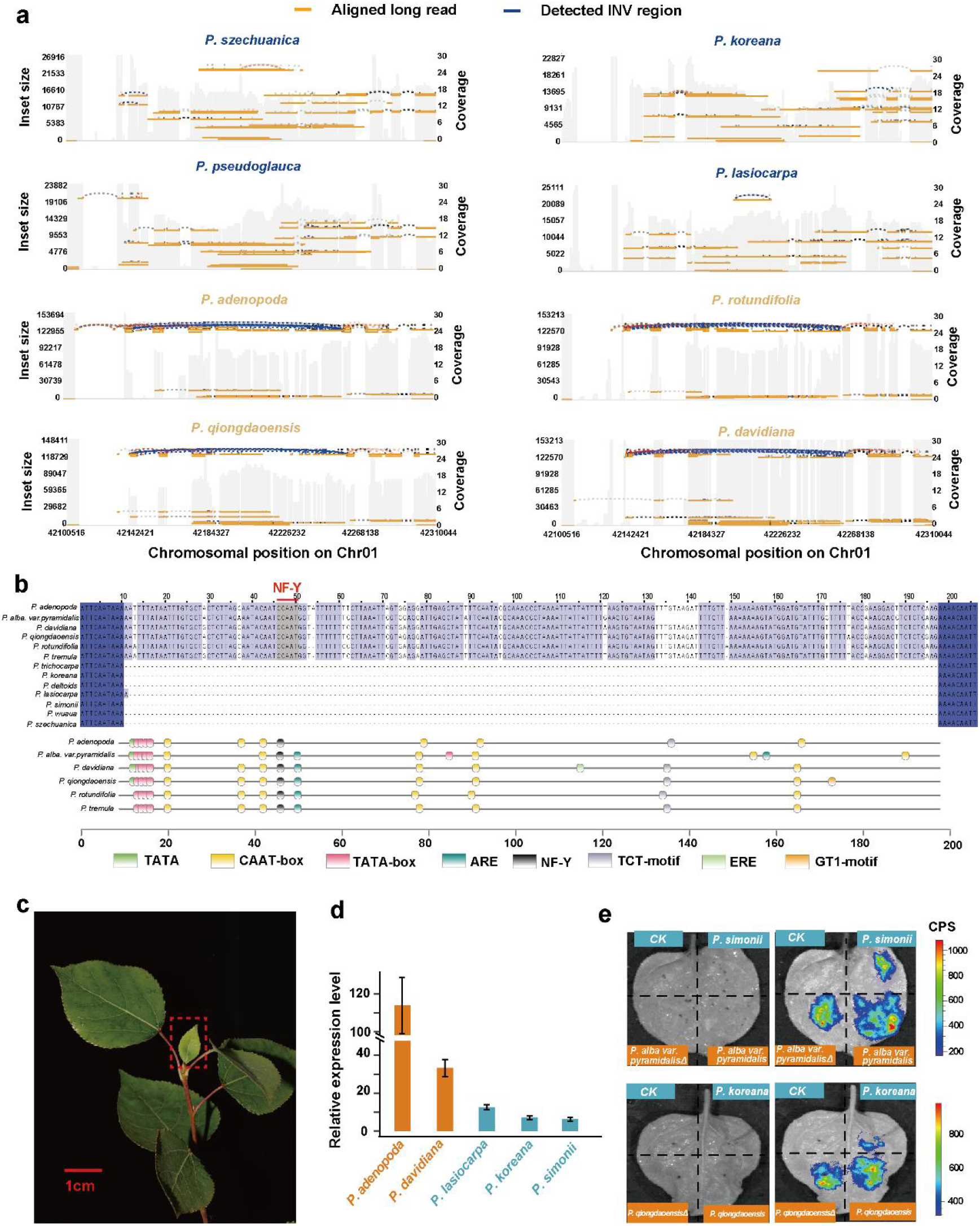
An inversion-mediated gene regulatory divergence that likely results in divergent patterning of the leaf margin between two clades of species. **a,** Inversion (Chr01: 42.15-42.26 Mb) validation by mapping the Nanopore long reads of the representative species from the two clades represented by Clade-Ⅰ (mostly belonging to sect. *Tacamahaca*) and Clade-Ⅱ (belonging to sect. *Populus*) genomes. Reads plotted in blue have large insert sizes and represent the detected inversion region. **b,** Predicted transcriptional factor binding motifs within the ∼180bp insertion located in *CUC2* promoter of multiple speceis from Clade-Ⅱ. **c,** Total RNA were extracted from young leaves, i.e., the leaf margin of the emerging young leaves in the terminal bud (shown in red box). **d**, *CUC2* expression levels (mean ± SD, n = 3) in species from the two clades that were determined by qPCR. **e,** Transient expression assay of luminescence intensity show that the deletion directly decreased the regulatory potential of the promoter on the *CUC2* gene. Representative images of *N. benthamiana* leaves 72 h after infiltration were shown.

