## Supplementary Figures for "The super-pangenome of *Populus* unveil genomic facets for adaptation and diversification in widespread forest trees"

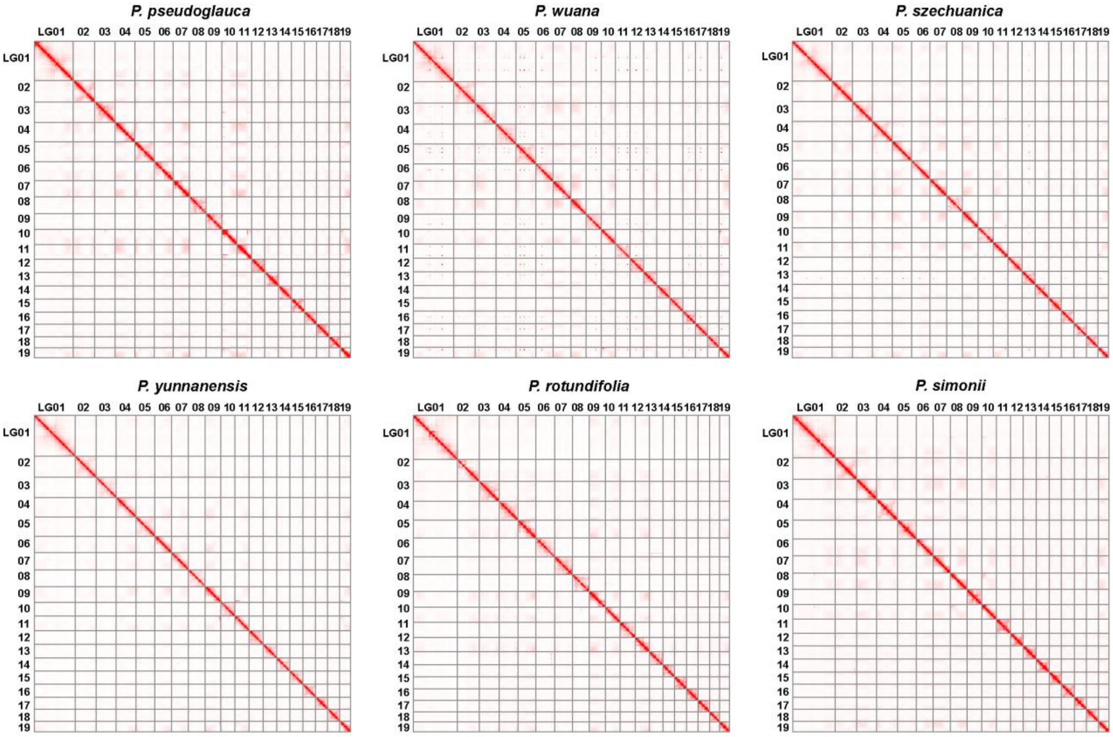


**Supplementary Fig. 1. Hi-C contact maps for 6 newly *de novo* assembled genomes.** Genome-wide interaction heat map of Hi-C links among chromosome groups (19 chromosomes). Each chromosome has higher intensity of interactions with itself than any other chromosomes (Darker red color means stronger interactions). The Hi-C heatmaps are shown at a resolution of 100 kb window.

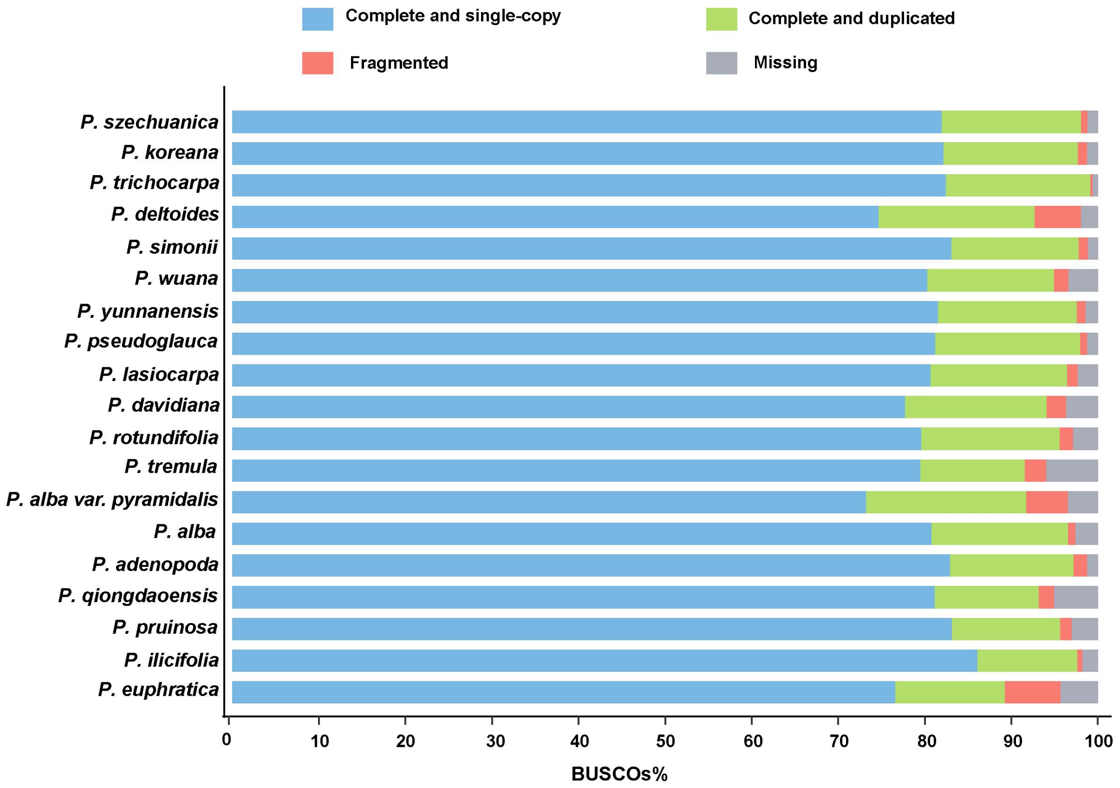


**Supplementary Fig. 2.** **BUSCO assessment of genome annotation quality.**

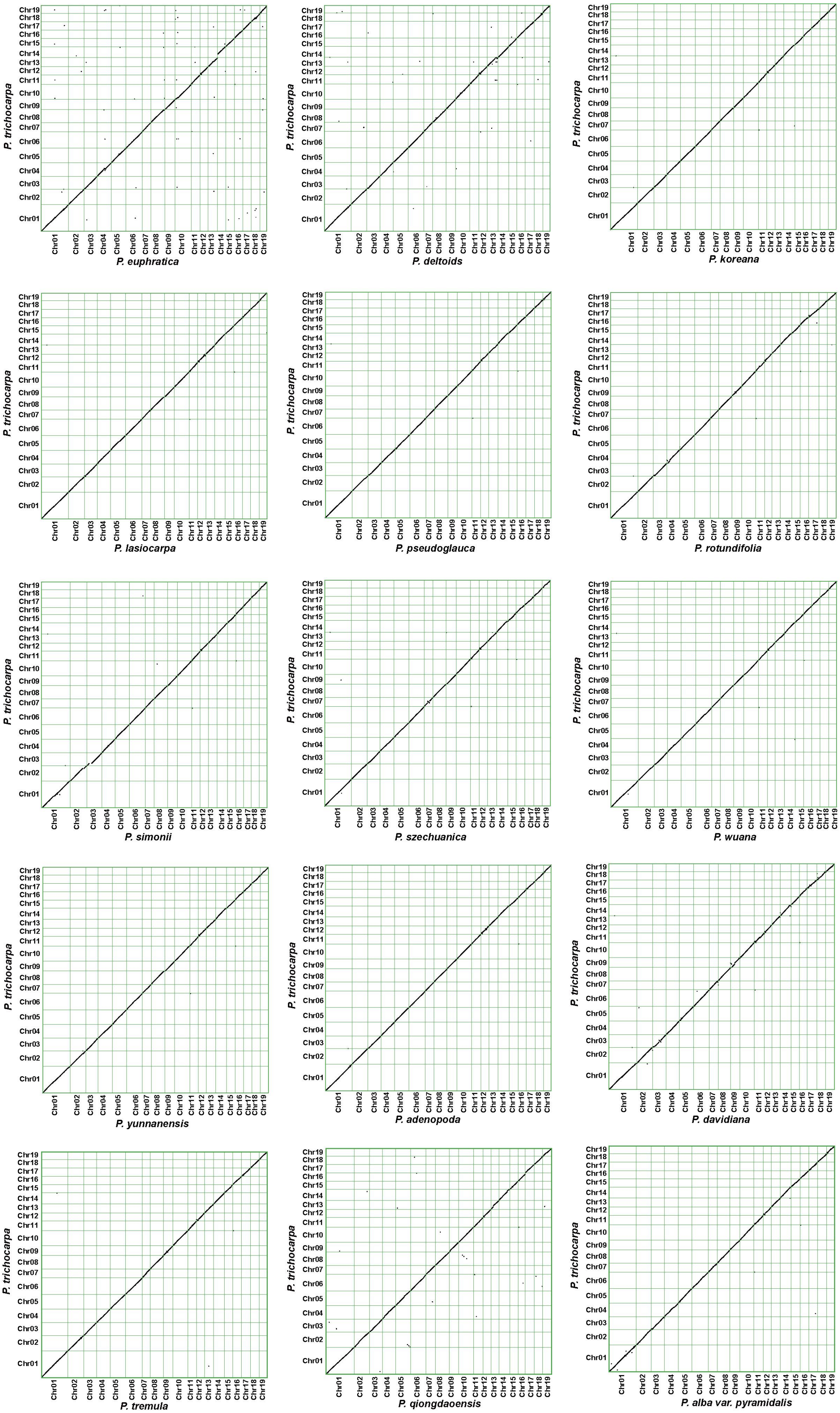
P. trichocarpa
P. euphratica
P. deltoids
P. koreana
P. lasiocarpa
P. pseudoglauca
P. rotundifolia
P. simonii
P. szechuanica
P. wuana
P. yunnanensis
P. adenopoda
P. davidiana
P. tremula
P. qiongdaoensis
P. alba var. pyramidalis

**Supplementary Fig. 3. Synteny between *P. trichocarpa* and the other poplar assemblies at the chromosome scale.** Synteny was assessed using the MCScanX program to identify collinear blocks of syntenic gene pairs.

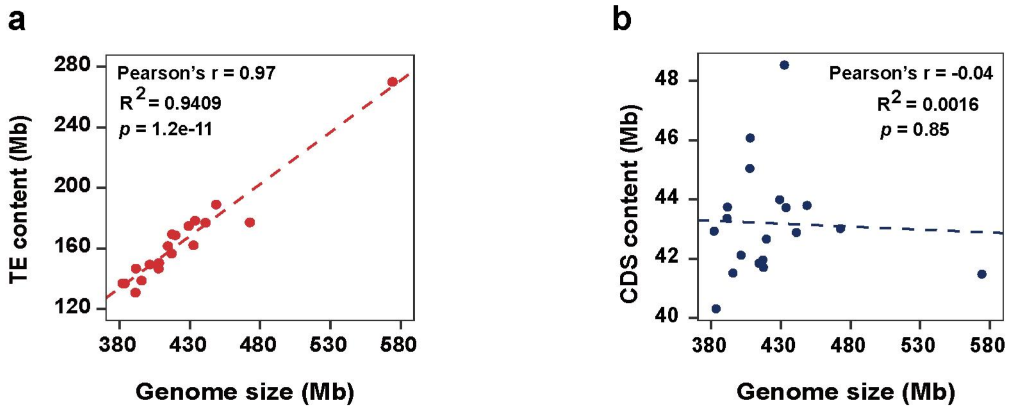


**Supplementary Fig. 4. Contribution of transposable element (a) and coding DNA sequence (b) to genome size variation across poplars.** Each dot represents a genomic assembly.

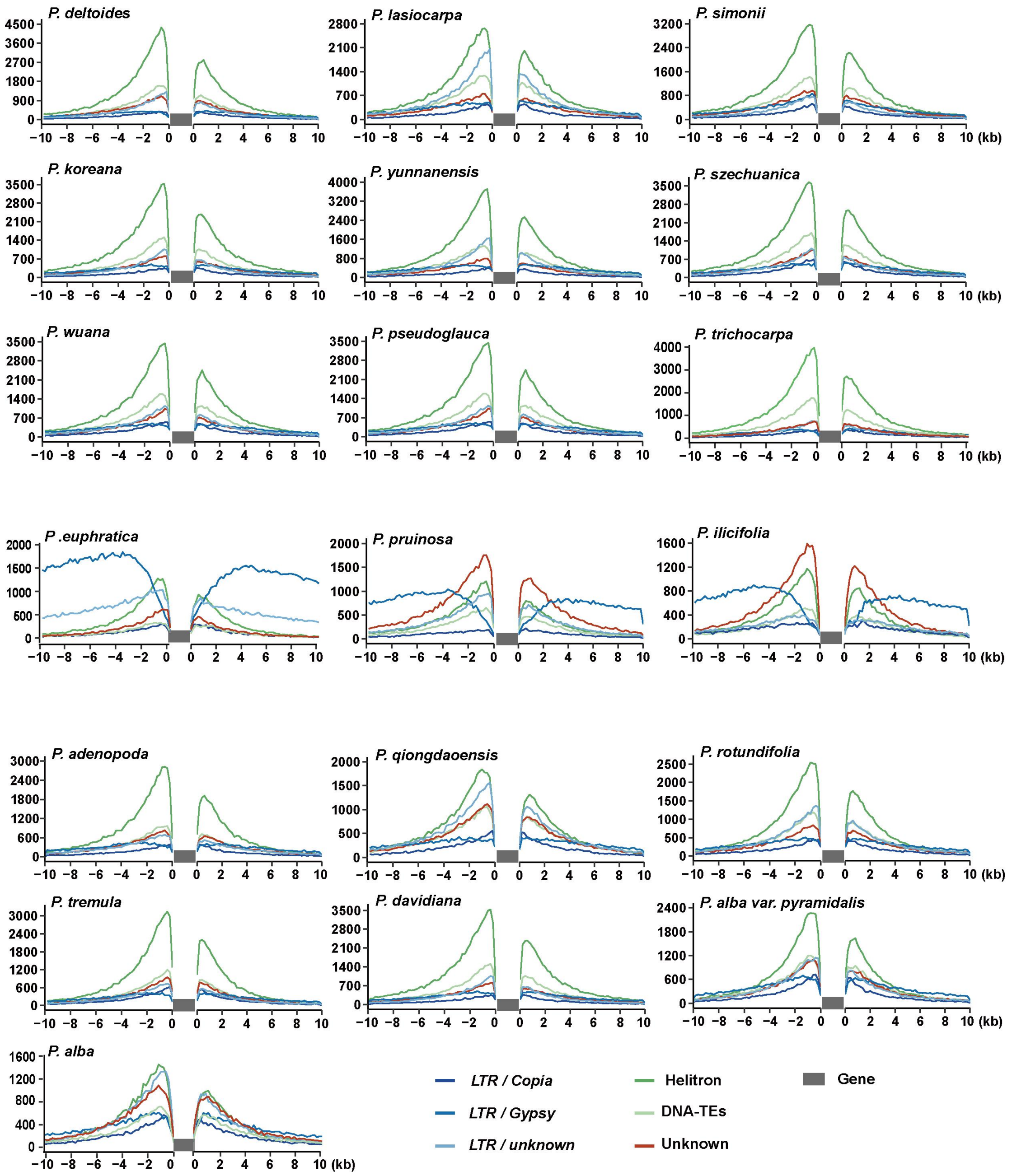


**Supplementary Fig. 5. TE landscape surrounding genes in poplars.** For all genes, the 10 kb upstream of the transcription start site and 10 kb downstream of the transcription end site were analyzed.

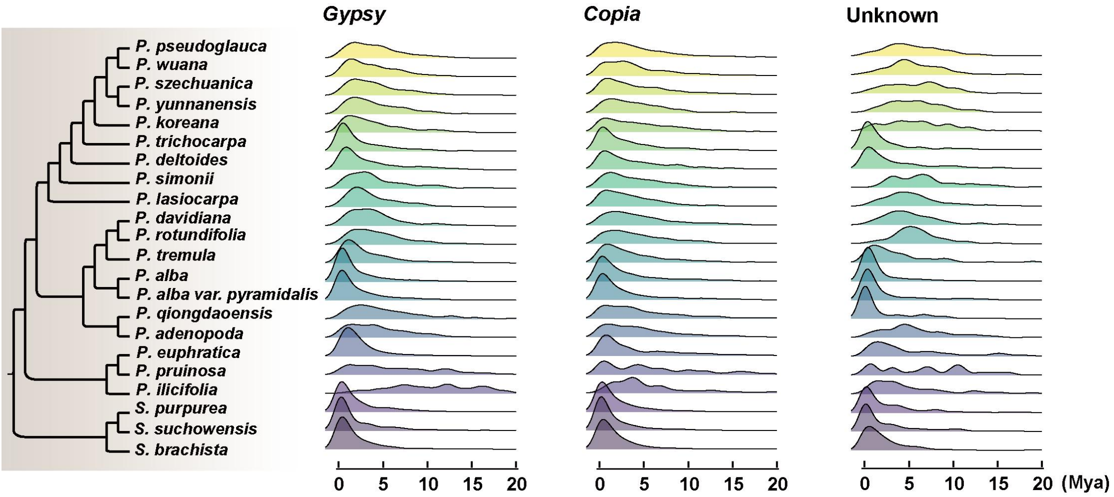


**Supplementary Fig. 6. Proliferation history of different classes of LTR retrotransposons (LTR-RTs).**

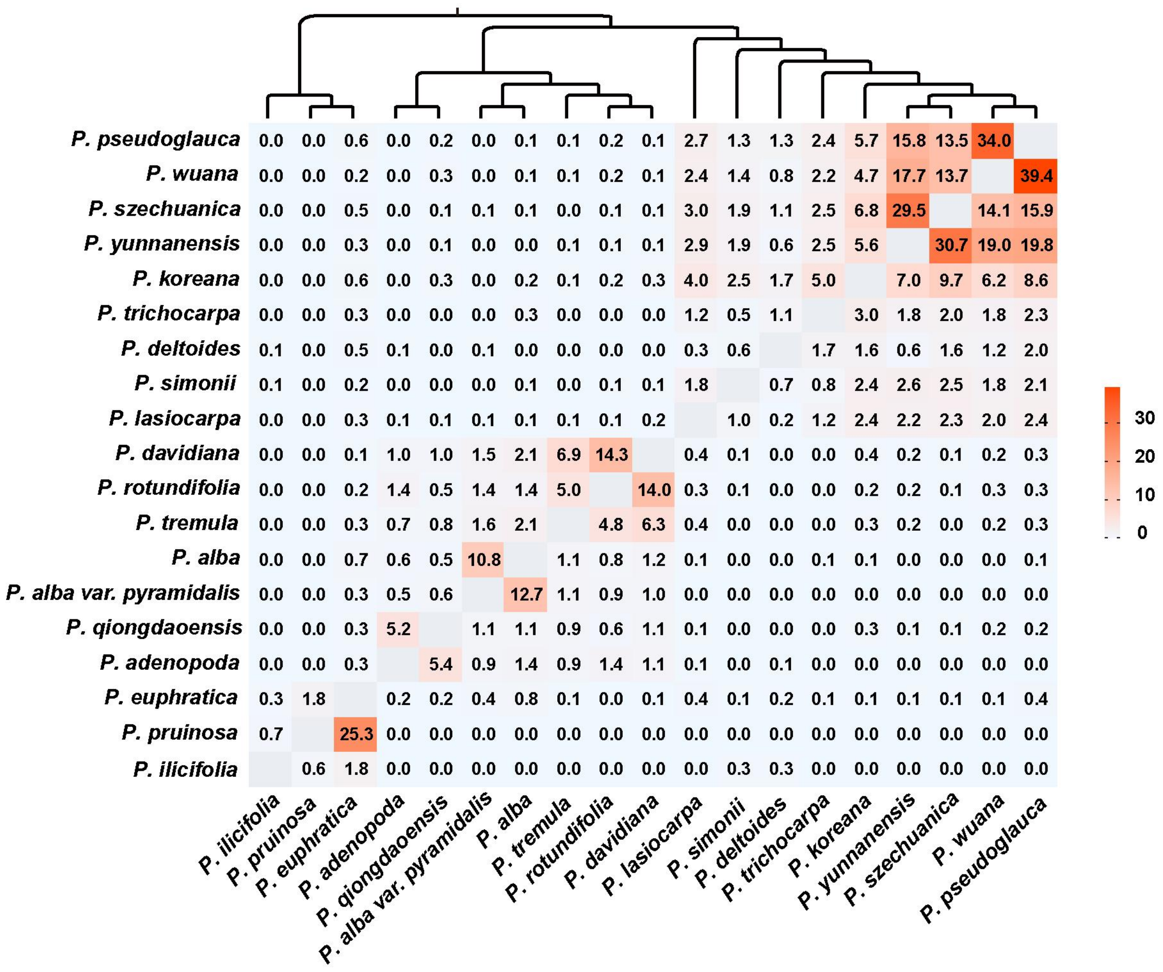


**Supplementary Fig. 7. Percentage of pairwise shared and still-intact fl-LTRs at syntenic positions for 19 *Populus* species.** Reading direction is column to row; for example, *P. davidiana* shares 14.0% of its fl-LTRs with *P. rotundifolia* and *P. rotundifolia* shares 14.3% with *P. davidiana*.

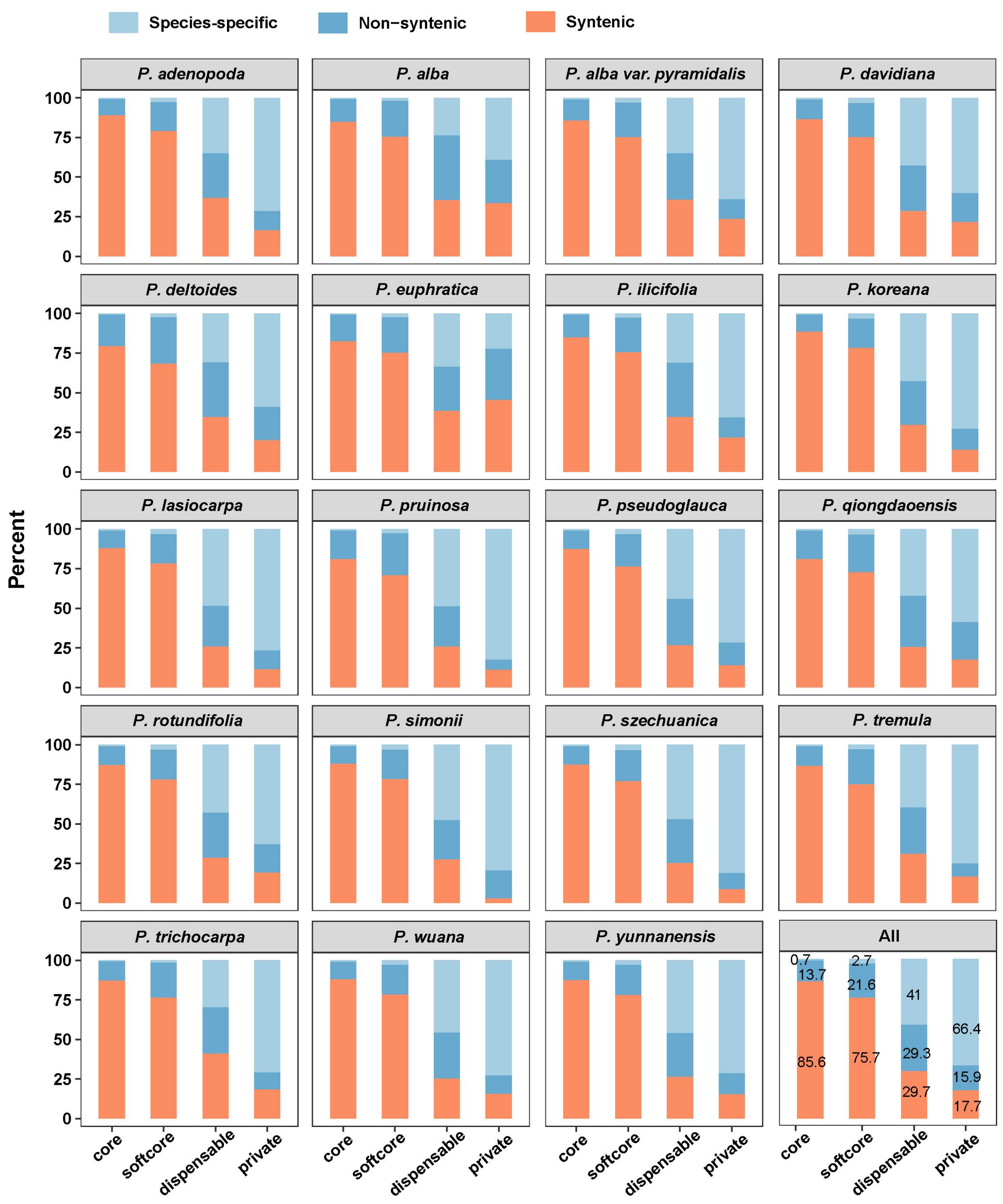


**Supplementary Fig. 8. Overall proportions of syntenic, non-syntenic, and species-specific loci in each genome based on their pan-genome classification.** Syntenic genes in each genome were calculated using *S. suchowensis* as query genome with MCScanX.

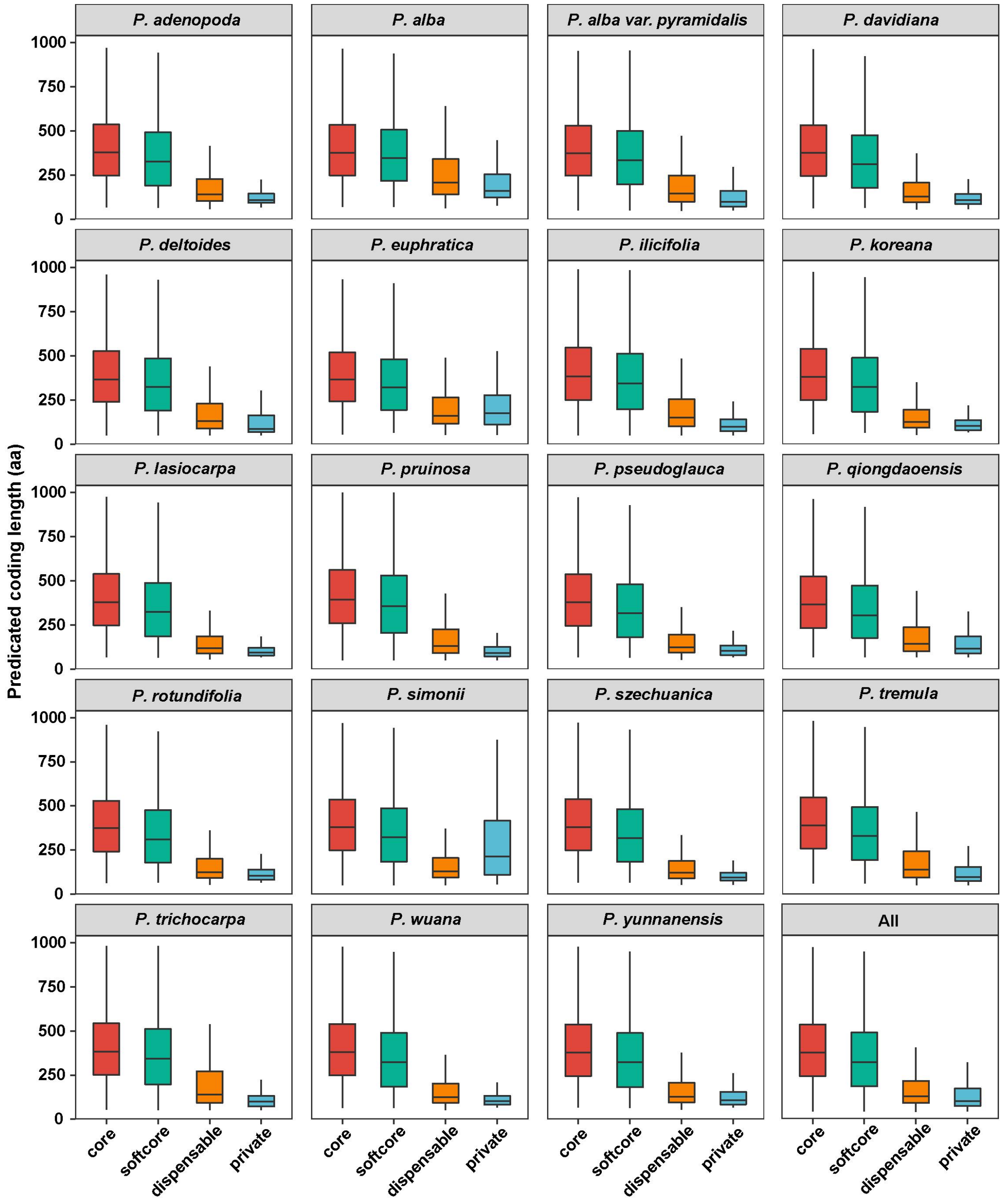


**Supplementary Fig. 9. CDS length of each gene in core, softcore, dispensable and private genes.**

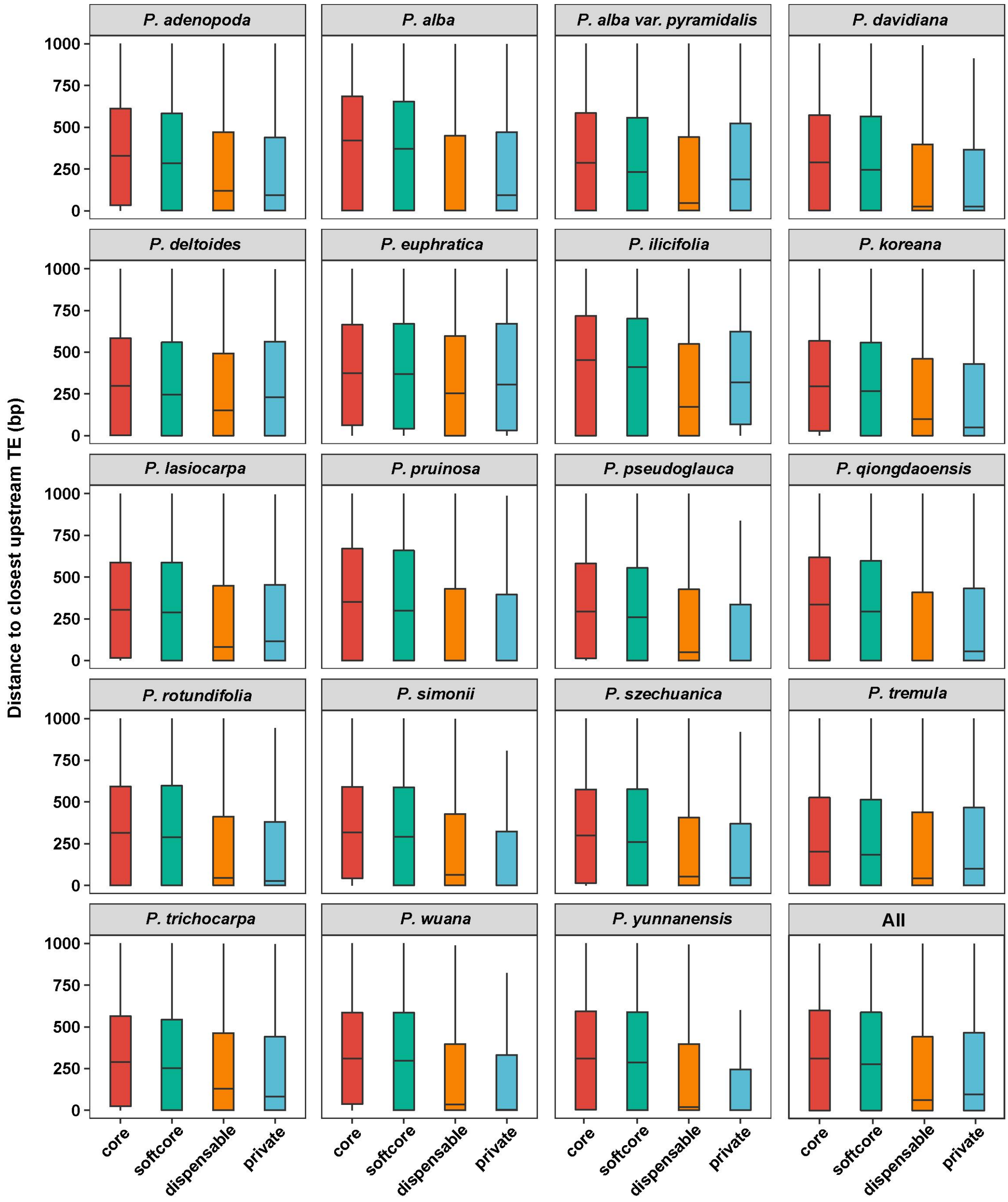


**Supplementary Fig. 10. Distance to closest upstream TE of each gene in core, softcore, dispensable and private genes.**

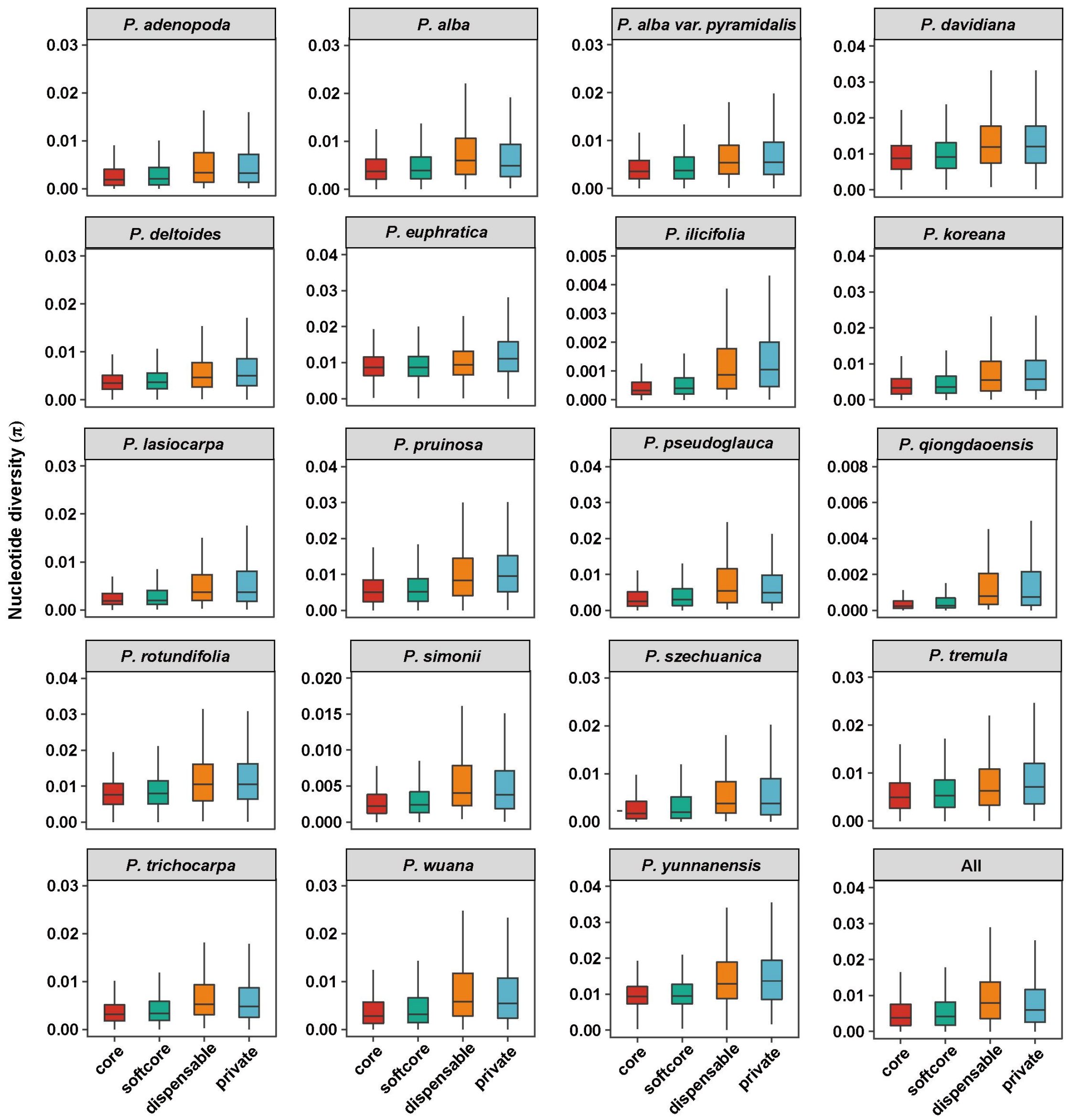


**Supplementary Fig. 11. Nucleotide diversity (π) of each gene in core, softcore, dispensable and private genes.**

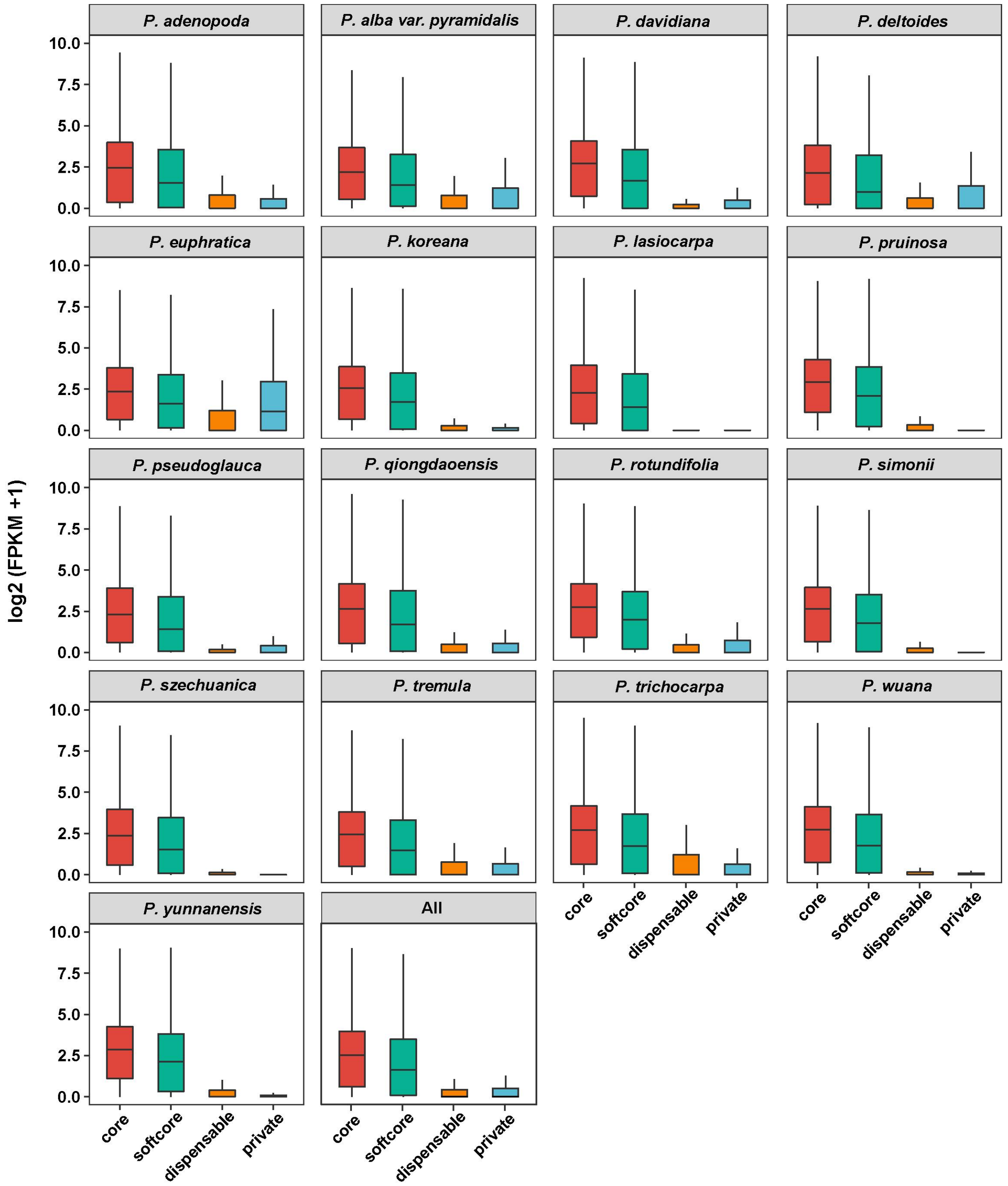


**Supplementary Fig. 12. The expression level (log2 FPKM in leaf tissue) of each gene in core, softcore, dispensable and private genes.**

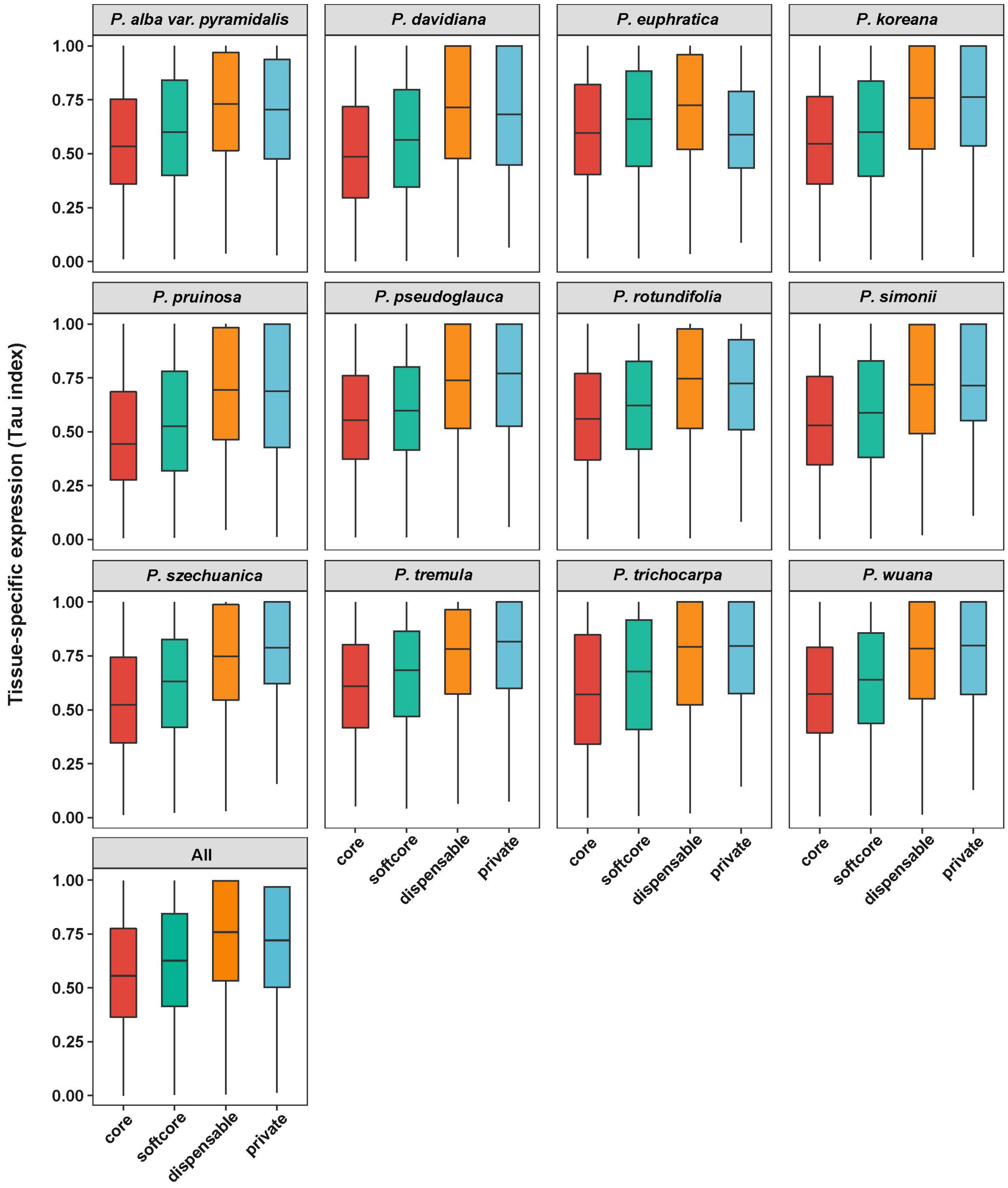


**Supplementary Fig. 13. Tissue-specific expression (Tau index) of each gene in core, softcore, dispensable and private genes.** Species with less than 3 tissues in the transcriptome dataset were not used for analysis.

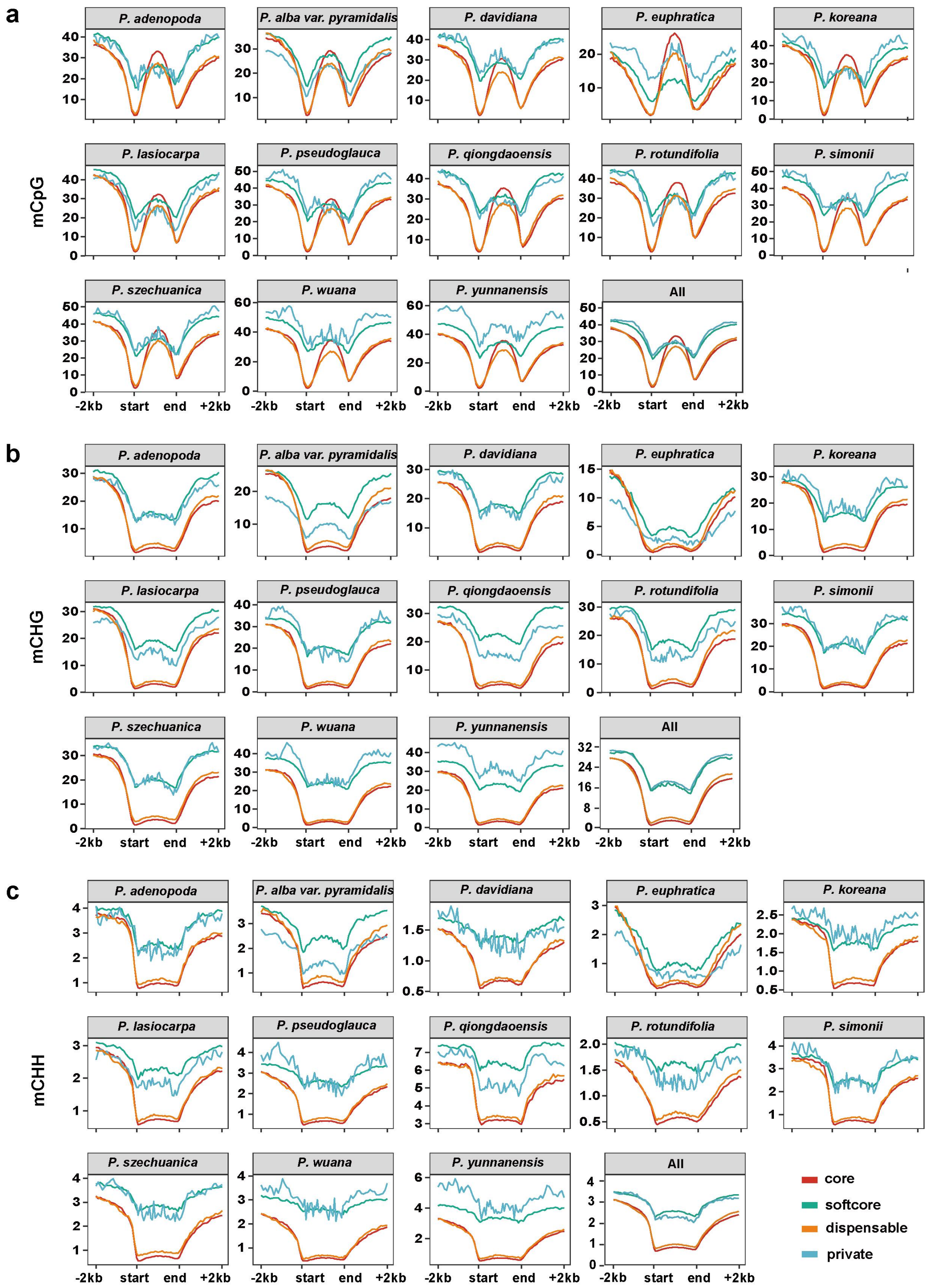


**Supplementary Fig. 14. Differences in average CG (a), CHG (b), and CHH (c) methylation level along the gene and flanking regions among core, softcore, dispensable and private genes.**

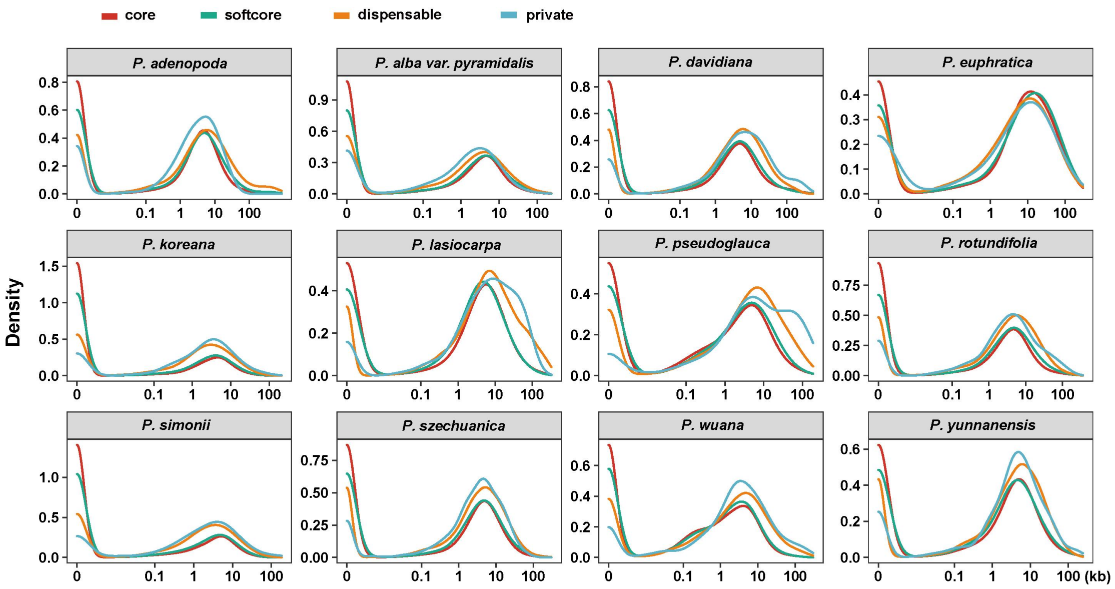


**Supplementary Fig. 15. A frequency distribution of accessible chromatin regions (ACRs) and their distance to the nearest genes based on different types of pan genes.**

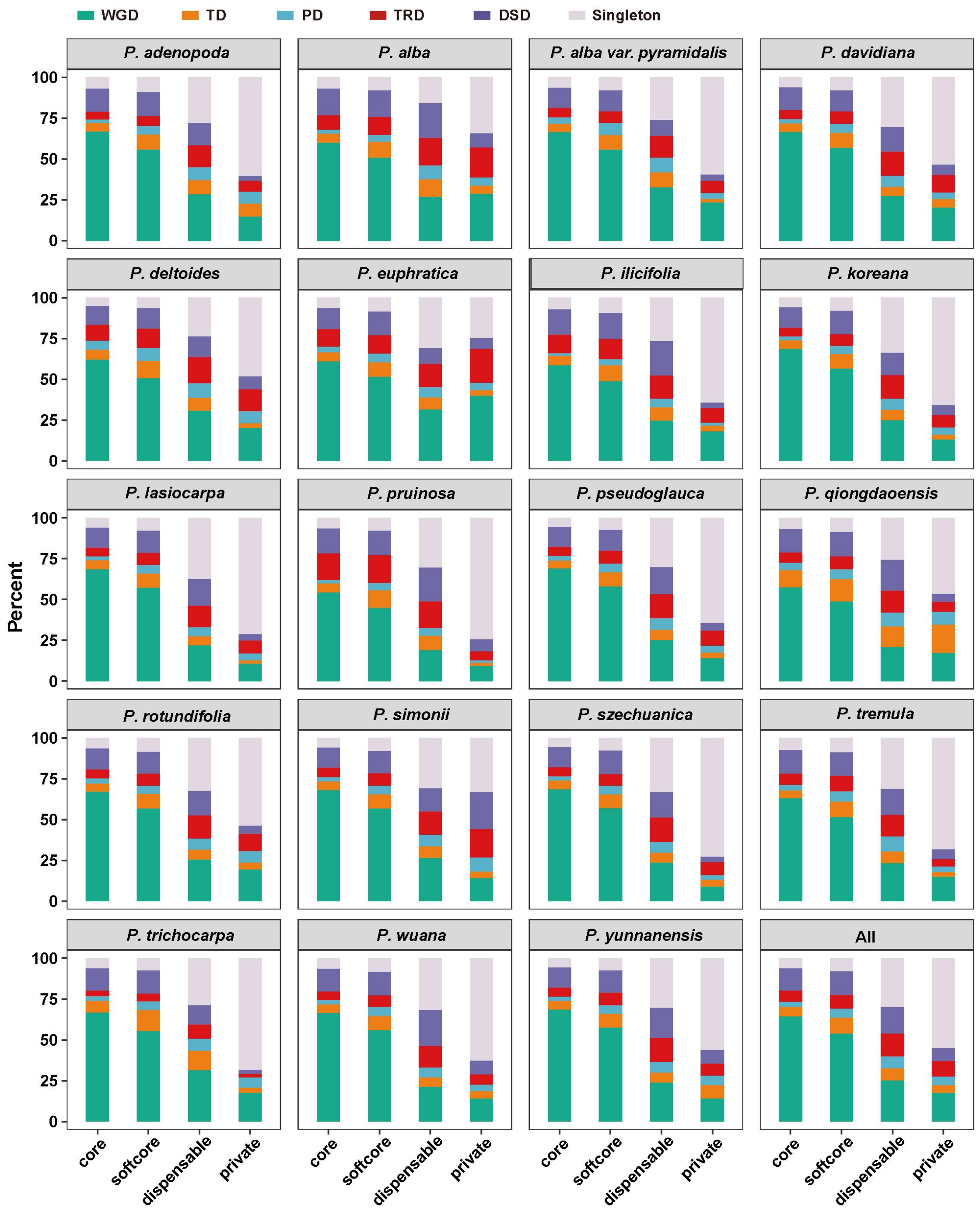


**Supplementary Fig. 16. Overall proportions of WGD, TD, PD, TRD, DSD and singleton loci in each genome based on their pan-genome classification.** WGD whole-genome duplication, TD tandem duplication, PD proximal duplication, TRD transposed duplication, DSD dispersed duplication.

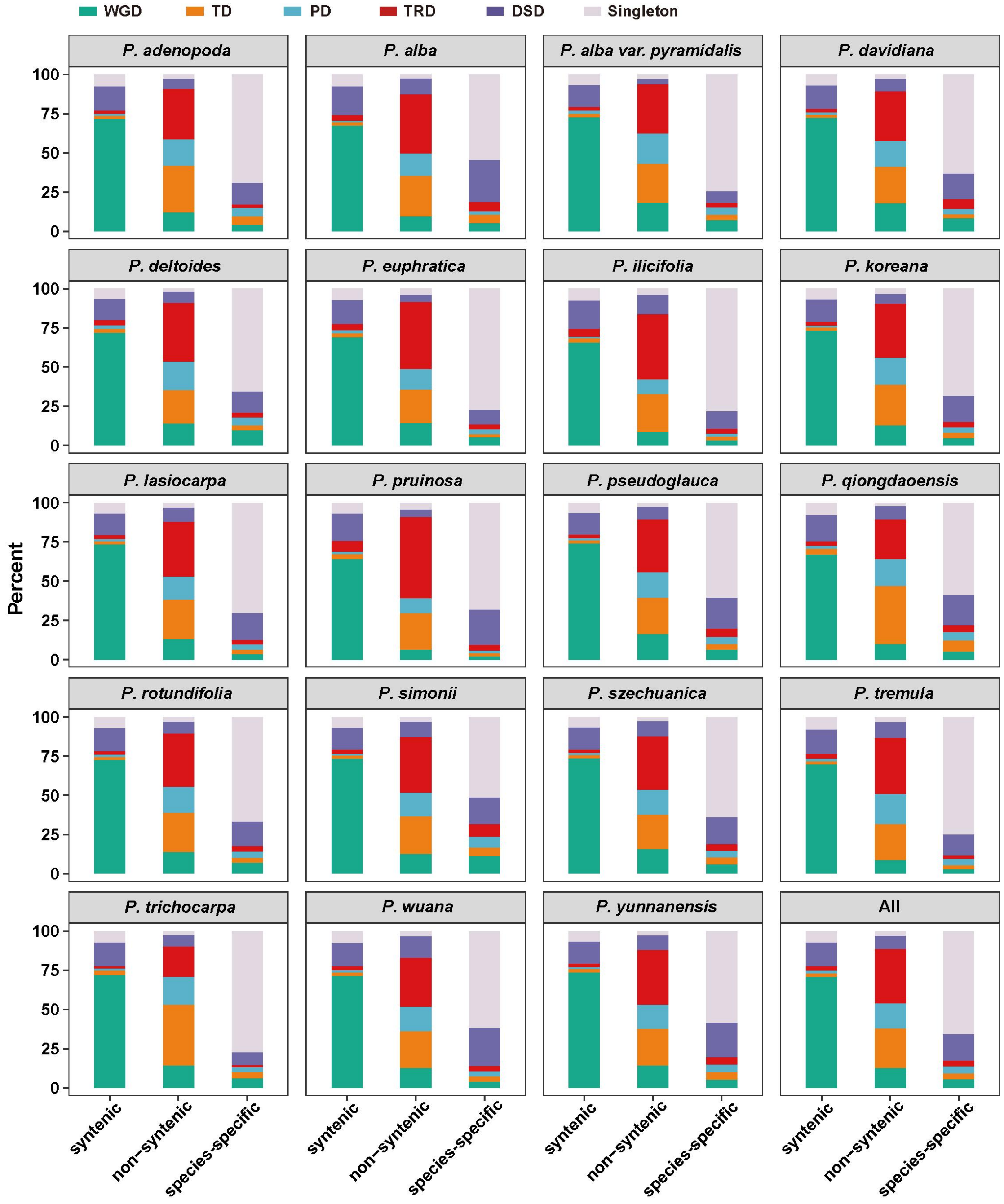


**Supplementary Fig. 17. Overall proportions of WGD, TD, PD, TRD, DSD and singleton loci in each genome based on their syntenic blocks with *S. suchowensis*.** Syntenic genes in each genome were calculated using MCScanX.

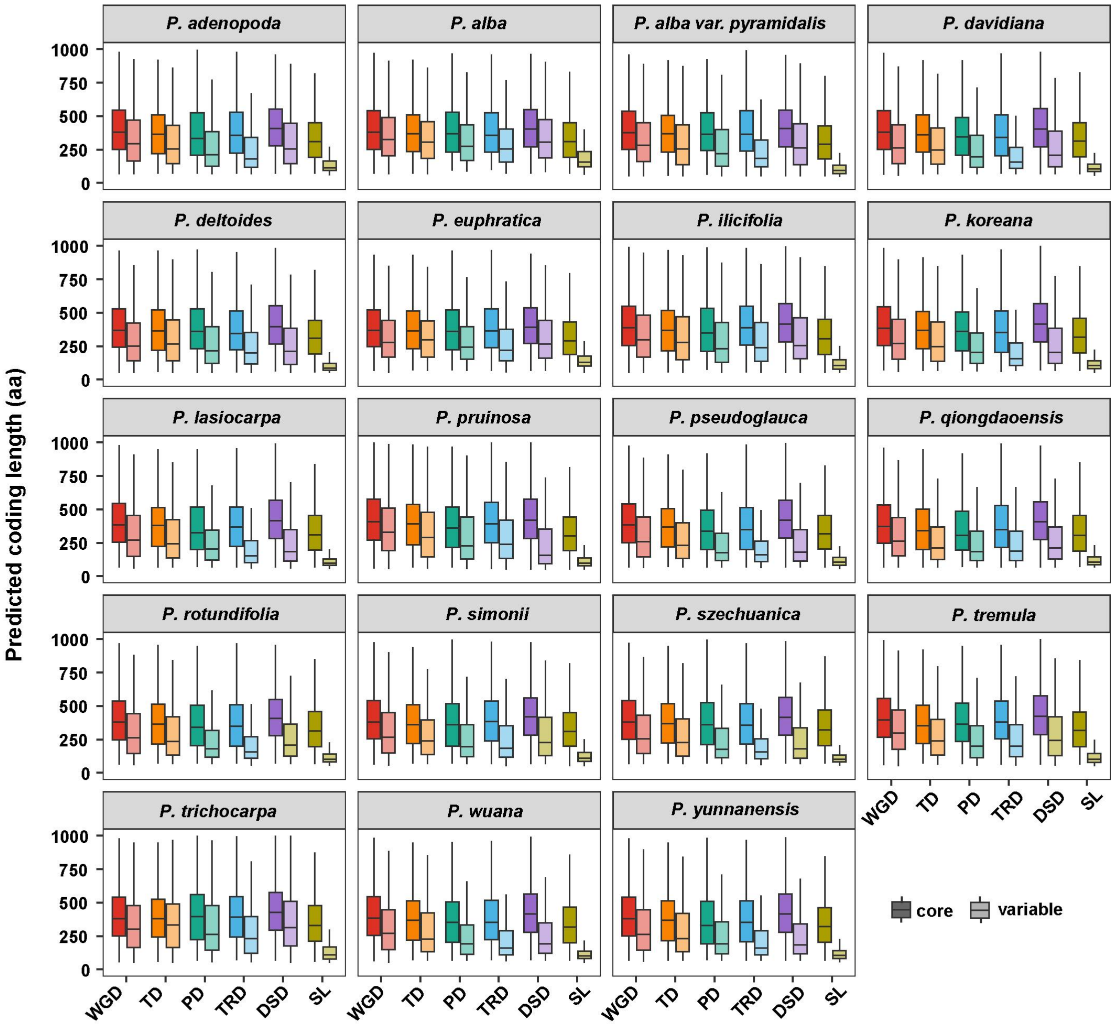


**Supplementary Fig. 18. CDS length of each gene that derived from different modes of duplication.**

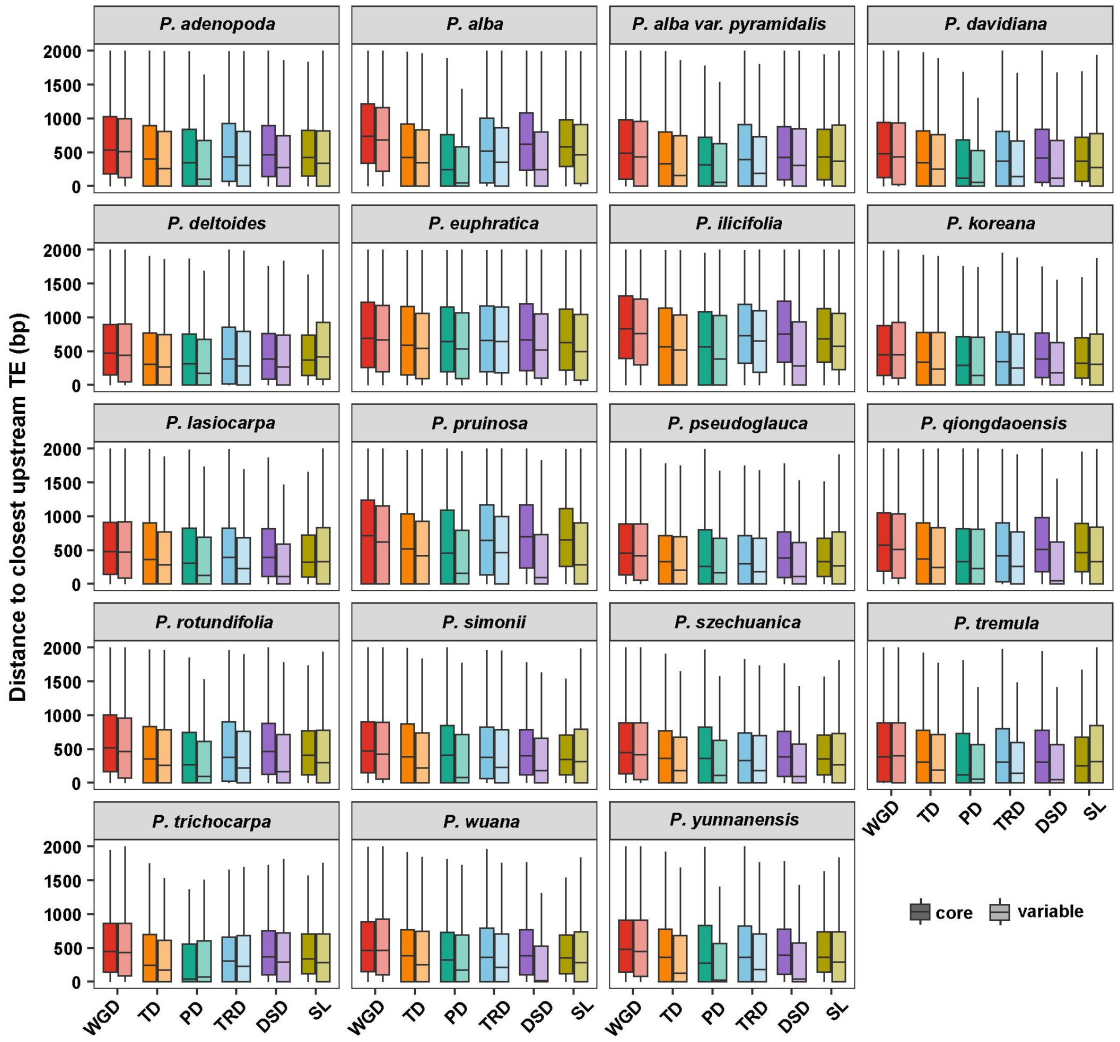


**Supplementary Fig. 19. Distance to closest upstream TE of each gene that derived from different modes of duplication.**

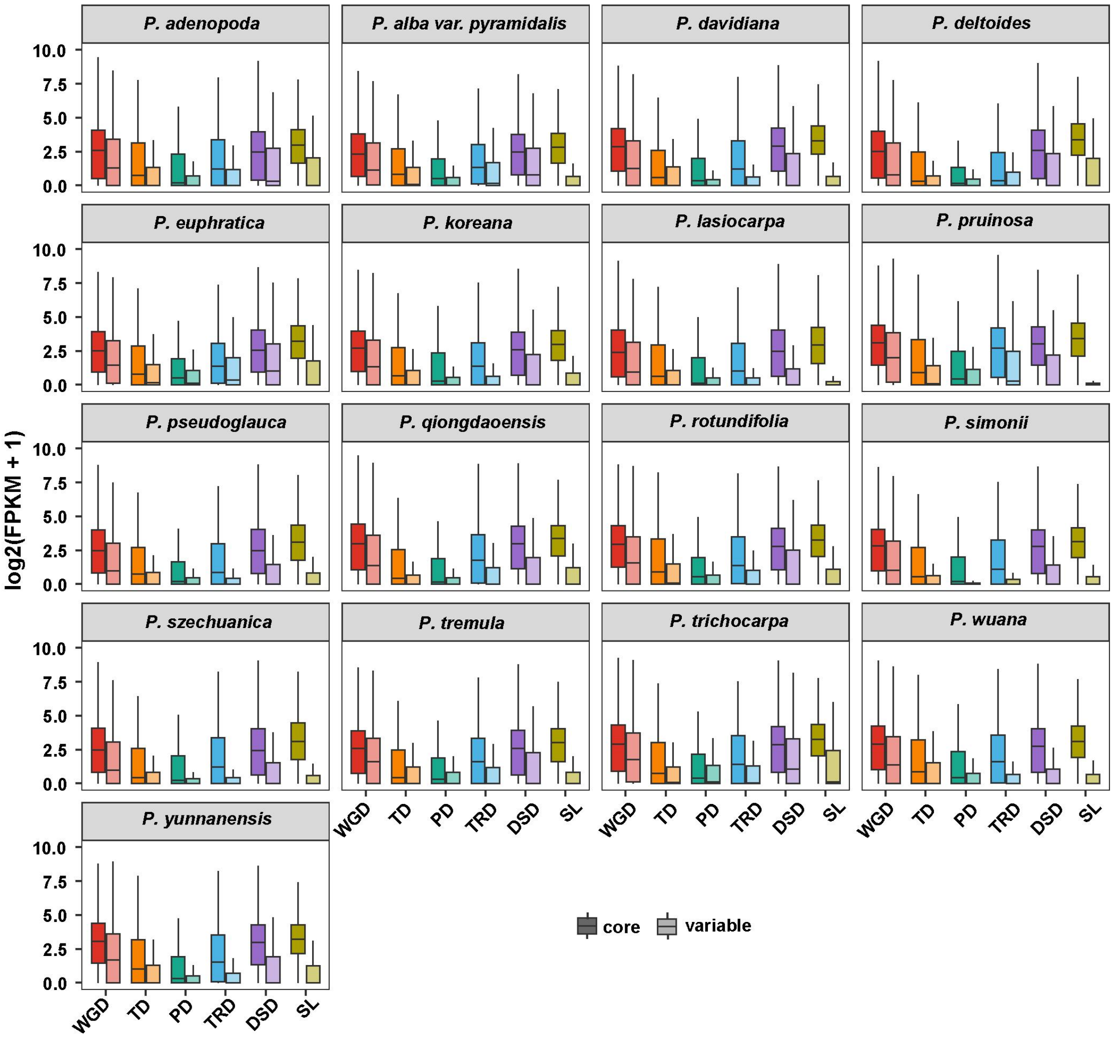


**Supplementary Fig. 20. The expression level (log2 FPKM in leaf tissue) of each gene that derived from different modes of duplication.**

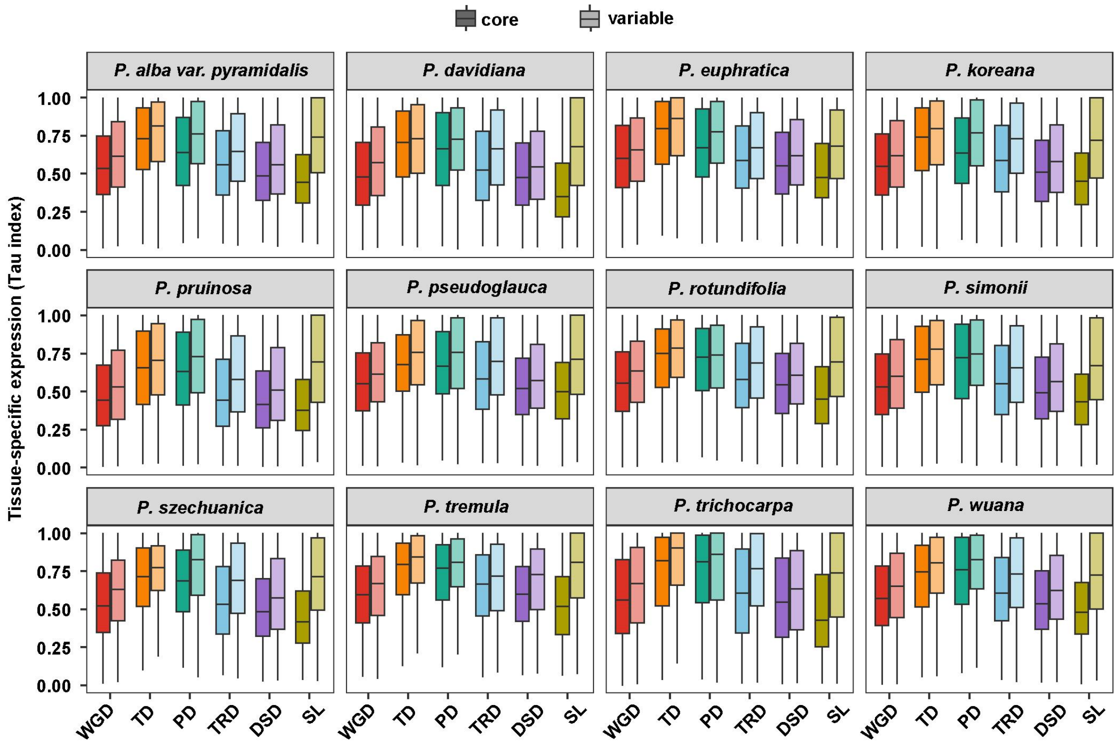


**Supplementary Fig. 21. Tissue-specific expression (Tau index) of each gene that derived from different modes of duplication.** Species with less than 3 tissues in the transcriptome dataset were not used for analysis.

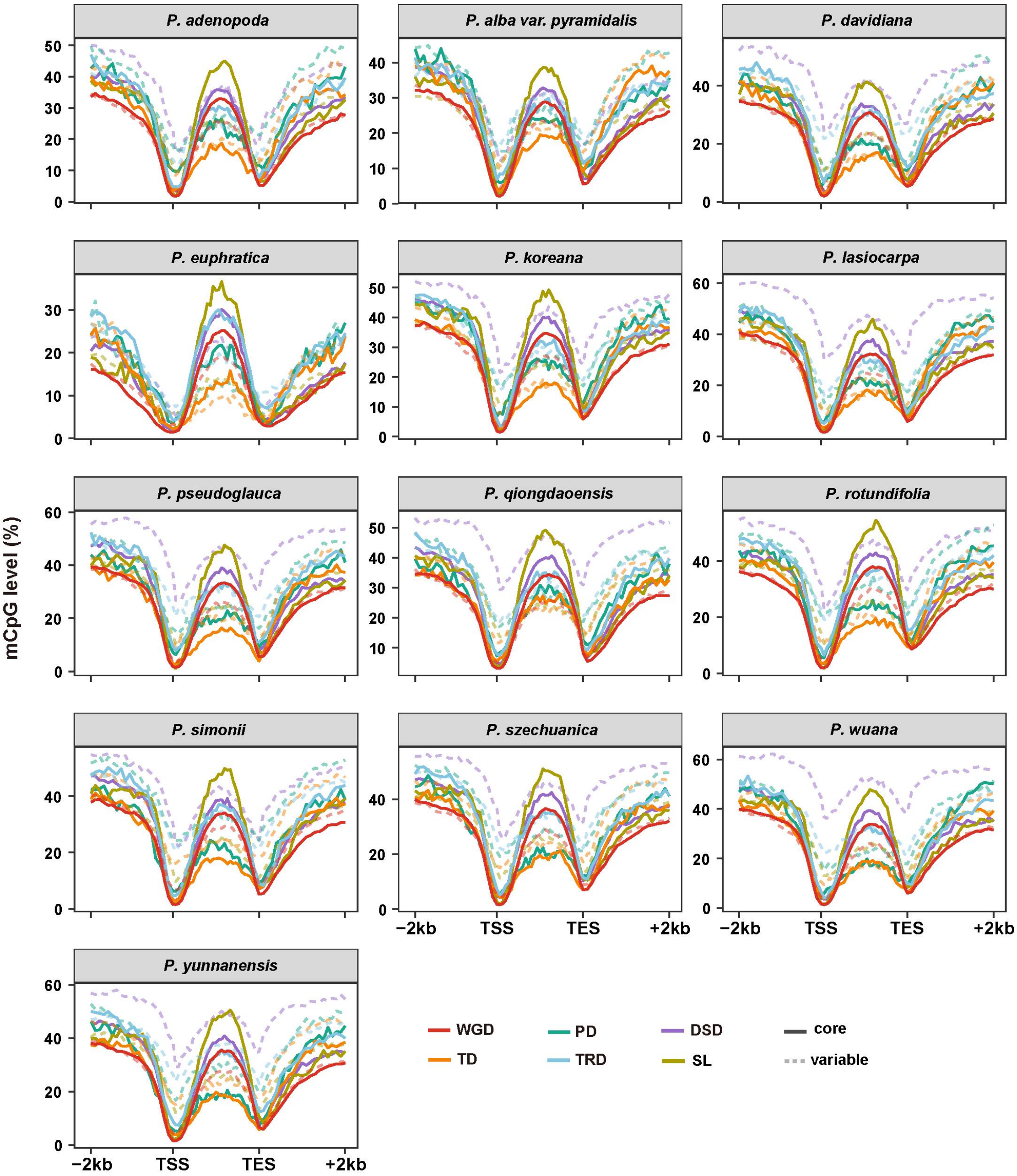
P. adenopoda
P. alba var. pyramidalis
P. davidiana
P. euphratica
P. koreana
P. lasiocarpa
P. pseudoglauca
P. qiongdaoensis
P. rotundifolia
P. simonii
P. szechuanica
P. wuana
P. yunnanensis
mCpG level (%)
−2kb
TSS
TES
+2kb
WGD
TD
PD
TRD
DSD
SL
core
variable

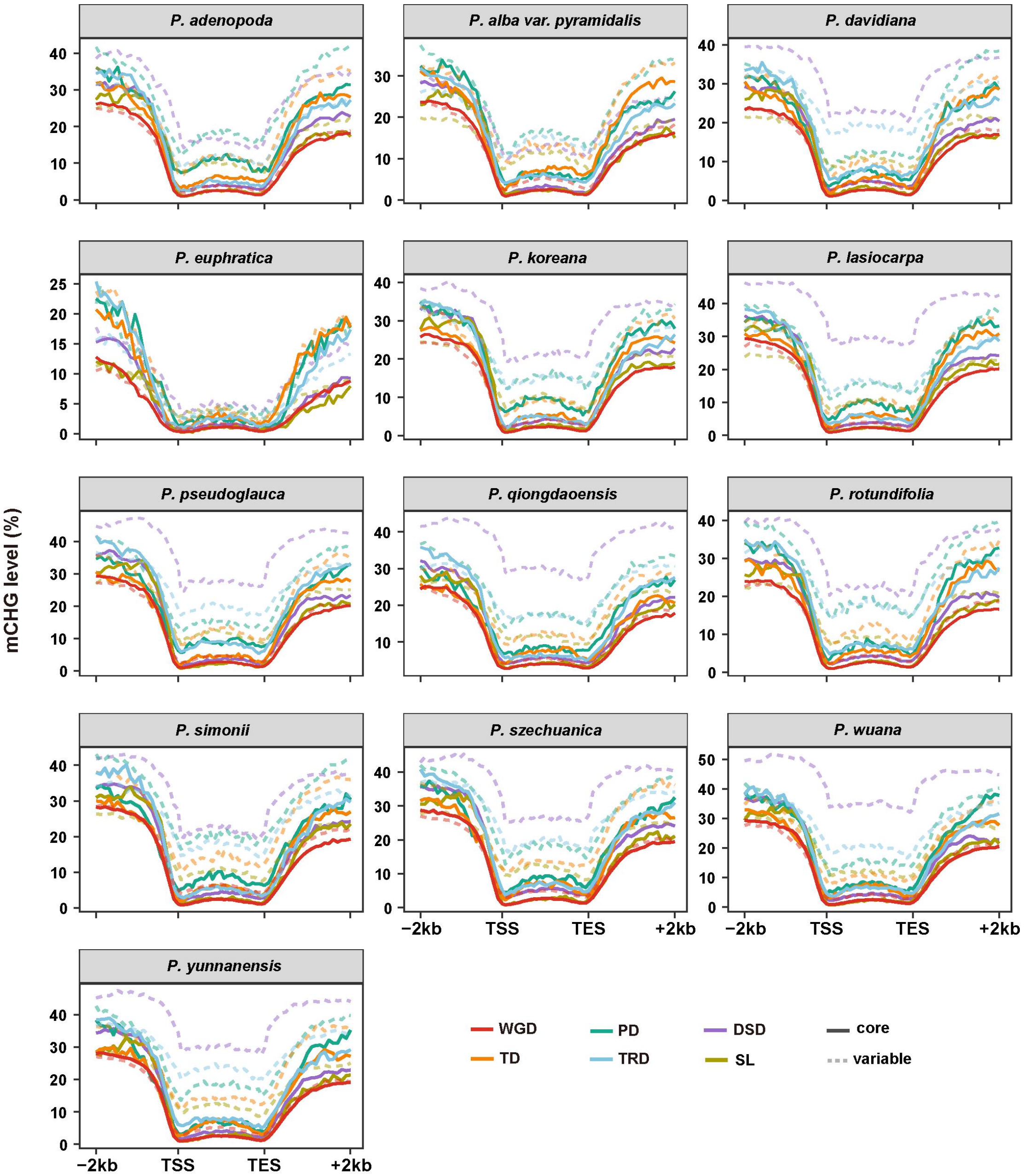

P. adenopoda
P. alba var. pyramidalis
P. davidiana
P. euphratica
P. koreana
P. lasiocarpa
P. pseudoglauca
P. qiongdaoensis
P. rotundifolia
P. simonii
P. szechuanica
P. wuana
P. yunnanensis
mCHG level (%)
−2kb
TSS
TES
+2kb
WGD
TD
PD
TRD
DSD
SL
core
variable

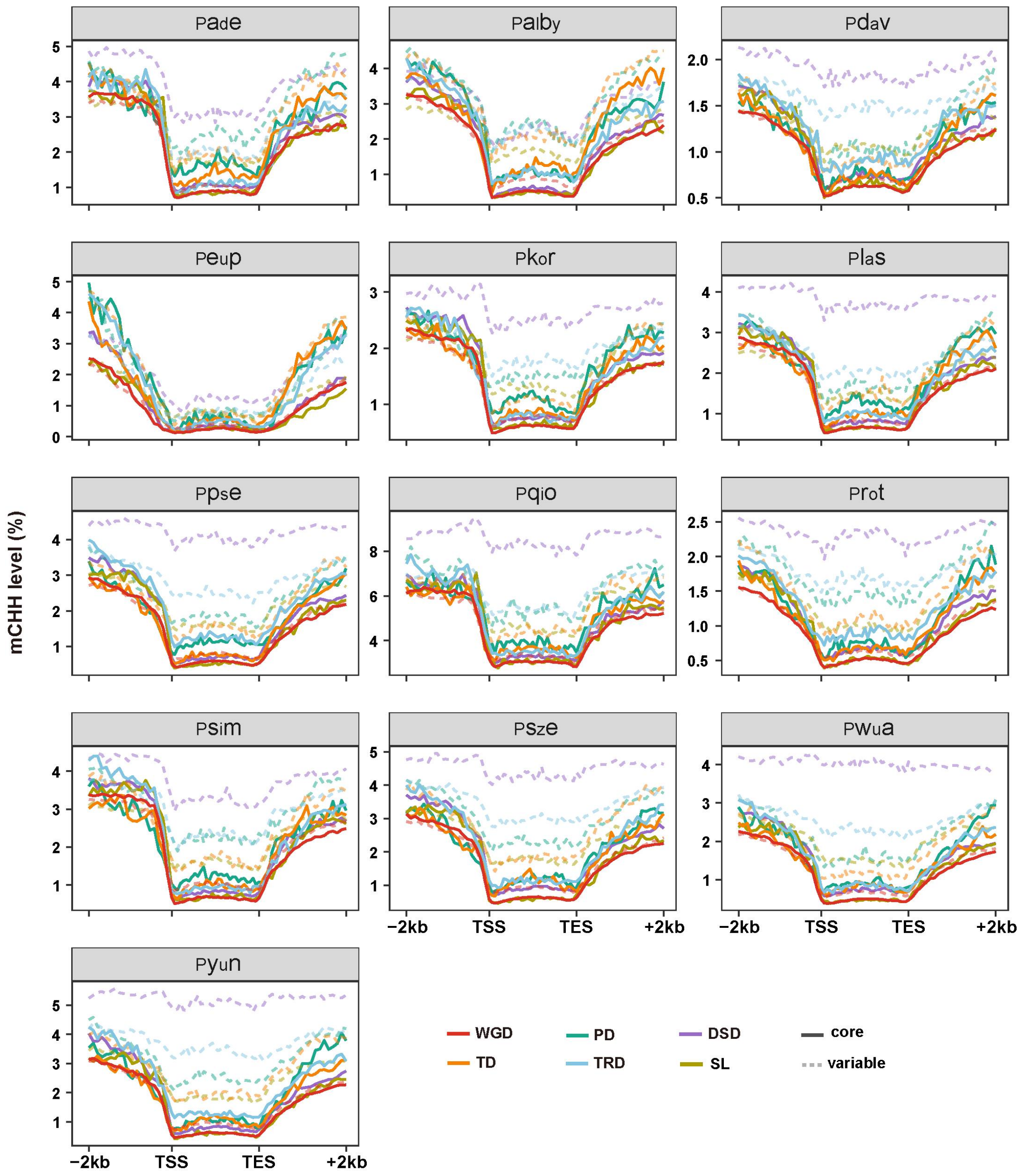


**Supplementary Fig. 22. Differences in average CG, CHG, and CHH methylation level along the gene and flanking regions among different gene groups based on the duplication status.**

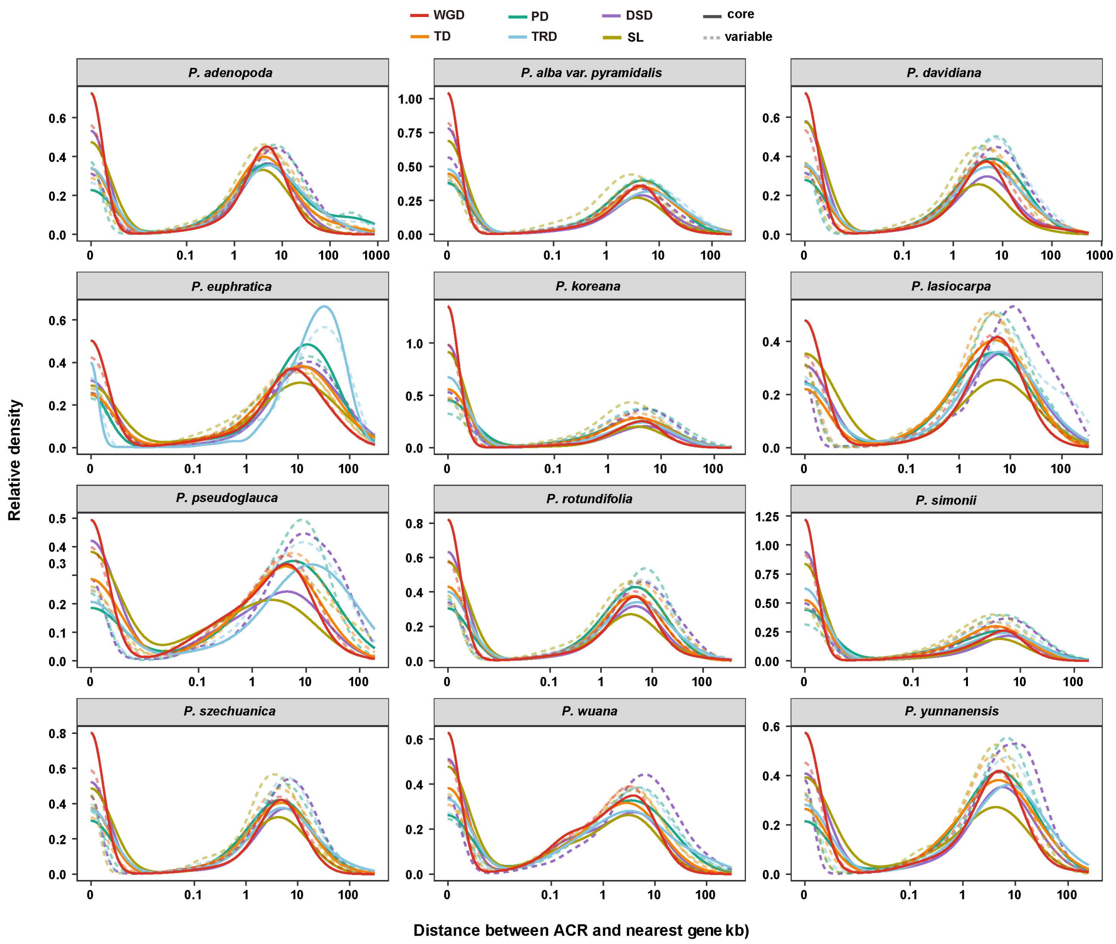


**Supplementary Fig. 23. A frequency distribution of ACRs and their distance to the nearest genes based on different modes of duplicated genes.**

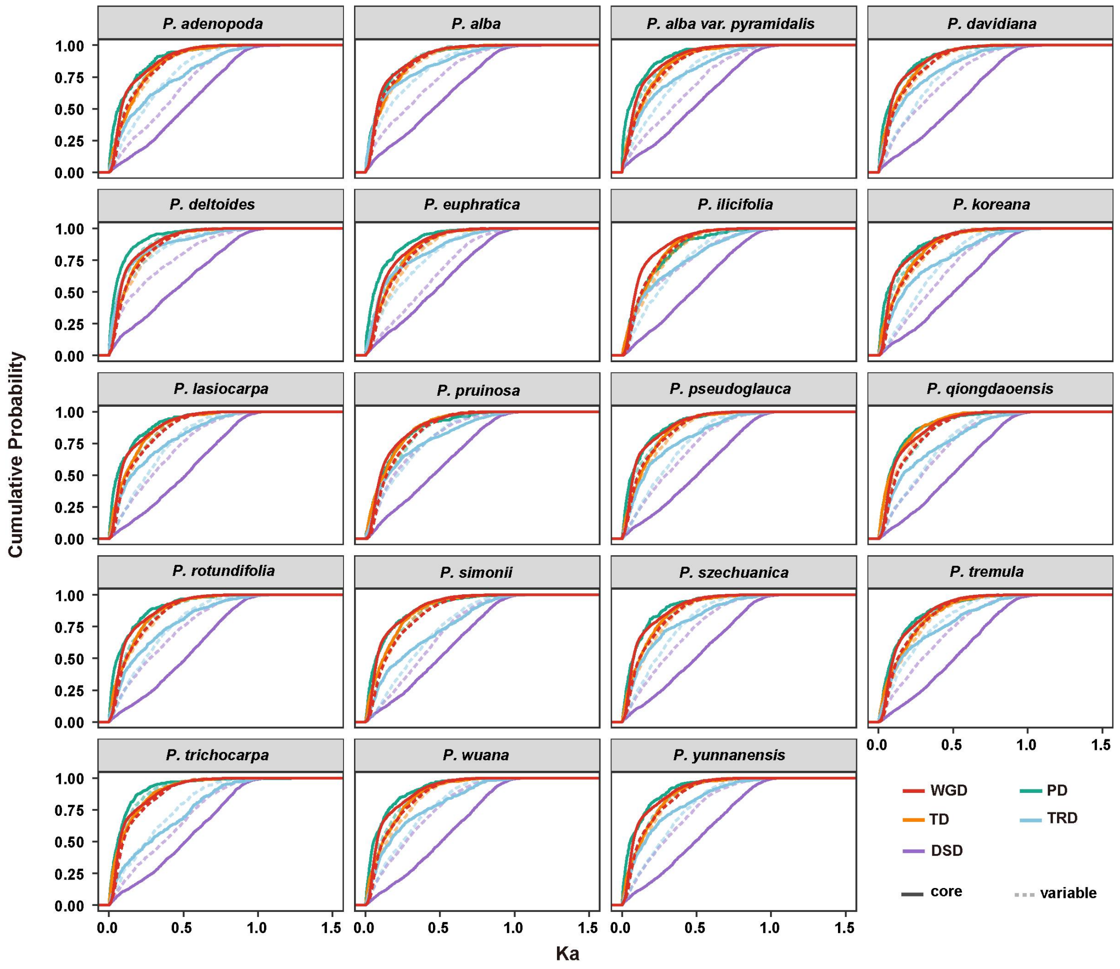


**Supplementary Fig. 24. The Ka distributions of gene pairs derived from different modes of duplication.**

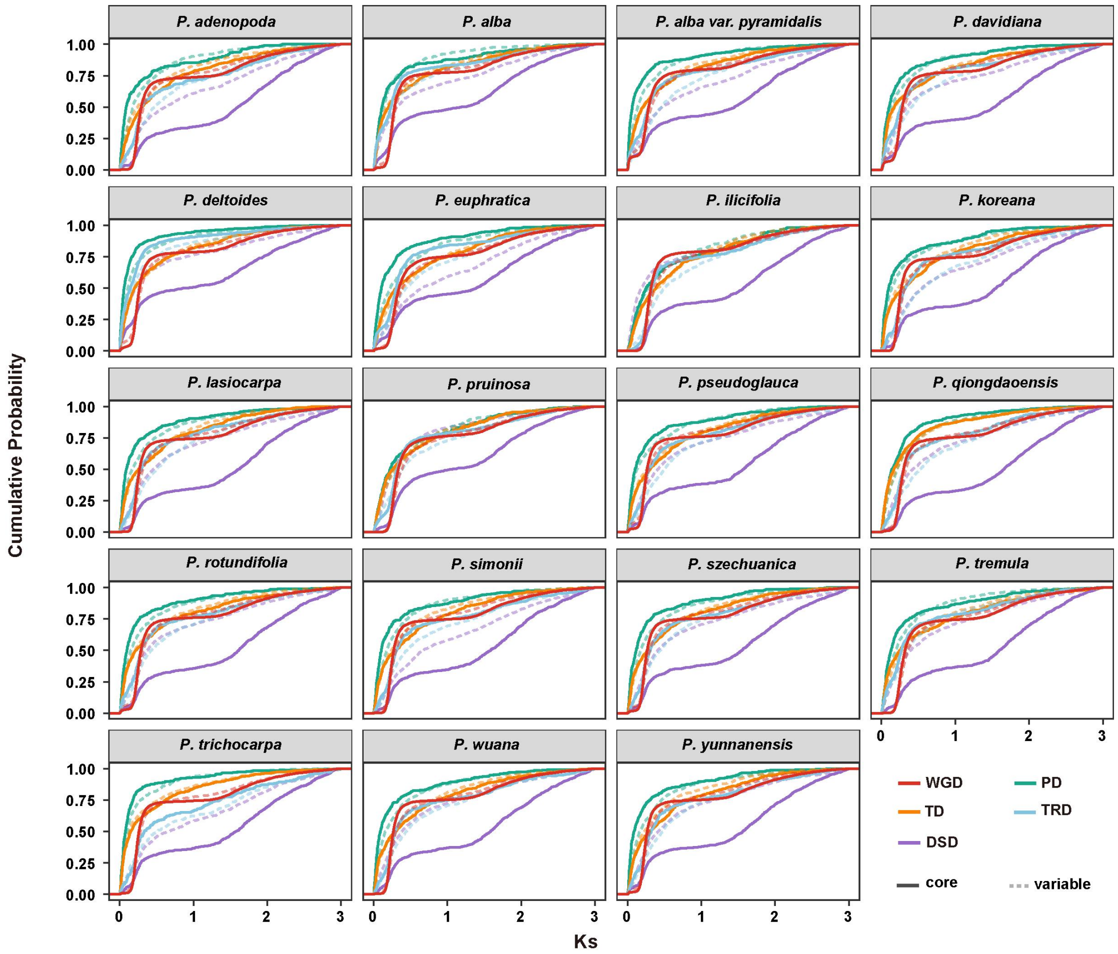


**Supplementary Fig. 25. The Ks distributions of gene pairs derived from different modes of duplication.**

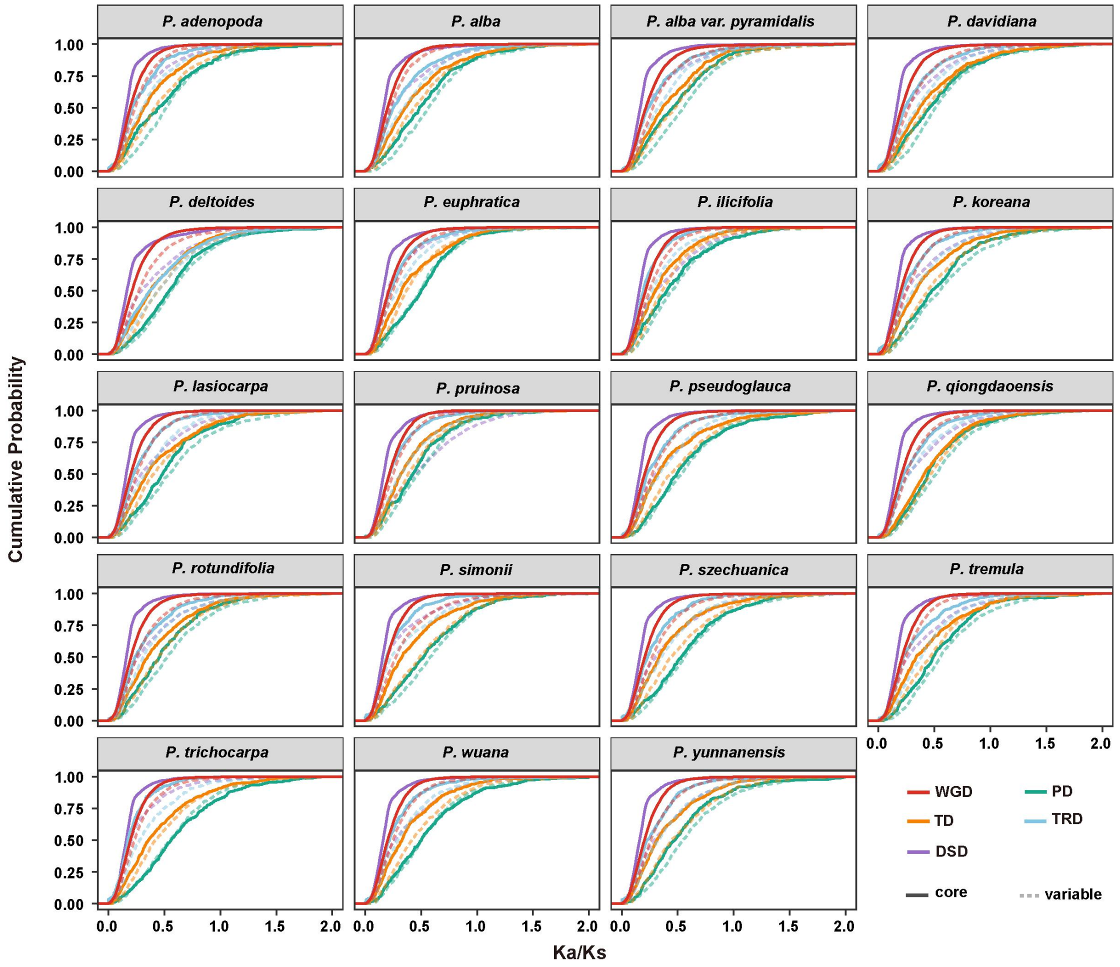


**Supplementary Fig. 26. The *Ka/Ks* ratio distributions of gene pairs derived from different modes of duplication.**

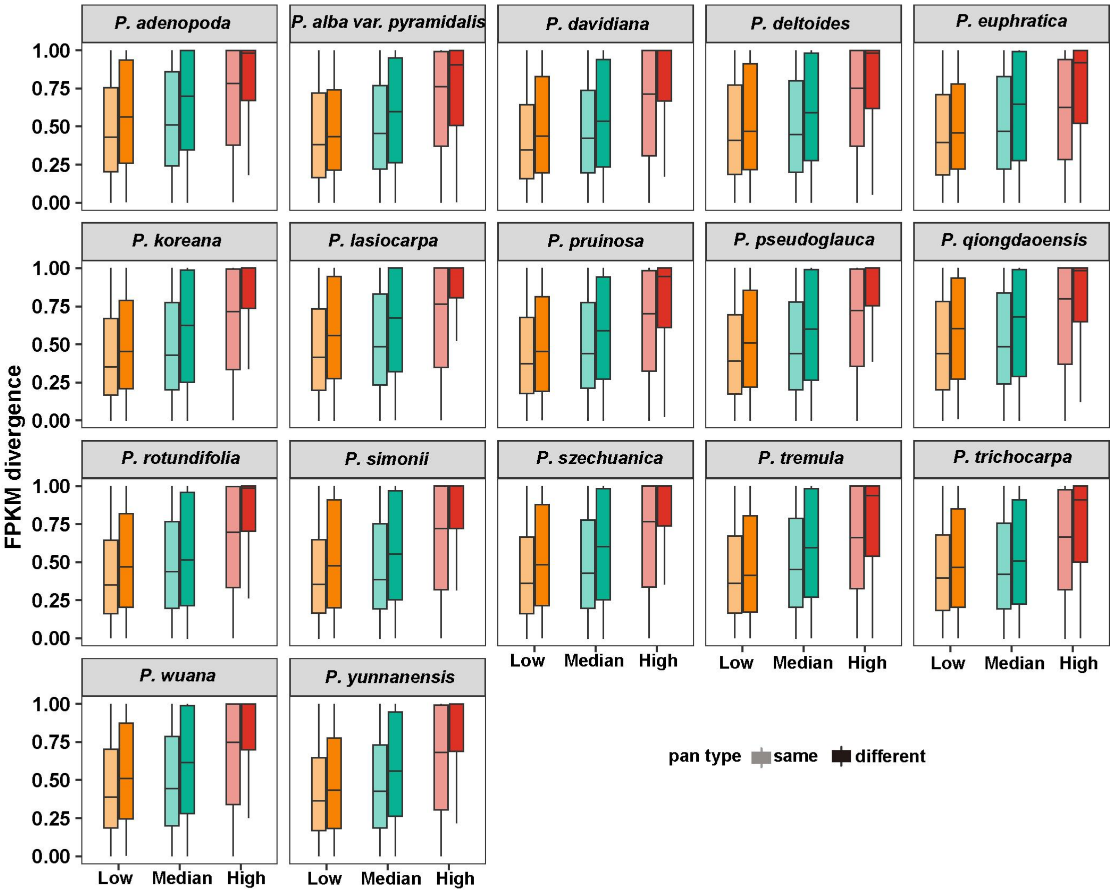


**Supplementary Fig. 27. Expression divergences of different WGD-derived duplicated gene groups divided by their *Ks* value and pan-gene type.**

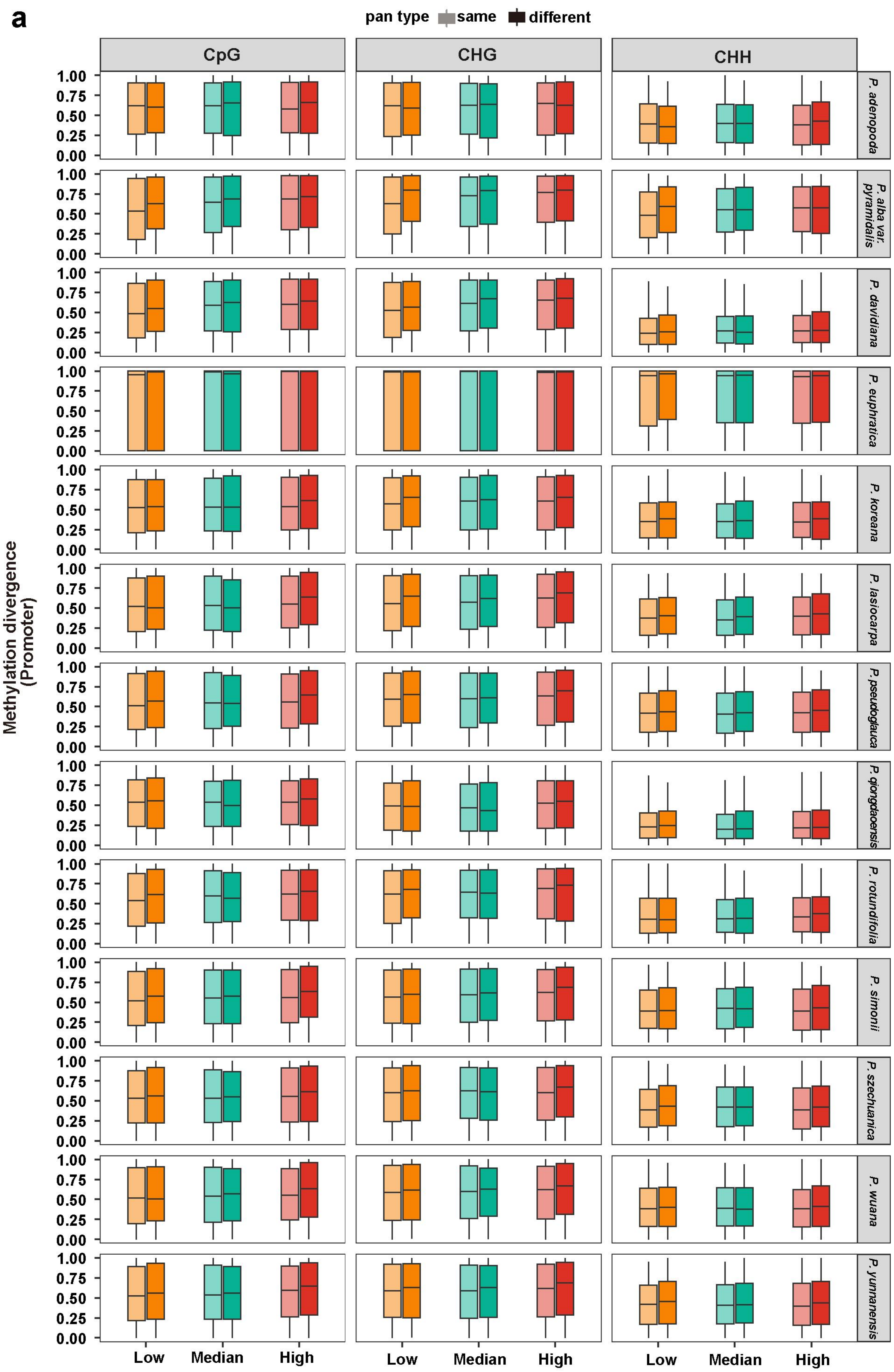


**Supplementary Fig. 28. Continued on next page.**

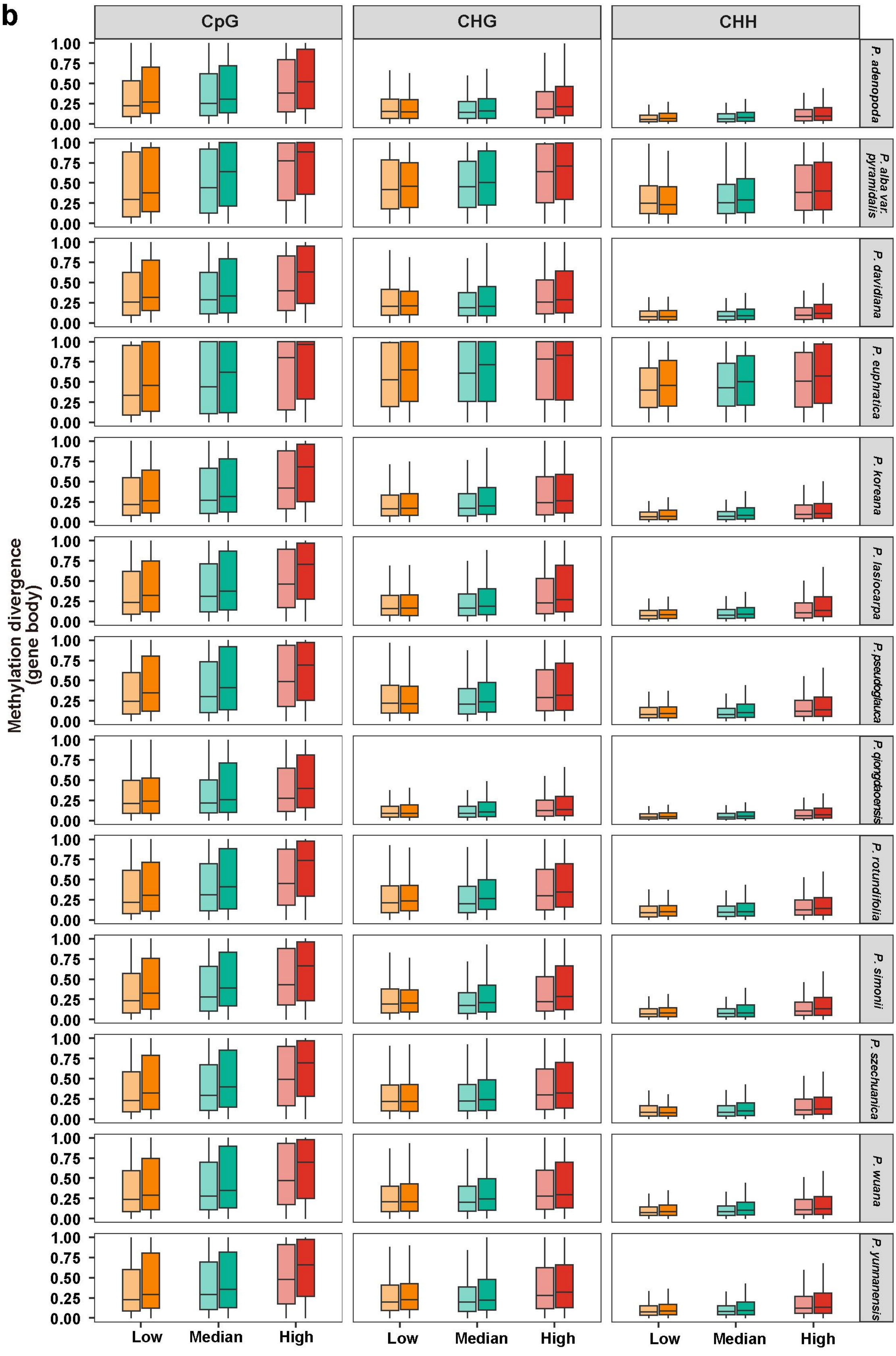


**Supplementary Fig. 28. Continued on next page.**

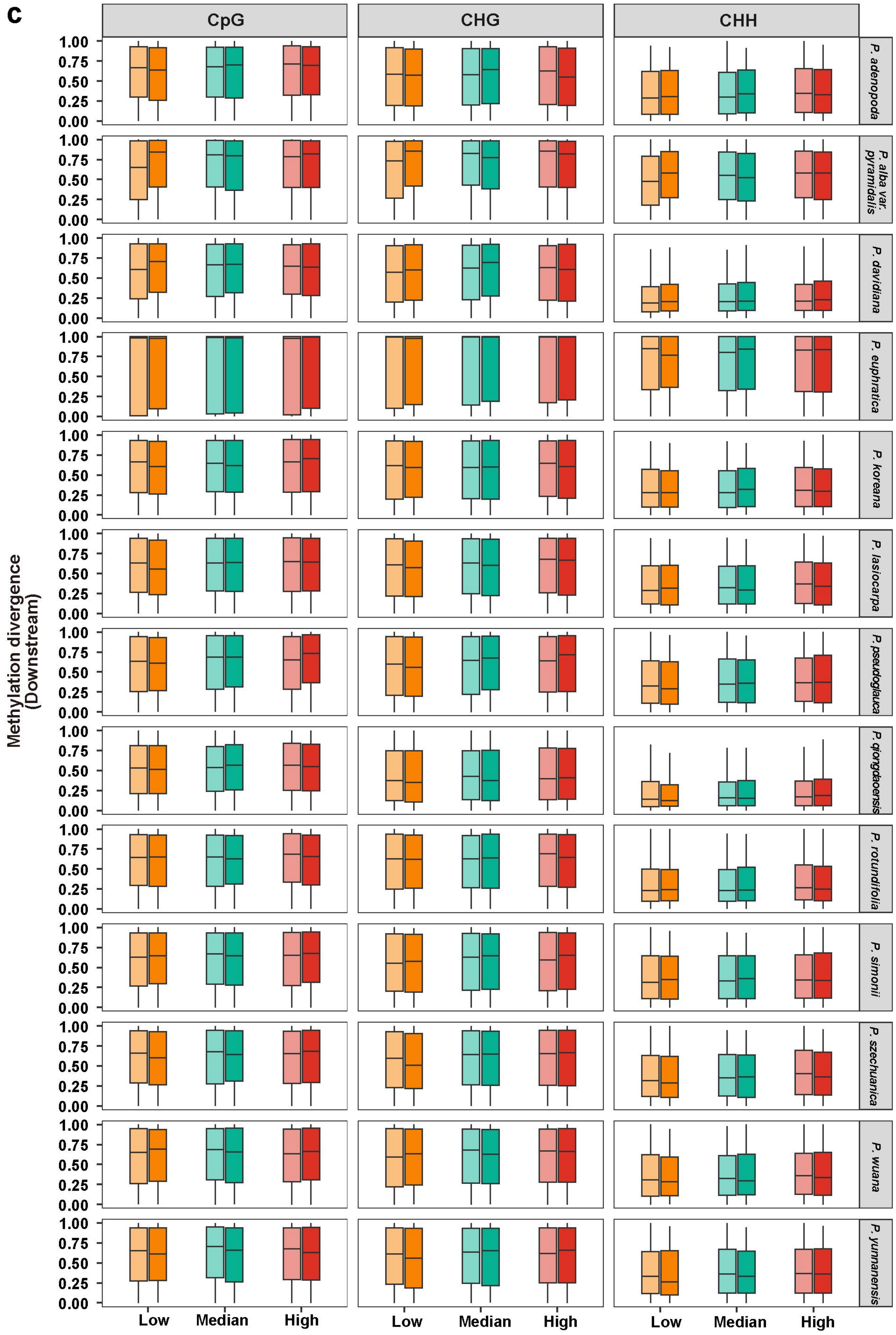


**Supplementary Fig. 28. Methylation divergences in promoter (a), gene-body (b) and downstream (c) regions of different WGD-derived duplicated gene groups divided by their *Ks* value and pan-gene type.**

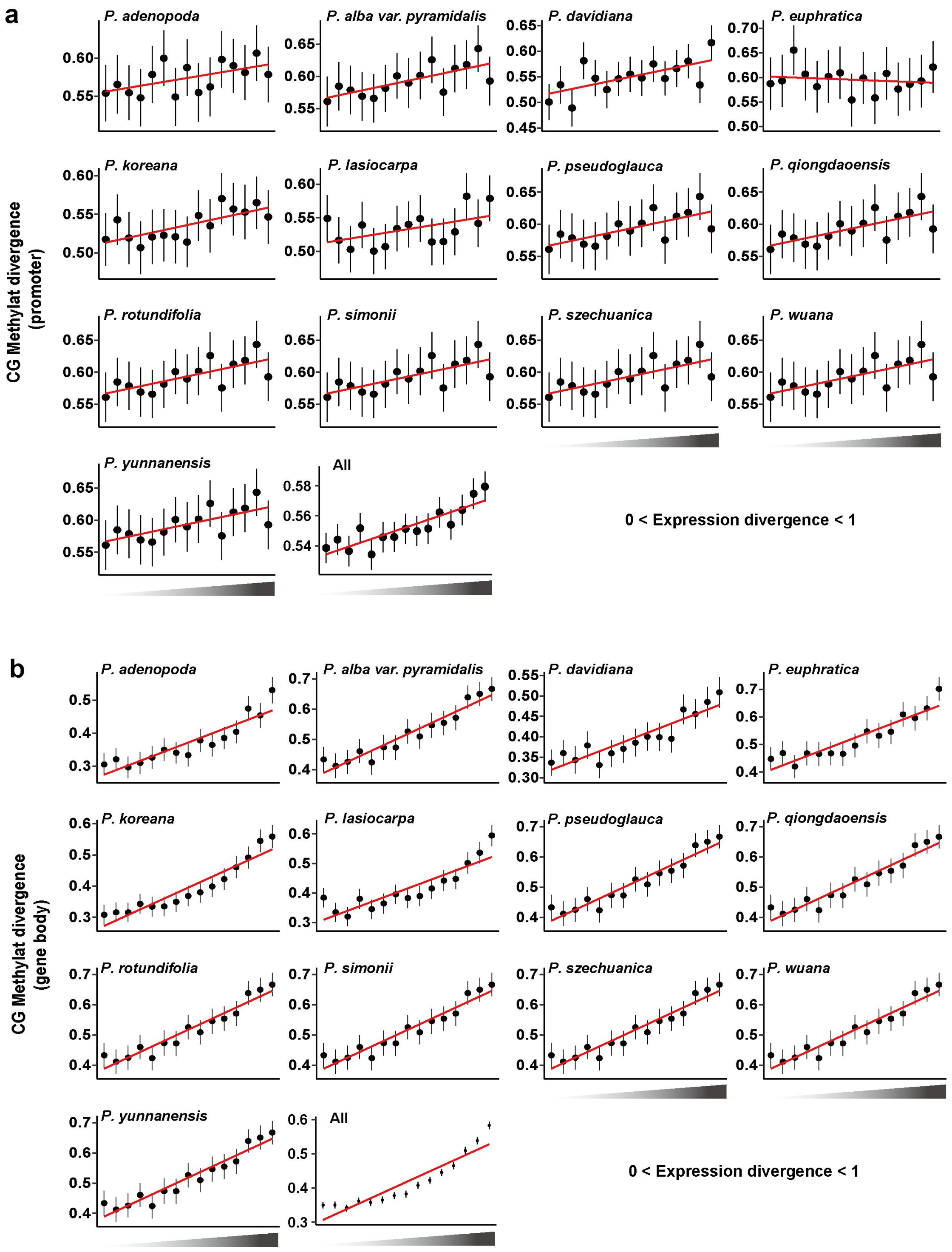


**Supplementary Fig. 29. Continued on next page.**

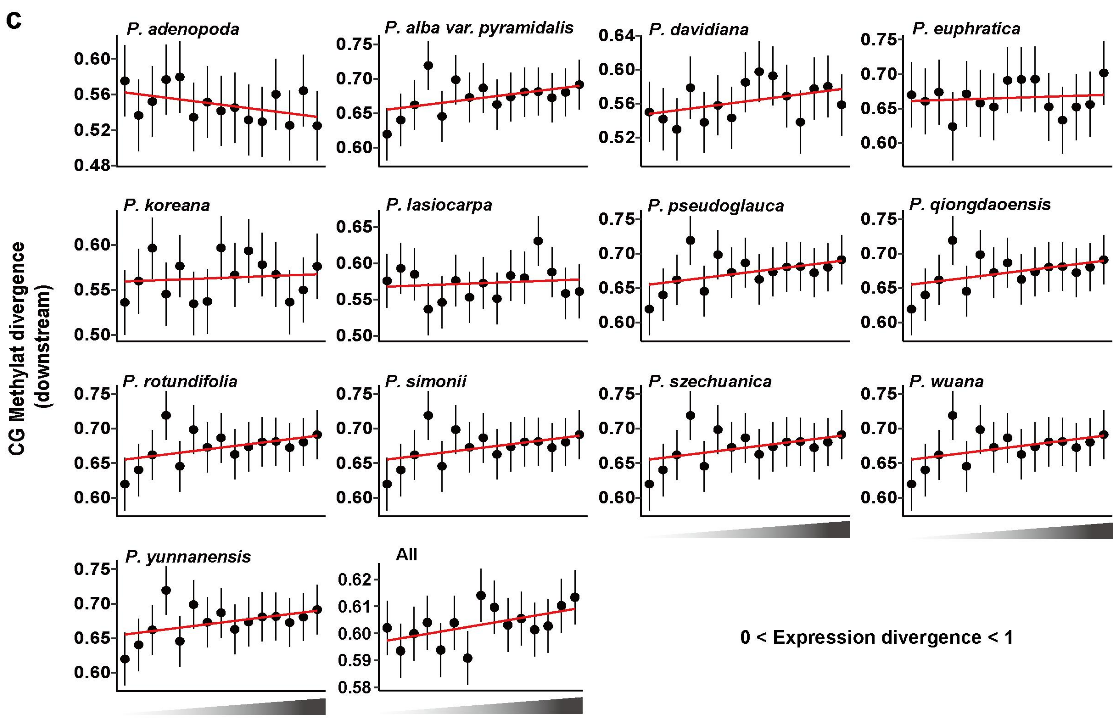


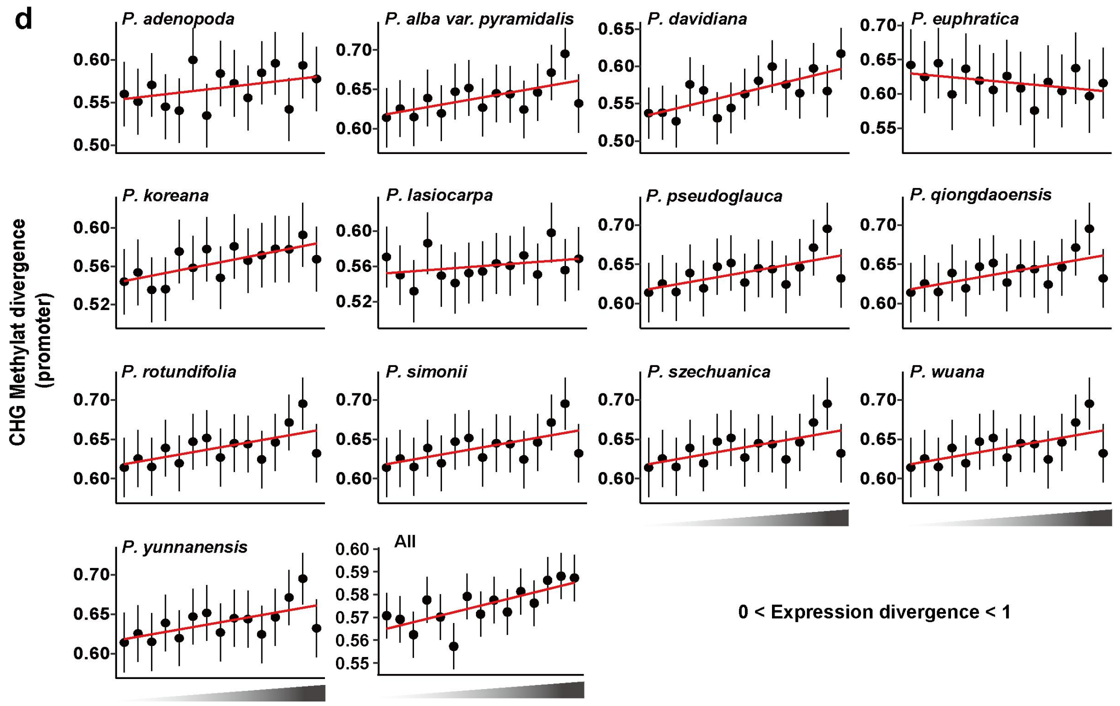


**Supplementary Fig. 29.** Continued on next page.

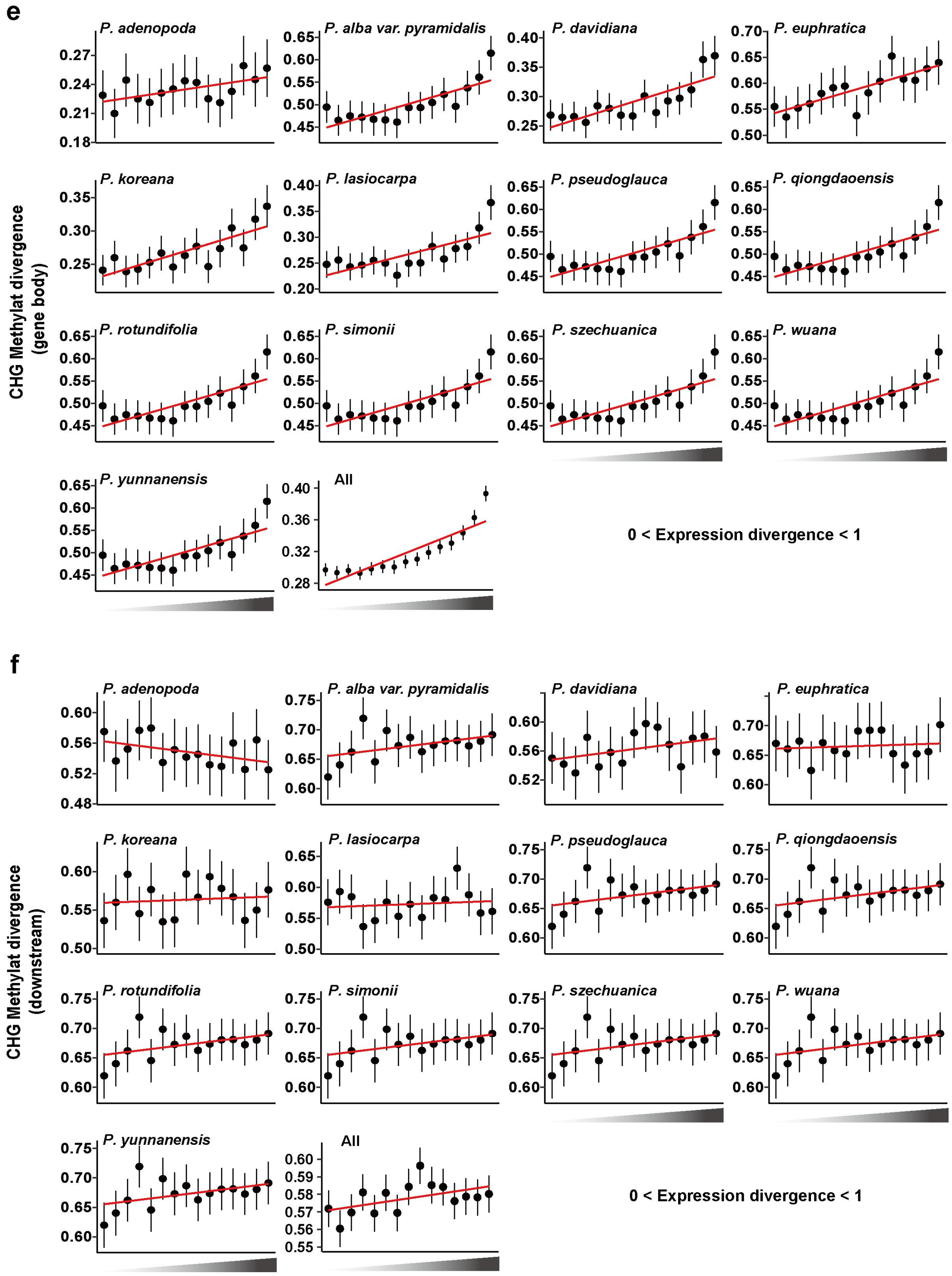


**Supplementary Fig. 29.** **Continued on next page.**

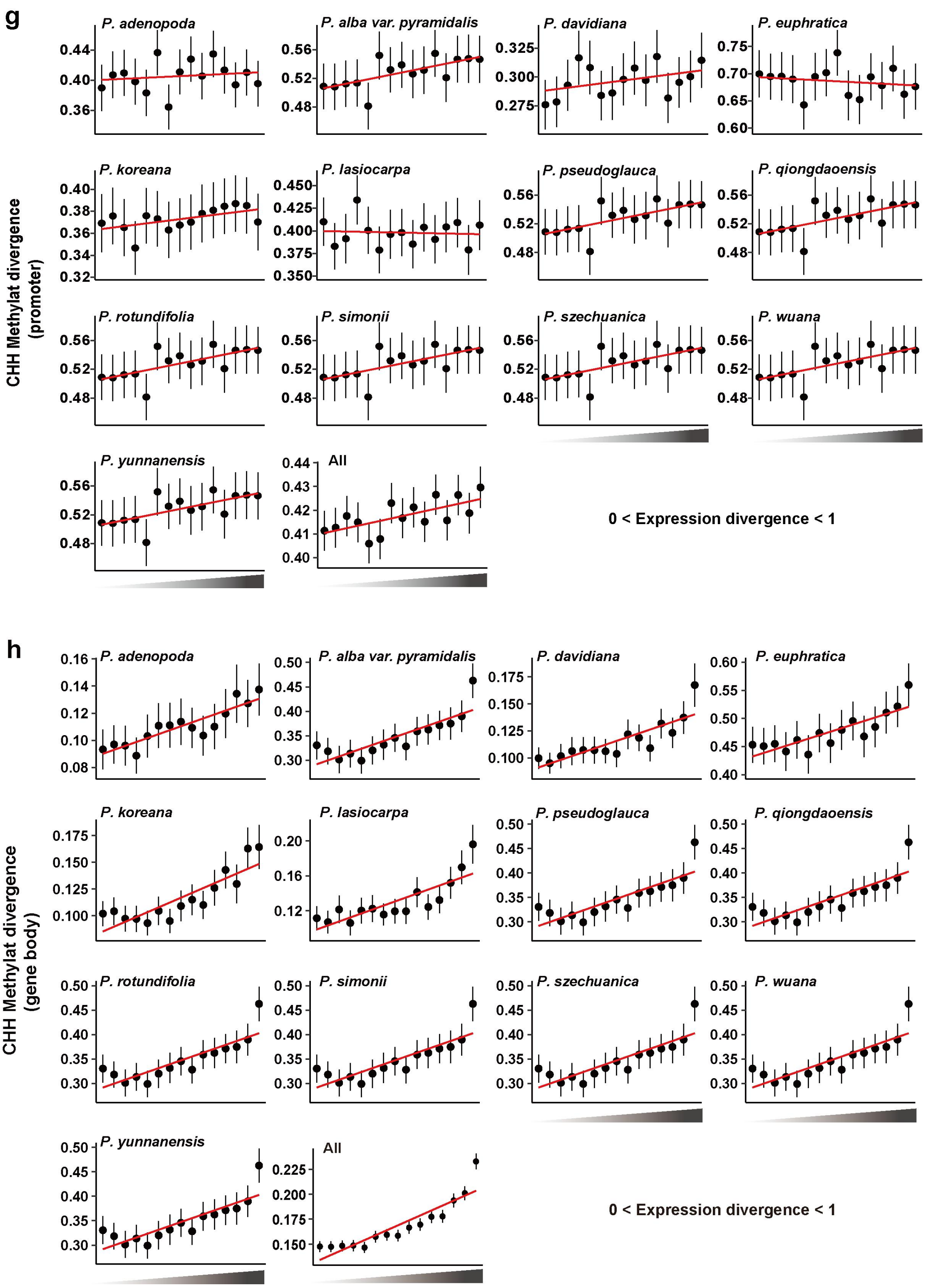


**Supplementary Fig. 29. Continued on next page.**

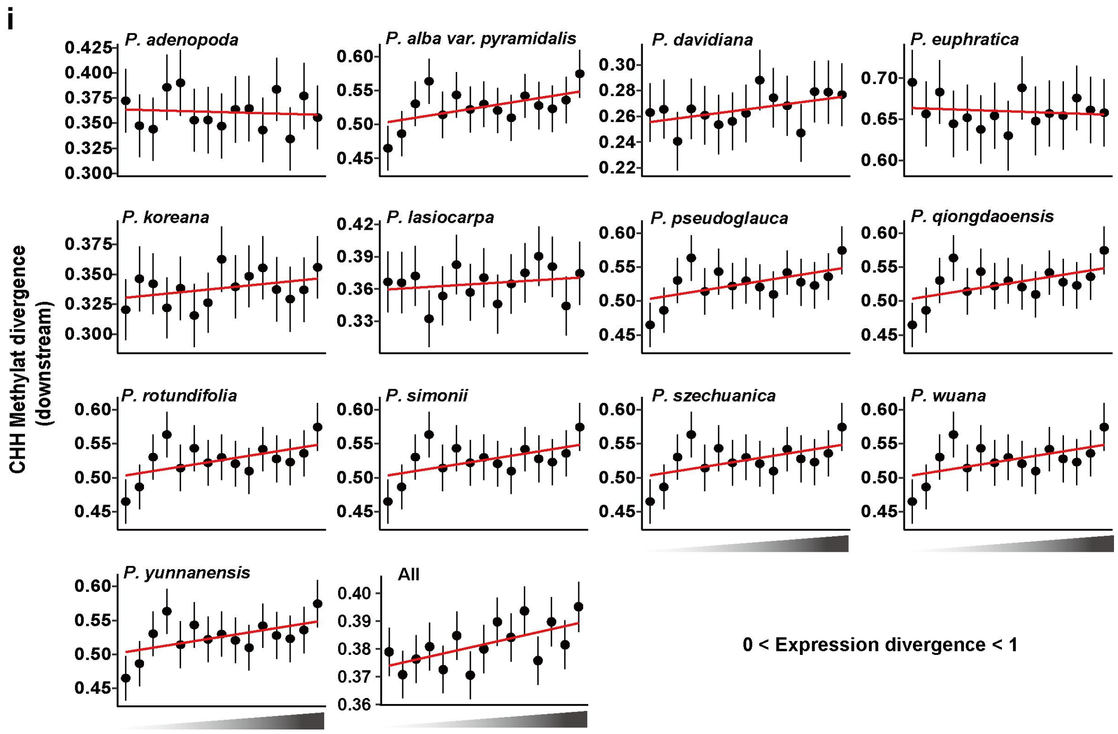


**Supplementary Fig. 29. Correlation between the expression divergence and methylation divergence of WGD-derived duplicated genes at CG (a-c), CHG (d-f), and CHH (g-i) contexts.**

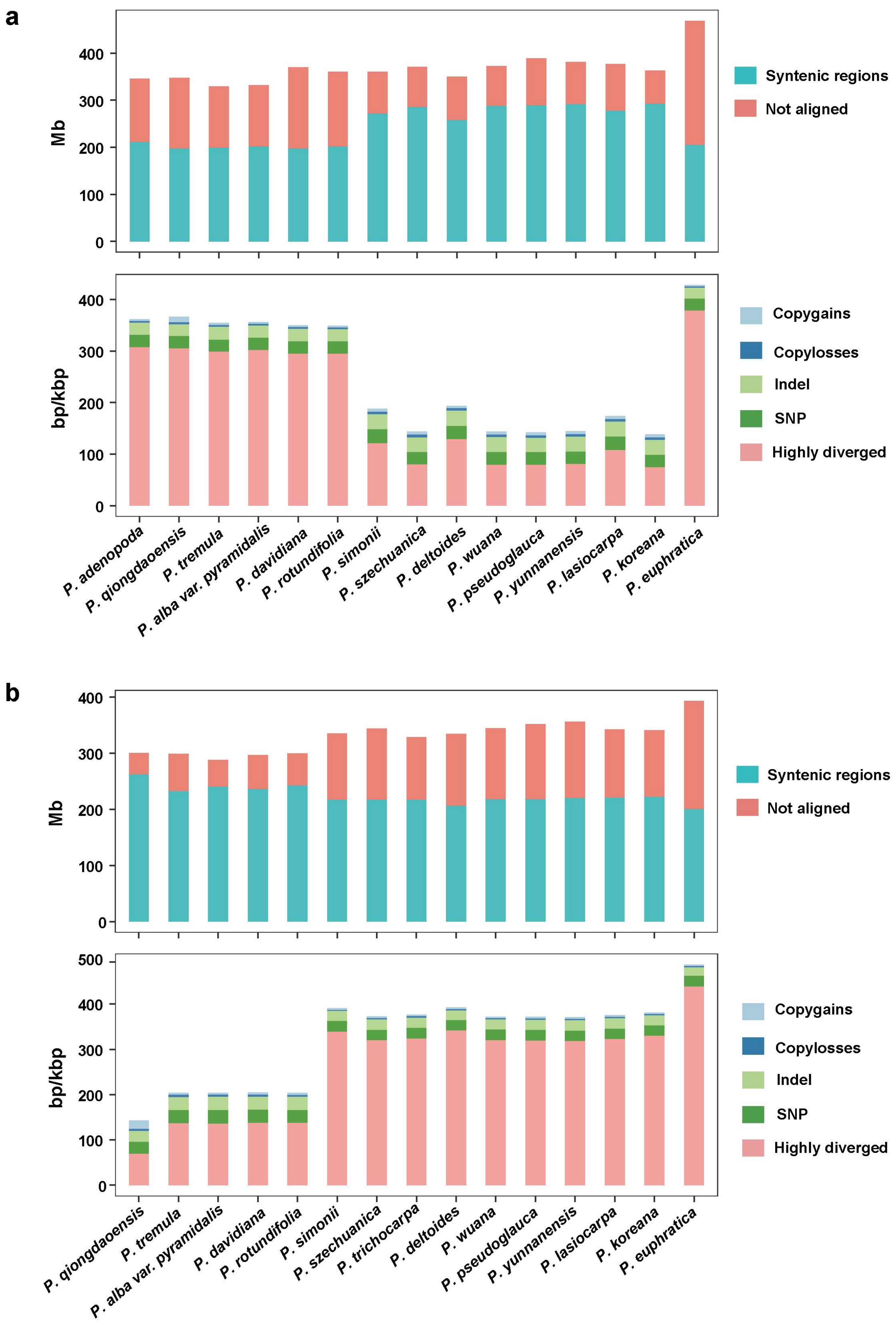


**Supplementary Fig. 30. Schematic of sequence variation detected using *P. trichocarpa* (a) and *P. adenopoda* (b) as reference genome, respectively.**

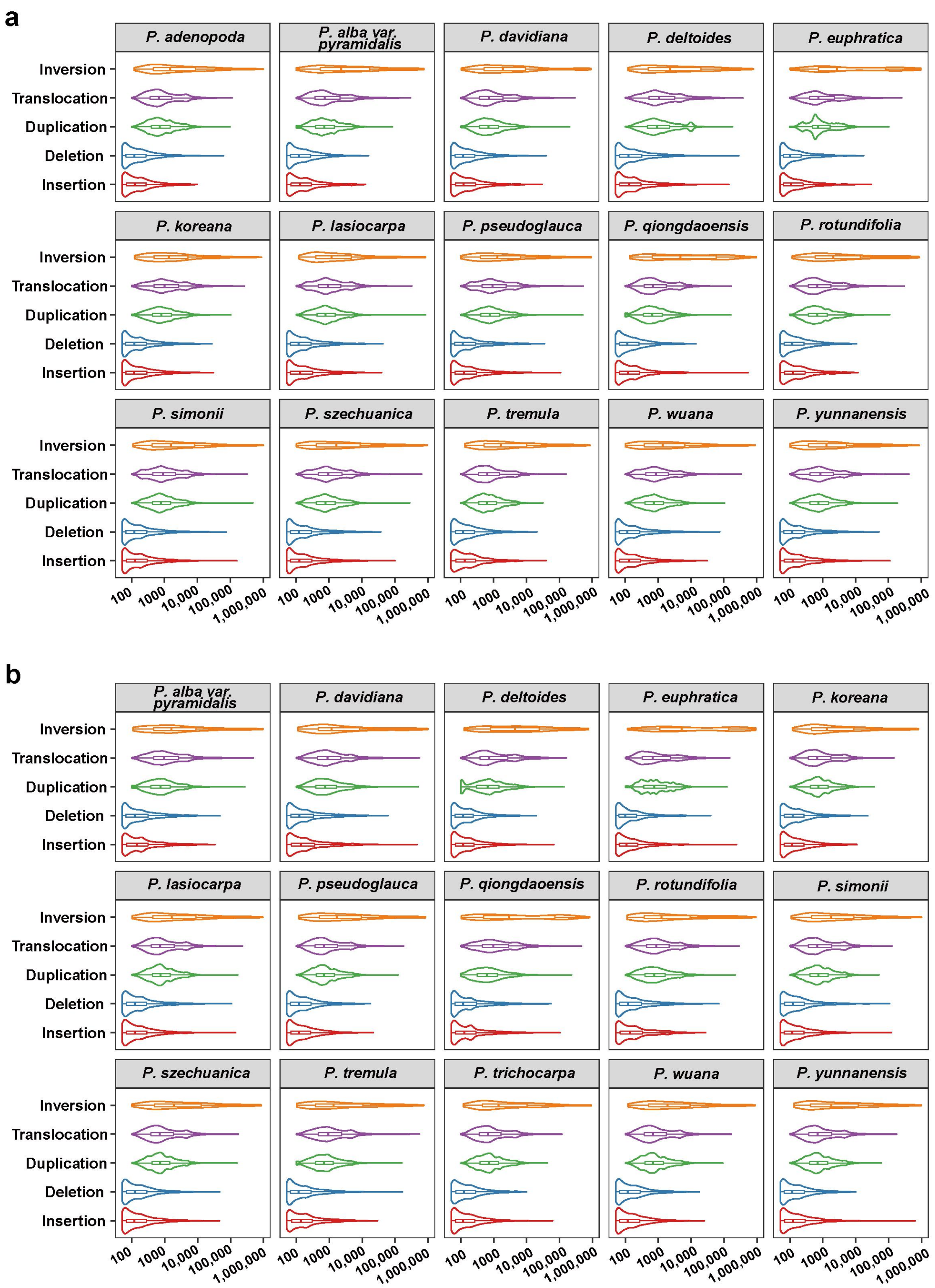


**Supplementary Fig. 31. Size distributions of different types of structural variation detected using *P. trichocarpa* (a) and *P. adenopoda* (b) as reference genome, respectively.**

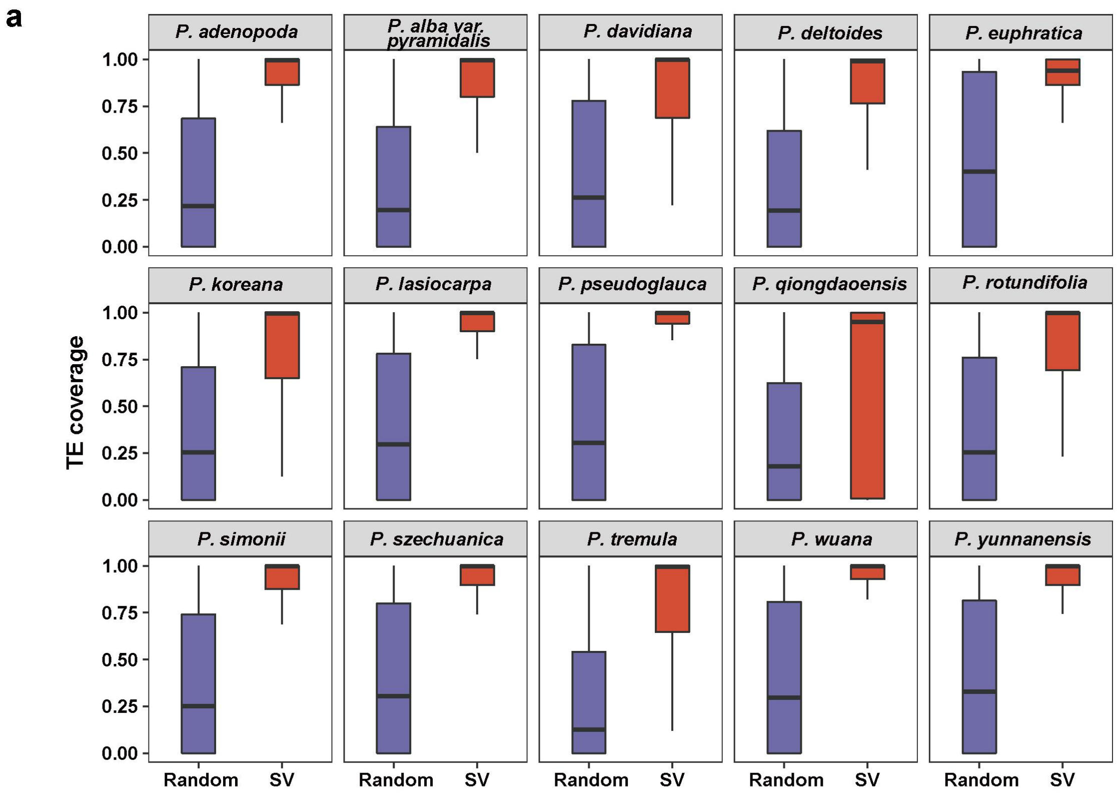


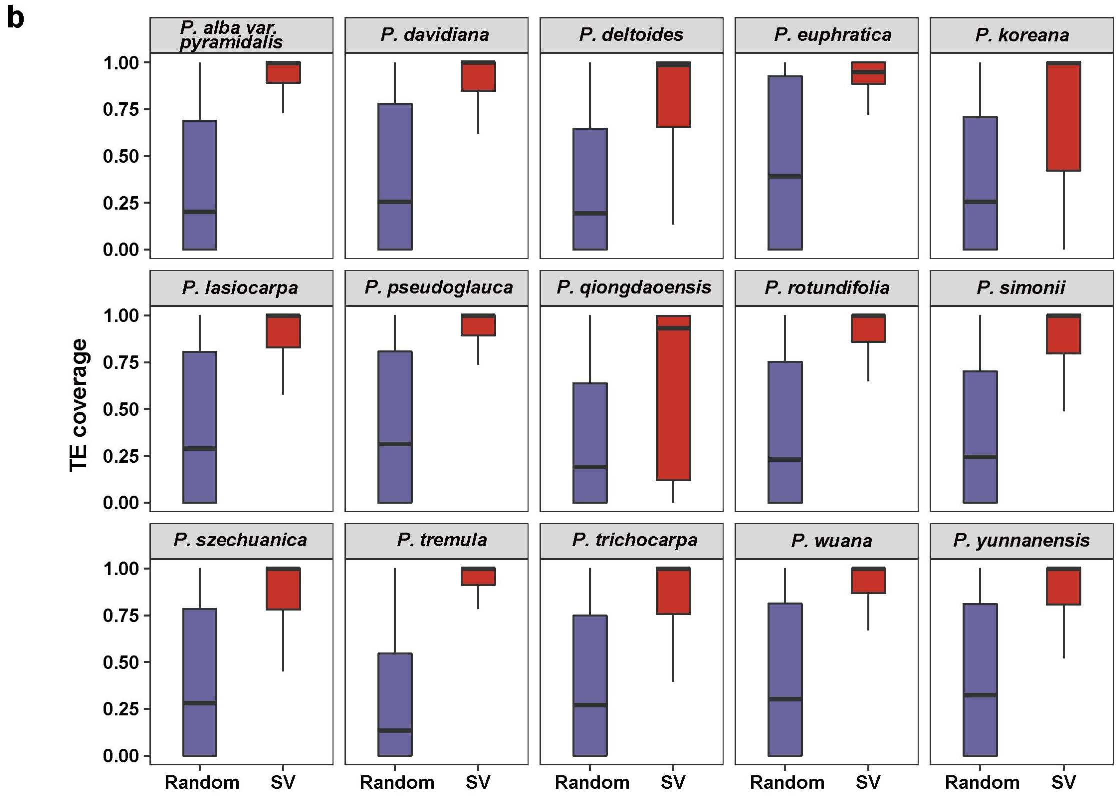


**Supplementary Fig. 32. Comparison of TE coverage between SV regions and genome regions selected randomly.** a, reference *P. trichocarpa.* b, reference *P.adenopoda.*

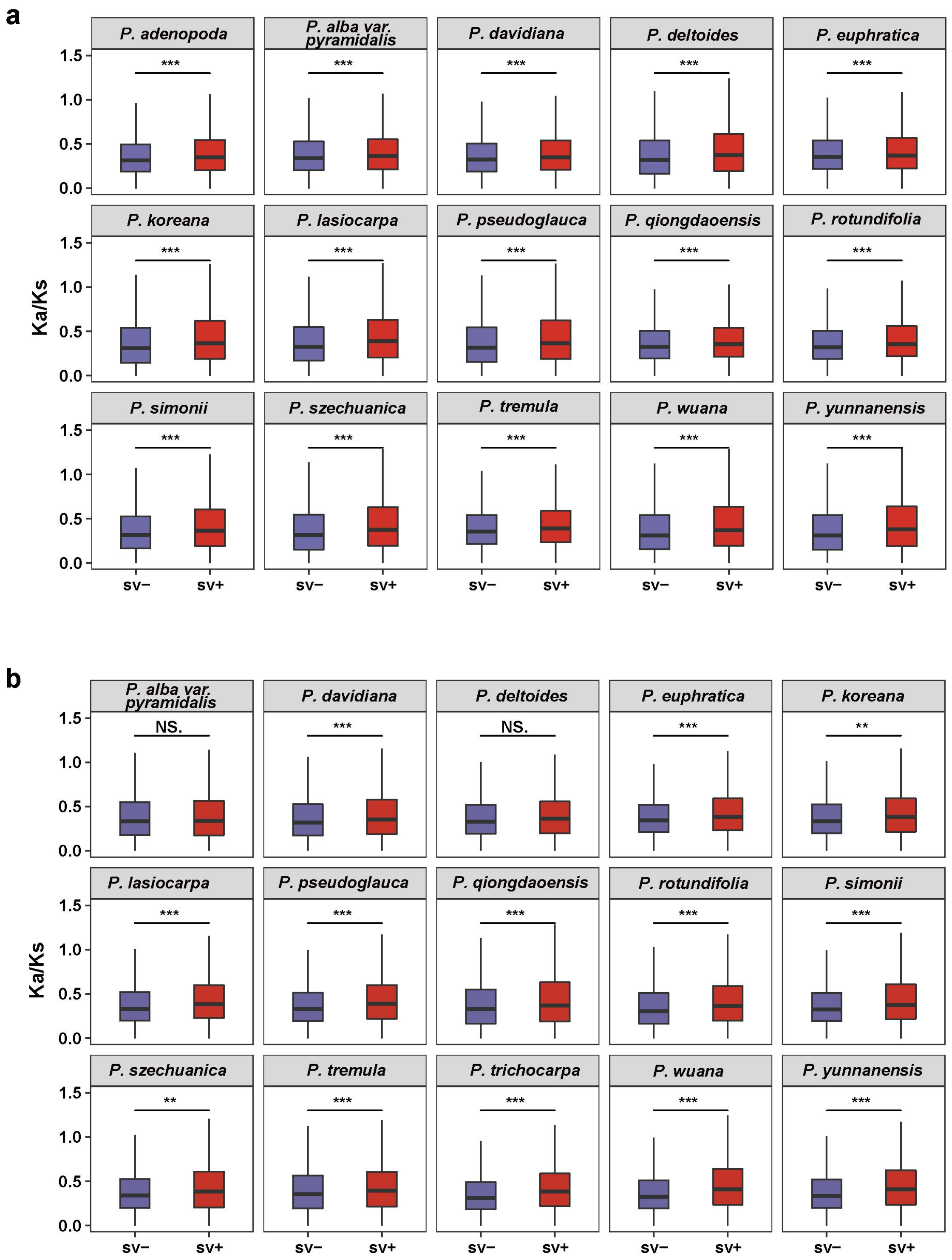


**Supplementary Fig. 33. Comparison of the selective constraint of genes with SVs and without SVs.** a, reference *P. trichocarpa.* b, reference *P.adenopoda*. Note, the *Ka/Ks* ratios were calculated relative to syntenic orthologs in reference genome.

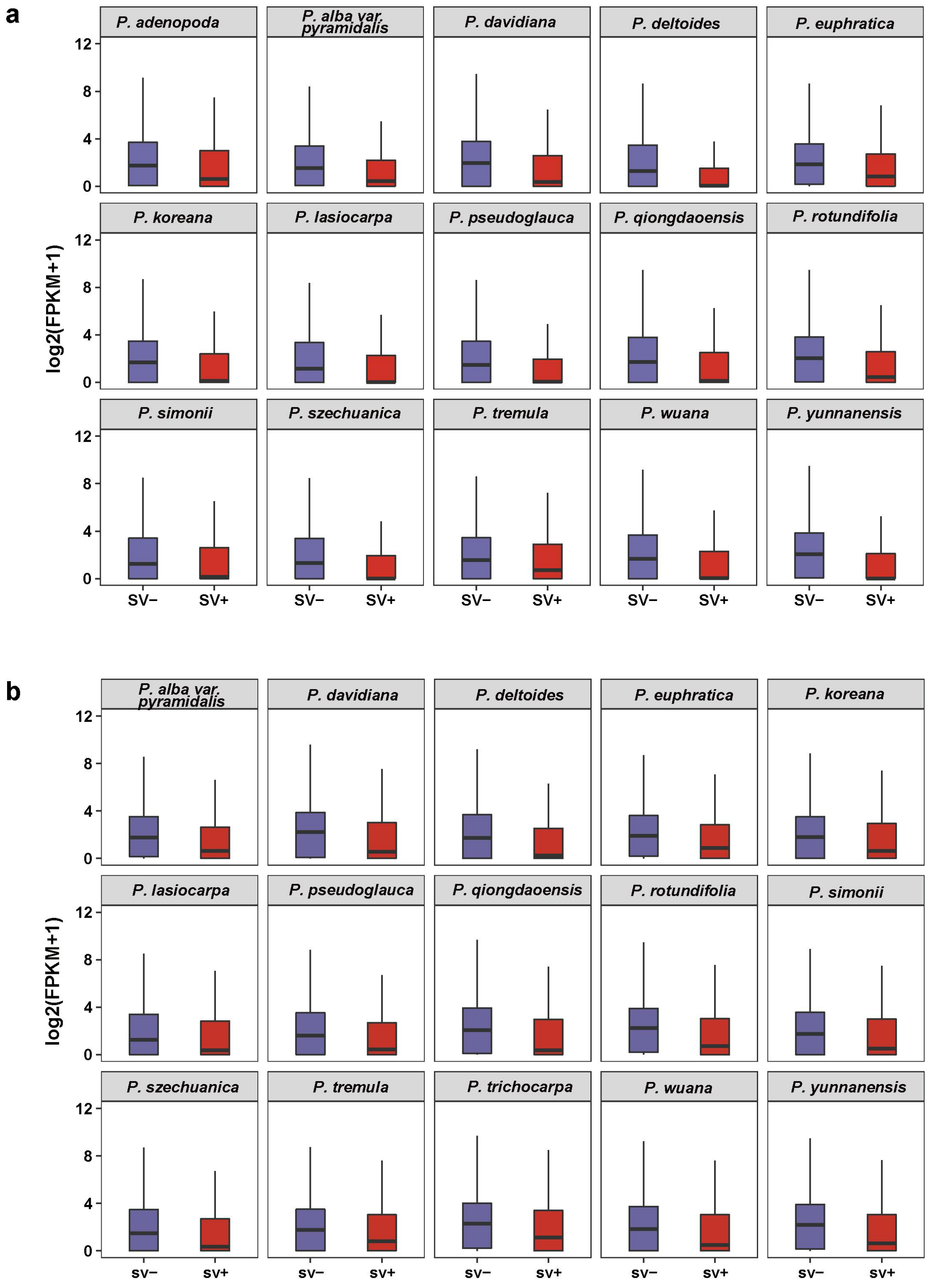


**Supplementary Fig. 34. Comparison of the expression levels (leaf tissue) of genes with SVs and without SVs.** a, reference *P. trichocarpa.* b, reference *P.adenopoda*.

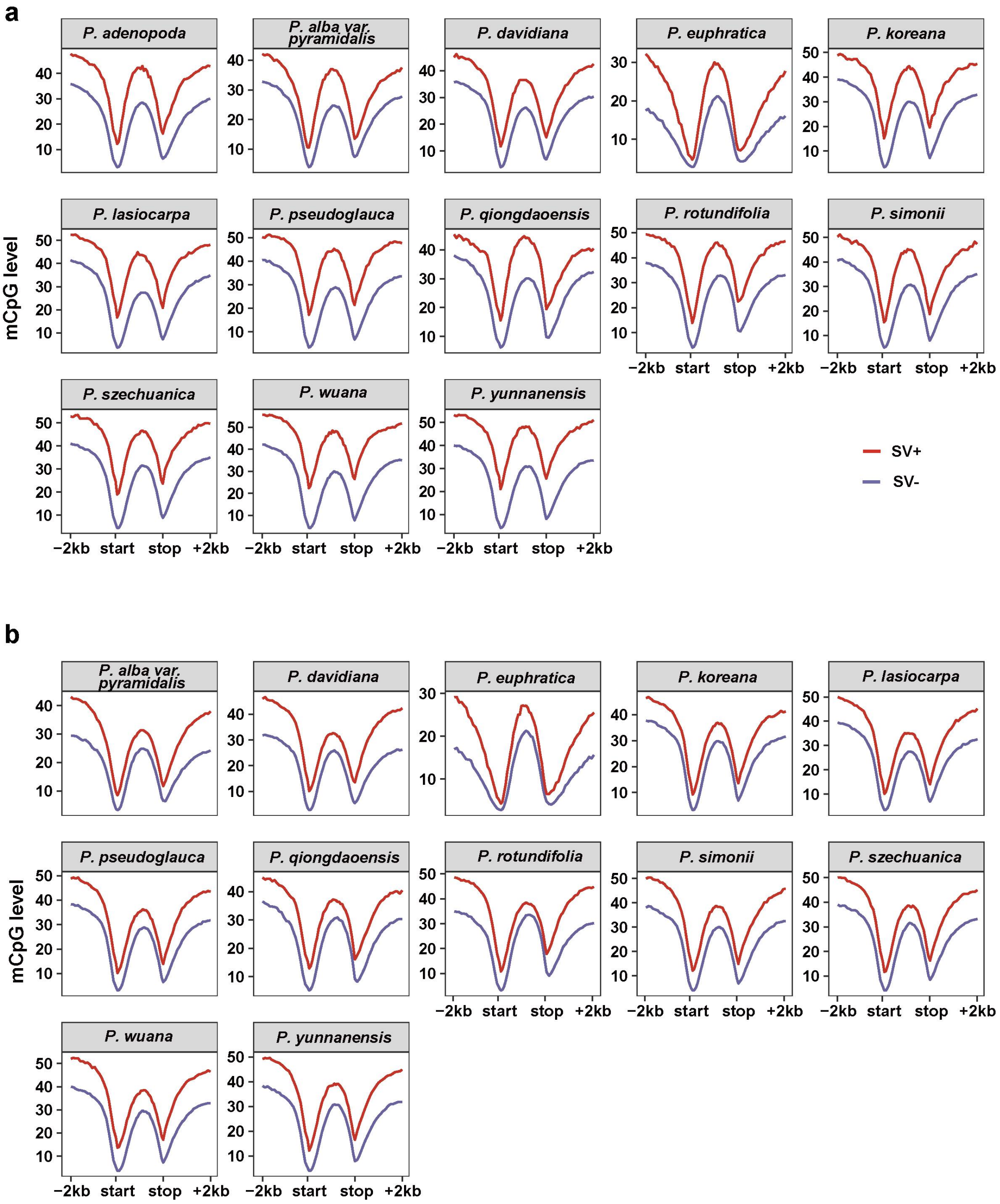


**Supplementary Fig. 35. Comparison of the methylation levels at CG context of genes with SVs and without SVs.** a, reference *P. trichocarpa.* b, reference *P.adenopoda*.

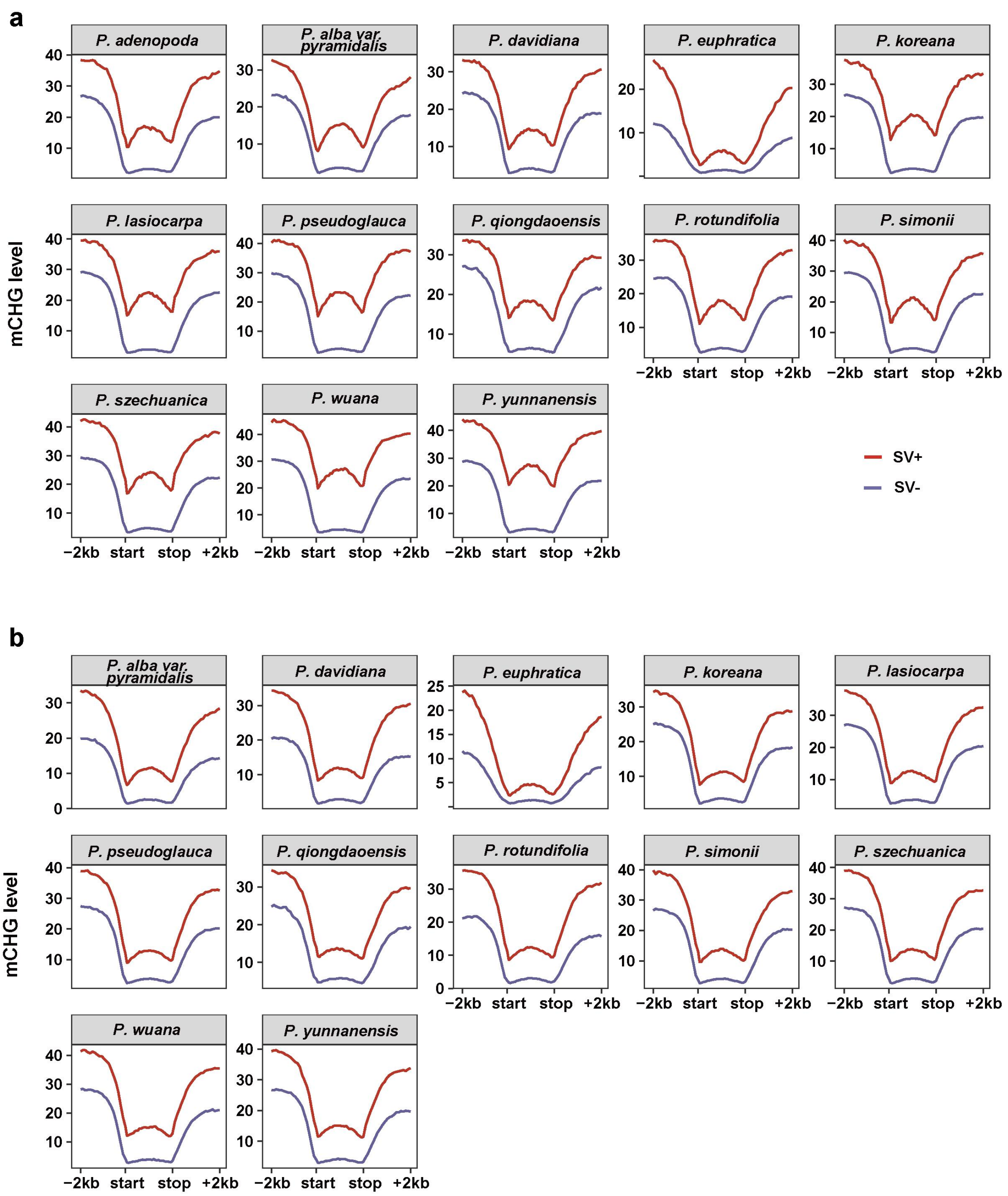


**Supplementary Fig. 36. Comparison of the methylation levels at CHG context of genes with SVs and without SVs.** a, reference *P. trichocarpa.* b, reference *P.adenopoda*.

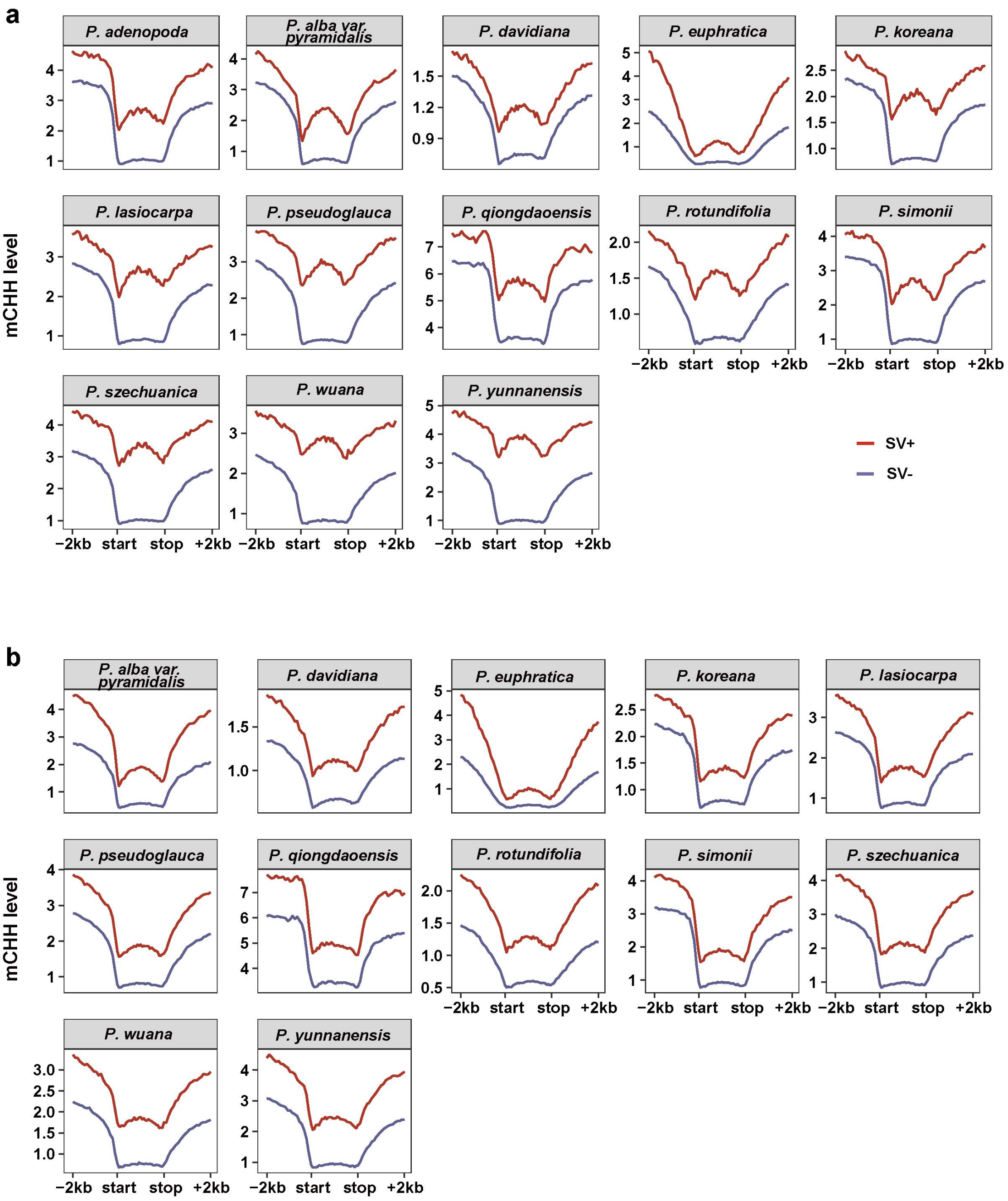


**Supplementary Fig. 37. Comparison of the methylation levels at CHH context of genes with SVs and without SVs.** a, reference *P. trichocarpa.* b, reference *P.adenopoda*.

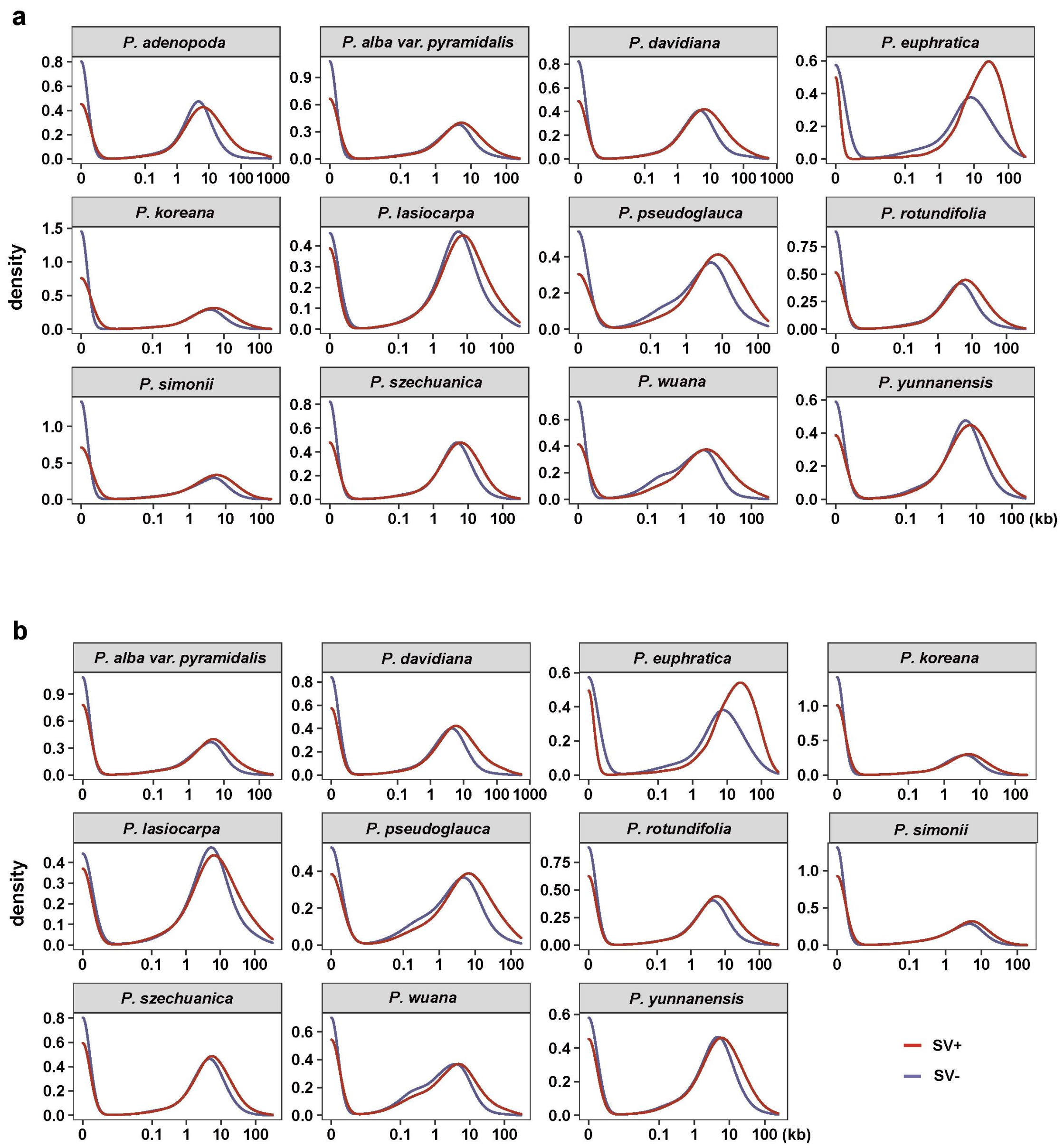


**Supplementary Fig. 38. Comparison of the accessible chromatin region (ACR) distribution of genes with SVs and without SVs.** a, reference *P. trichocarpa.* b, reference *P.adenopoda*.

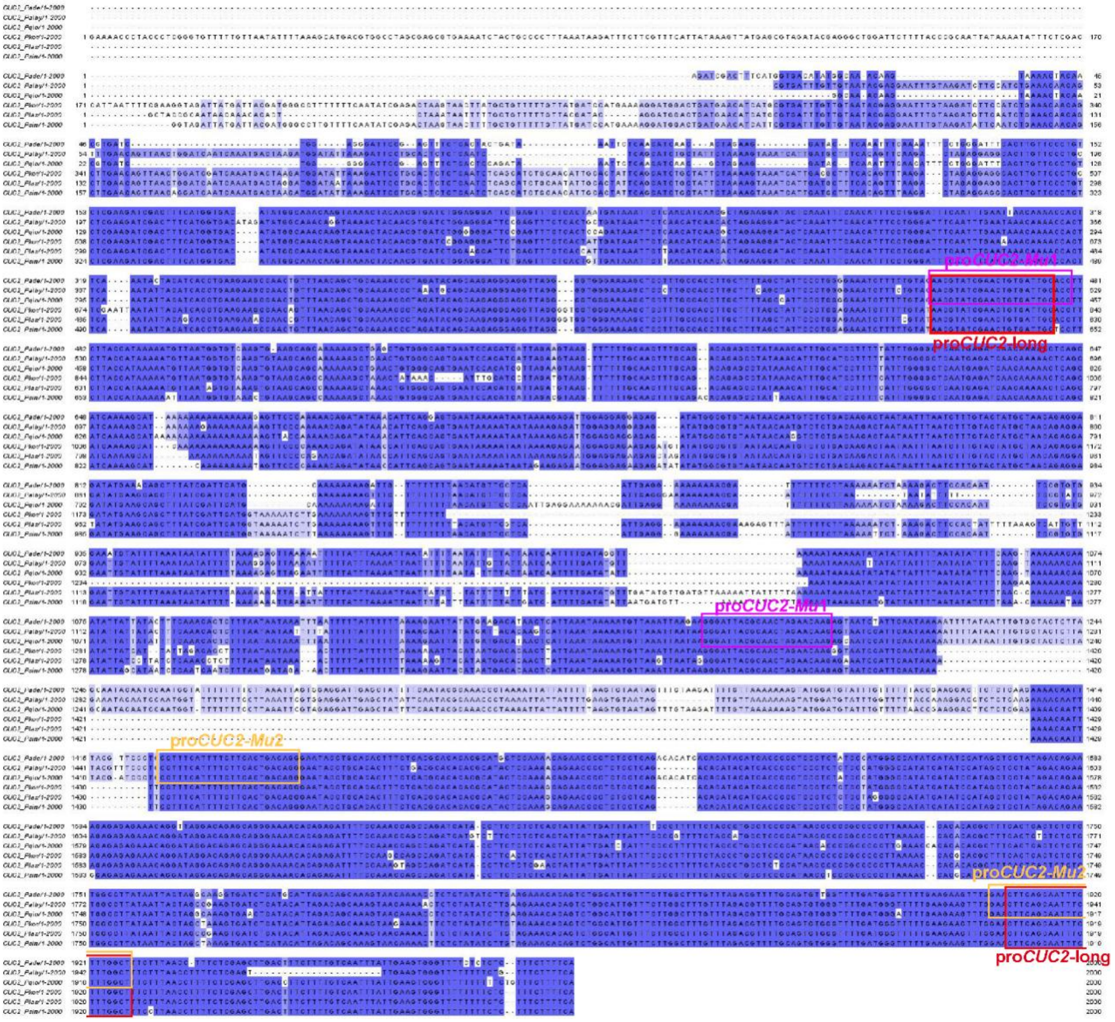


**Supplementary Fig. 39. The upstream (2 kb) promoter sequences of *CUC2* gene in different genomes.** Primers were designed based on the flanking sequence of the Indel. The target sites are represented by boxes of different colors.
